## Supplementary methods and figures for "Structural mimicry of UM171 and neomorphic cancer mutants co-opts E3 ligase KBTBD4 for HDAC1/2 recruitment"

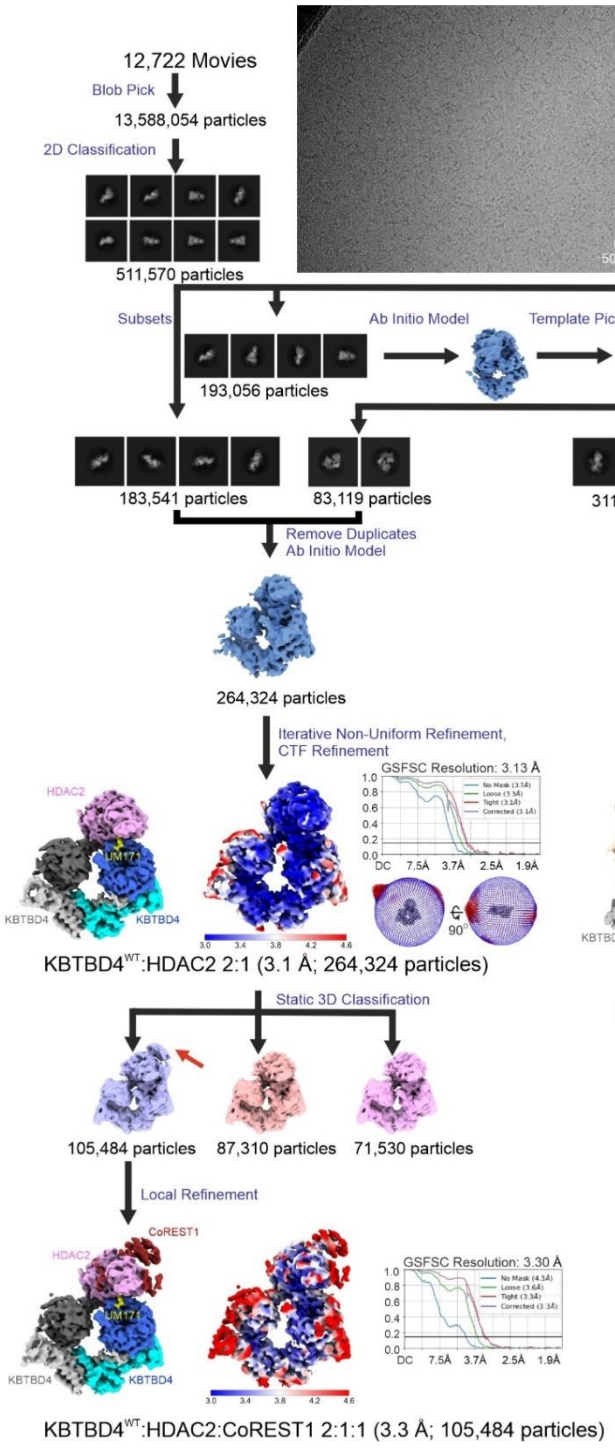


**Supplementary Figure 1. Schematic of pre-processing, classification and refinement procedures used to generate the cryo-EM maps for KBTBD4$^{WT\text{-}UM171}$-HDAC2 models.** The maps coloured by subunits are the final maps deposited and used for model building. Local resolutions of each map are shown coloured from highest resolution (dark blue) to lowest resolution (red). The nominal resolution of each map was calculated at FSC=0.143 cut-off as shown by the gold-standard Fourier shell correlation (GSFSC) graphs. The orientation distribution of the 2D images for each corresponding 3D model is shown in the Fourier 3D space.

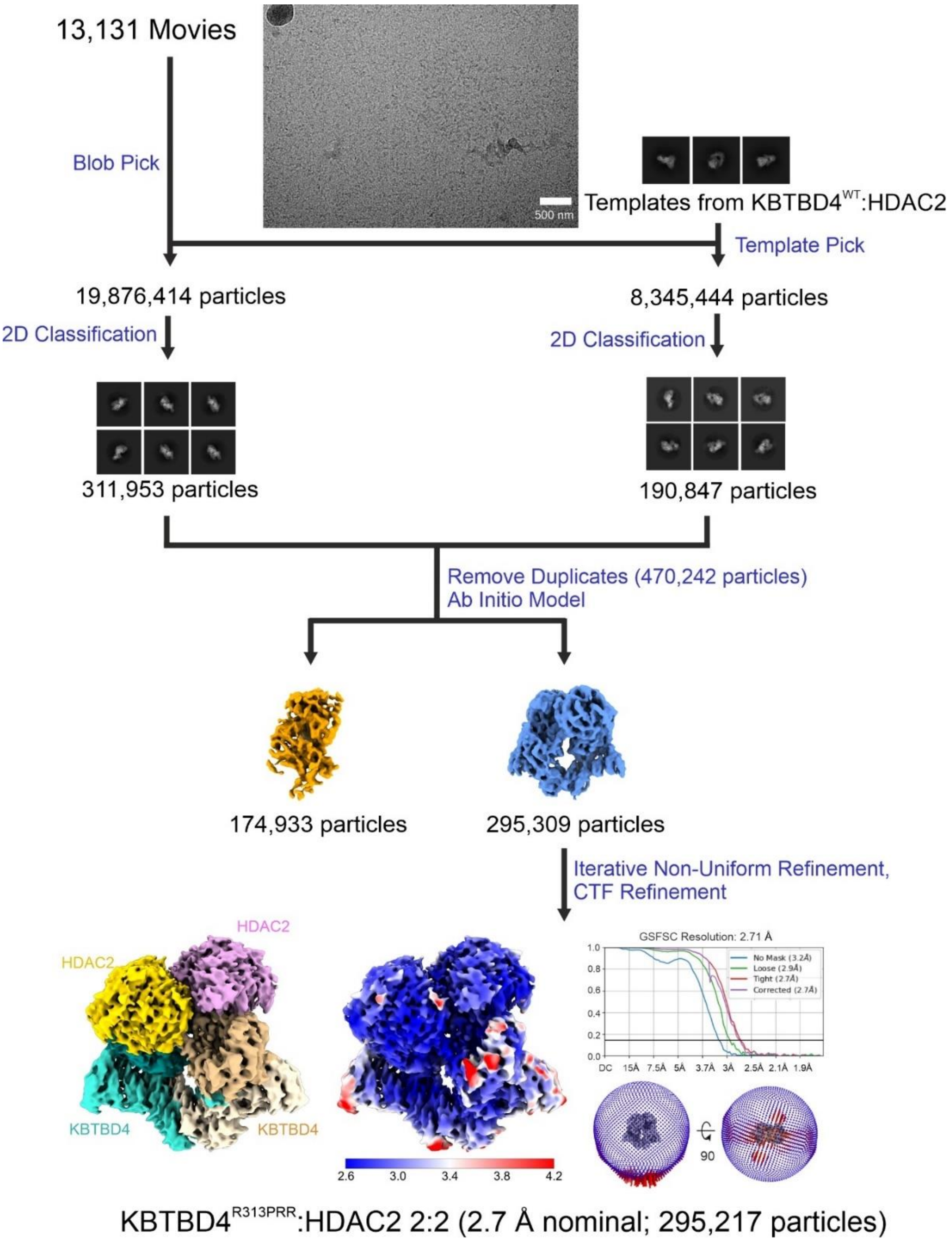


**Supplementary Figure 2. Schematic of pre-processing, classification and refinement procedures used to generate the cryo-EM map for KBTBD4$^{R313PRR}$-HDAC2 model.** The map coloured by subunits are the final map deposited and used for model building. Local resolutions of the map are shown coloured from highest resolution (dark blue) to lowest resolution (red). The nominal resolution of the map was calculated at FSC=0.143 cut-off as shown by the gold-standard Fourier shell correlation (GSFSC) graph. The orientation distribution of the 2D images is shown in the Fourier 3D space.

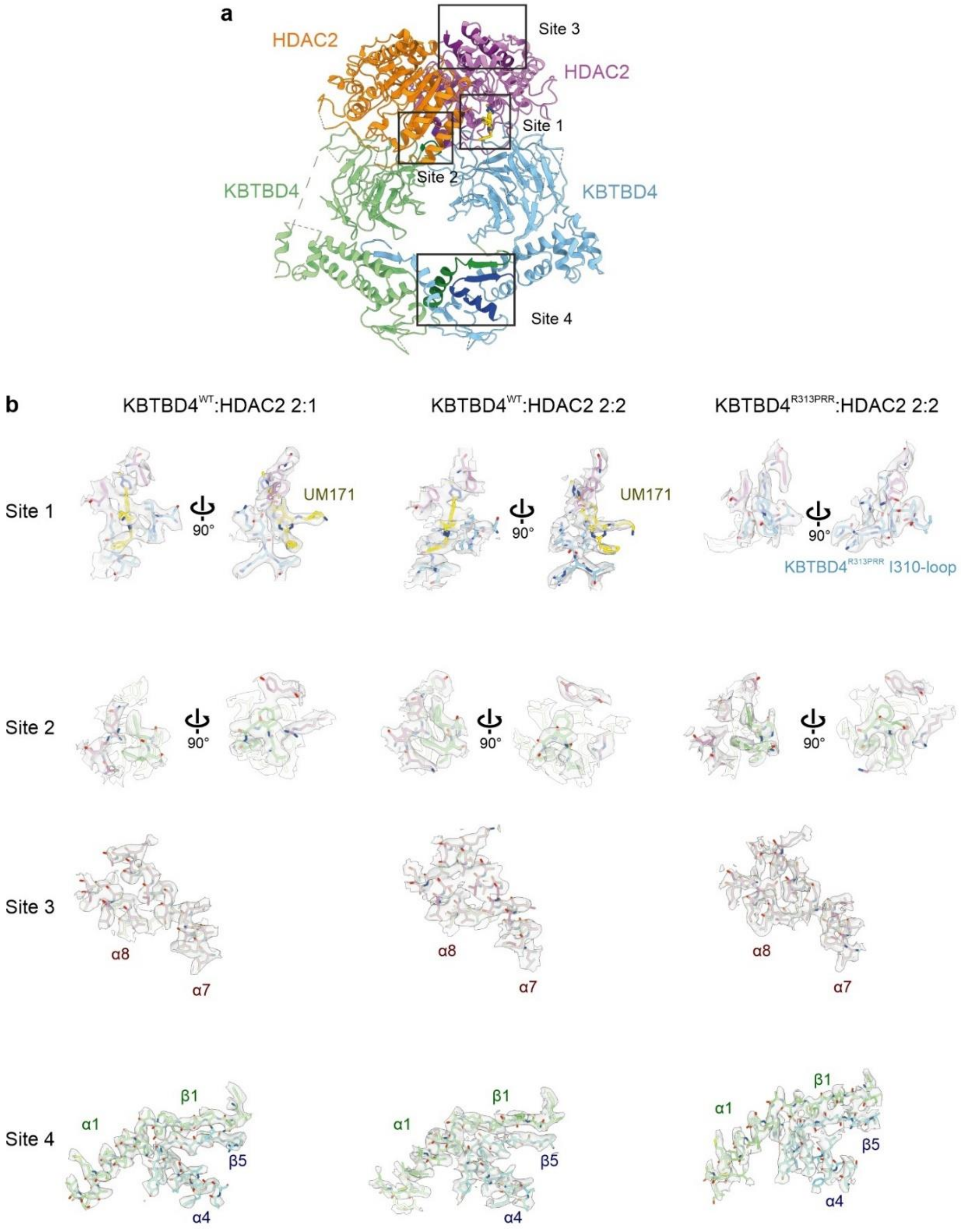


**Supplementary Figure 3. Analysis of the quality of the cryo-EM maps for all models. a,** Ribbon representation of the 2:2 KBTBD4$^{WT\text{-}UM171}$-HDAC2 model illustrating the locations of representative sites. HDAC2 chains are shown in purple and orange. KBTBD4 chains are shown in blue and green. UM171 is shown in yellow sticks. These sites as well as the equivalent sites in the 2:1 KBTBD4$^{WT\text{-}UM171}$-HDAC2 and 2:2 KBTBD4$^{R313PRR}$-HDAC2 models were analyzed as shown in **b**. **b,** Cryo-EM densities and the fitted atomic models are shown for all models as indicated. The colour scheme is the same as in **a**.

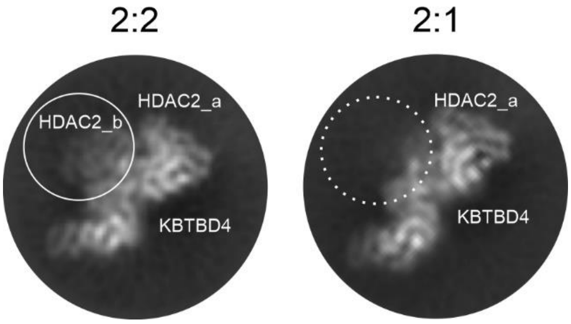


**Supplementary Figure 4. Example 2D classes for 2:2 and 2:1 KBTBD4$^{R313PRR}$-HDAC2 complexes with a similar view.** HDAC2_a and the KBTBD4 dimer are labelled in both 2:2 and 2:1 2D class images to indicate the similar orientation. HDAC2_b subunit is circled and labelled in the 2:2 image, while a dashed-line circle indicates the absence of it in the 2:1 image.

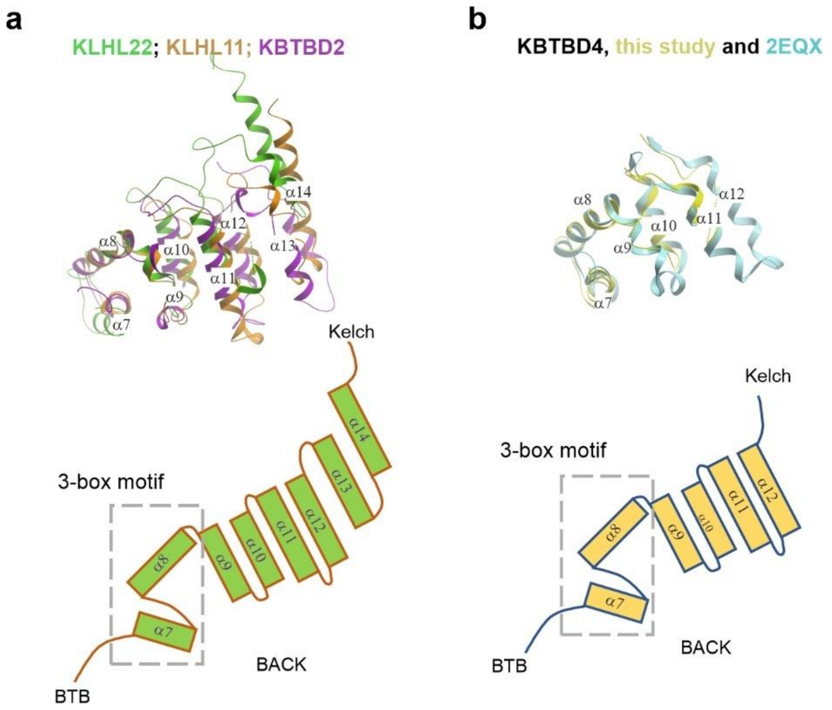


**Supplementary Figure 5. Superposition of previously determined BACK domain structures. a**, Superposition of the canonical BACK domains in KLHL22 (PDB 8KHP, green), KBTBD2 (PDB 8GQ6, purple) and KLHL11 (PDB 4APF, clay) comprise 8 helices. The topology diagram in green illustrates their arrangement, the CUL3-binding 3-box motif, as well as the loops connecting the N-terminal BTB and C-terminal Kelch domains. **b**, KBTBD4 structures from this study (PDB 9GGL, yellow) and previous NMR work (PDB 2EQX, cyan) show a smaller BACK domain of six helices. The topology diagram in yellow illustrates their arrangement as shown in **a**.

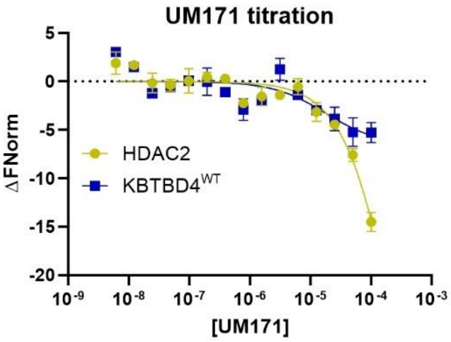


**Supplementary Figure 6. MST assay showing UM171 titration into KBTBD4$^{WT}$ or HDAC2 alone.** Recombinant KBTBD4$^{WT}$ or HDAC2 was purified and labelled with fluorescent dye. Different concentrations of UM171 were titrated into 40 nM labelled KBTBD4$^{WT}$ or 80 nM labelled HDAC2 for MST measurements.

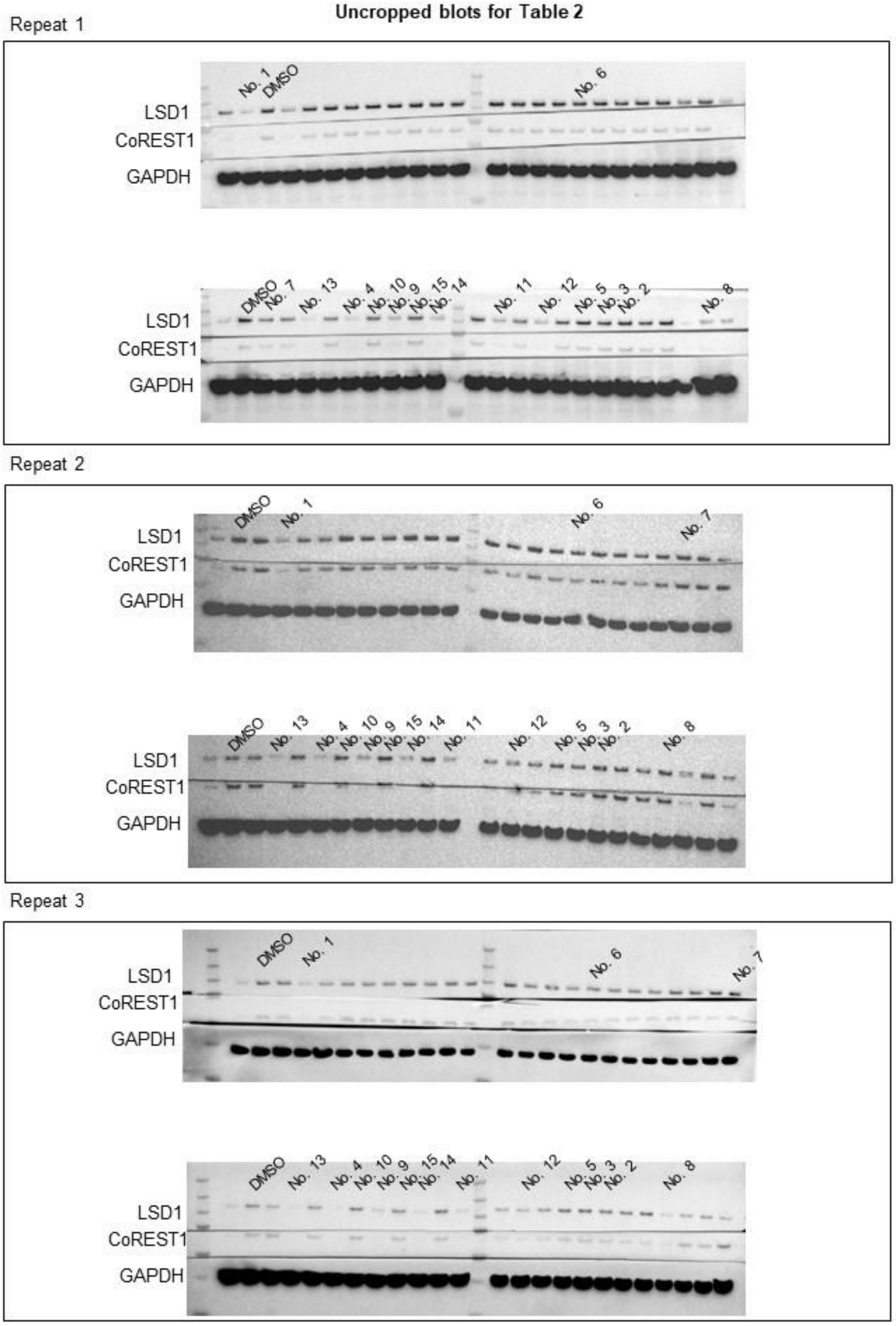


**Supplementary Figure 7. Uncropped Immunoblots for Table 2 showing the protein levels of CoREST1 and LSD1 in HEK293 cells treated with UM171 and analogs**. The cellular degradation assay was performed in triplicates. LSD1 and CoREST1 were detected by corresponding antibodies. GAPDH1 was probed as a loading control for protein abundance normalization. The lanes corresponding to compounds 1-15 are labelled as shown.

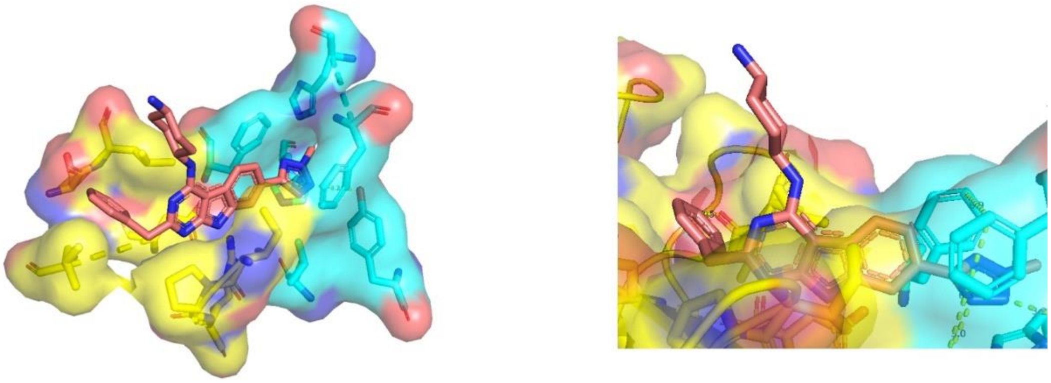

**Supplementary Figure 8. UM171 buries in the protein interface explaining the steep SAR observed in the cellular assay.** KBTBD4$^{WT}$ and HDAC2 are shown in surface representation while UM171 and the key interacting protein residues are shown in sticks. KBTBD4$^{WT}$ is coloured in yellow, HDAC2 in cyan and UM171 in pink.

### Compound synthesis report

All reagents obtained from commercial sources were used without further purification. Anhydrous solvents were obtained from commercial sources and used without further drying. The reactions were monitored using LC-MS and GC-MS instruments.

Analytical LC-MS: Agilent HP1200 LC with Agilent 6140 quadrupole MS, operating in positive or negative ion electrospray ionisation mode. The molecular weight scan range was 100 to 1350 m/z. Parallel UV detection was done at 210 nm and 254 nm. Samples were supplied as a 1 mM solution in MeCN or in THF/water (1:1) with 5 µL loop injection. LCMS analyses were performed on two instruments, one of which was operated with basic, and the other with acidic eluents.

Basic LC-MS: Gemini-NX, 3 µm, C18, 50 mm × 3.00 mm i.d. column at 23°C, at a flow rate of 1 mL min-1 using 5 mM aq. NH4HCO3 solution and MeCN as eluents.

Acidic LC-MS: ZORBAX Eclipse XDB-C18, 1.8 µm, 50 mm × 4.6 mm i.d. column at 40°C, at a flow rate of 1 mL min-1 using water and MeCN as eluents, both containing 0.02 V/V% formic acid.

Combination gas chromatography and low-resolution mass spectrometry were performed on Agilent 6850 gas chromatograph and Agilent 5975C mass spectrometer using 15 m × 0.25 mm column with 0.25 µm HP-5MS coating and helium as carrier gas. Ion source: EI+, 70 eV, 230°C, quadrupole: 150°C, interface: 300°C.

Flash chromatography was performed on ISCO CombiFlash Rf 200i with pre-packed silica-gel cartridges (RediSep®Rf Gold High Performance).

Preparative HPLC purifications were performed on an Armen Spot Liquid Chromatography system with a Gemini-NX® 10 µm C18, 250 mm × 50 mm i.d. column running at a flow rate of 118 mL min-1 with UV diode array detection (210 – 400 nm).

1H NMR and proton-decoupled 13C NMR measurements were performed on Bruker Avance III 500 MHz spectrometer and Bruker Avance III 400 MHz spectrometer, using DMSO-d6 or CDCl3 as solvent. 1H and 13C NMR data are in the form of delta values, given in part per million (ppm), using the residual peak of the solvent as internal standard (DMSO-d6: 2.50 ppm (1H) / 39.5 ppm (13C); CDCl3: 7.26 ppm (1H) / 77.0 ppm (13C)). Splitting patterns are designated as: s (singlet), d (doublet), t (triplet), q (quartet), sp (septet), m (multiplet), br s (broad singlet), dd (doublet of doublets), td (triplet of doublets), qd (quartet of doublets). In some cases, two sets of signals appear in the spectra due to hindered rotation.

HRMS were determined on a Shimadzu IT-TOF, ion source temperature 200°C, ESI +/-, ionization voltage: (+-)4.5 kV. Mass resolution min. 10000.

All obtained products had an LC purity above 96% that was corroborated by their 1H NMR spectrum unless specifically mentioned otherwise.

**General Procedure A** – *Nitro reduction (partial)*

To the suspension of the appropriate nitro derivative (1.0 eq.) and the $NaBH_4$ (2.0 eq.) in EtOH (2.4 mL/mmol) and $H_2O$ (1.2 mL/mmol) was added dropwise $Pd(OAc)_2$ (0.2 mol%) in DCM (0.1 mL/mmol) at 0 °C under inert atmosphere. Once the conversion was full by LC-MS, the reaction was quenched by water, extracted with DCM (3x), dried, concentrated, and purified by either recrystallization from *n*-heptane:EtOAc or by normal-phase column chromatography using *n*-heptane and EtOAc eluents to give the desired product.

**General Procedure B1** – *Acylation I*

The solution of the appropriate hydroxylamine derivative (1.0 eq.) and TEA (3.0 eq.) in THF (2.0 ml/mmol) was cooled to 0 °C, the acylating agent (acid chloride or acid anhydride, 1.1-1.5 eq.) was added dropwise. The mixture was stirred for 15-30 minutes on 0 °C. The reaction was monitored by LC-MS. In case the diacylated product appeared LiOH (1.6 eq.) in water (0.5 mL/mmol) and MeOH (0.5 mL/mmol) was added and stirred at 45 °C for 45 minutes to hydrolyze the O-Ac. After full conversion the volatiles were evaporated. If adding LiOH was necessary, the residue was acidified using 1M HCl solution and it was extracted with EtOAc (3x), washed with brine (2x), dried, concentrated and purified by normal-phase column chromatography using DCM and MeOH eluents to give the desired product.

**General Procedure B2** – *Acylation II*

To the solution of the appropriate hydroxylamine derivative (1.0 eq.) in $Et_2O$ (5.7 mL/mmol) $NaHCO_3$ (1.2 eq.) was added, the mixture was cooled to 0 °C under inert atmosphere, and acetyl chloride (1.2 eq.) in $Et_2O$ (3.8 mL/mmol) was added dropwise. The reaction mixture was stirred at 0 °C for 3 hours. The reaction was monitored by LC-MS. The mixture was filtered and extracted with EtOAc (3x), dried, concentrated and purified by normal-phase column chromatography using DCM and MeOH as eluent. The desired product was recrystallized from heptane-EtOAc.

**General Procedure C1** – *Indole synthesis I*

The solution of the appropriate acylated derivative (1.0 eq.) in DCM (7.5 mL/mmol) was cooled to 0 °C, propanedinitrile (1.0 eq.) and TEA (1.1 eq.) were added, and the mixture was allowed to warm up and it was stirred for 1 hour. After concentration, the residue was diluted in MeOH (7.5 mL/mmol) and NaOMe (1.0 eq.) was addedn and the mixture was refluxed for 45 minutes. The reaction was monitored by LC-MS, the 1:1 mixture of two regioisomers were present. Upon cooling the desired isomer precipitated and was collected by filtration, dried in vacuum, and was used without further purification. The residue with the other regioisomer was purified using reversed-phased column chromatography, using 5mM NH4HCO3 solution and MeCN as eluent.

**General Procedure C2** – *Indole synthesis II*

To the solution of the appropriate acylated derivative (1.0 eq.) in DMSO (2.1 mL/mmol), malonitrile (2.0 eq.), CuI (0.1 eq.), L-proline (0.2 eq.) and $K_2CO_3$ (2.0 eq.) were added under inert atmosphere, and the mixture was stirred at 60 °C for 16 hours. The reaction was monitored by LC-MS. The mixture was cooled to RT, then water was added to precipitate the desired product and it was collected by filtration, washed with cold water, heptane and DIPE. The precipitate was dried in vacuum to give the desired product.

**General Procedure D1** – *Pyrimidinone ring closure I*

A microwave tube was charged with the solution of the appropriate indole derivative (1.0 eq.) in AcOH (10.0 mL/mmol), triethylorthoformate (8.0 eq.) and TFA (5.0 eq.) were added to the mixture. The sealed tube was stirred at 120 °C for 1 hour in a microwave reactor. The reaction was monitored by LC-MS. The precipitate was collected by filtration, washed with ice-cold water, dried in vacuum to give the desired product, which was used without further purification.

**General Procedure D2** – *Pyrimidinone ring closure II*

To the solution of the appropriate indole derivative (1.0 eq.) in DMA (6.0 mL/mmol) TEA (3.0 eq.) was added, it was cooled to 0 °C and 2- phenylacetyl chloride (2.0 eq.) was added in small portions and it was stirred at 0 °C for 30 minutes. The reaction was monitored by LC-MS. The mixture was poured into ice-cold water, the precipitate was collected by filtration and was recrystallized from EtOAc to give the intermediate.

To the solution of amino-phenylacetyl indole derivative (1.0 eq.) in 2-propanol (12.0 mL/mmol) potassium tert-butoxide (6.0 eq.) was added and it was stirred at 65 °C for 2 hours. The reaction was monitored by LC-MS. The mixture was evaporated, the residue was taken up in water and the pH was adjusted to 6 using 6 N HCl solution. The precipitate was collected by filtration, dried in vacuum to give the desired product, which was used without further purification.

**General Procedure D3** – *Pyrimidinone ring closure III*

A microwave tube was charged with the solution of the appropriate indole derivative (1.0 eq.) in MeOH (3.5 mL/mmol), methyl 2-phenylacetate (2.2 eq.), NaOMe (8.7 eq.) were added, and the sealed tube was stirred at 140 °C for 1.5 hours in a microwave reactor. The reaction was monitored by LC-MS. The mixture was cooled to RT, water and AcOH were added, and the slurry mixture was stirred for 30 minutes. The precipitate was collected by filtration, washed with cold MeOH, dried in vacuum, purified by reversed-phase column chromatography using 5mM $NH_4HCO_3$ solution and MeCN as eluent to give the desired product.

**General Procedure E** – *Nitrile hydrolysis*

To the solution of the appropriate pyrimidoindole derivative (1.0 eq.) in the mixture of 2-methoxyethanol (8.0 mL/mmol) and water (0.8 mL/mmol) NaOH (15.0 eq.) was added, and it was refluxed for 5 days. The reaction was monitored by LC-MS. The mixture was acidified using 2M HCl solution, and the precipitate was collected by filtration, washed with water, dried in vacuum to give the desired product, which was used without further purification.

**General Procedure F** – *Esterification*

To the solution of the appropriate carboxylic acid derivative (1.0 eq.) in the mixture of toluene (10.0 mL/mmol) and MeOH (2.0 mL/mmol) TMS-diazomethane (5.0 eq., 2 M solution in hexane) was added dropwise and the slurry was stirred for 3 hours. The reaction was monitored by LC-MS. The remaining TMS-diazomethane was quenched with a few drops of AcOH, and the solution was evaporated, and the residue was dried in vacuum to give the desired product, which was used without further purification.

**General Procedure G** – *Hydroxyl- chloride exchange*

A microwave tube was charged with the appropriate pyrimidinone derivative and $POCl_3$ (8.0 mL/mmol). The sealed tube was stirred at 175 °C for 22-40 minutes in a microwave reactor. The reaction was monitored by LC-MS. The mixture was cooled and poured into ice, the pH was adjusted to 7 using 50% NaOH solution. The precipitate was collected by filtration, washed with water and dried in vacuum to give the desired product, which was used without further purification.

**General Procedure H1** – *$SN_{Ar}$ I*

A microwave tube was charged with the solution of the appropriate chloride derivative (1.0 eq.) in DMA (15.0 mL/mmol) and the appropriate amine derivative (2.2 eq.-3.5 eq.) and DIPEA (15.0 eq.) or DBU (15.0 eq.) were added. The sealed tube was stirred at 120°-140 °C for 30 minutes – 8 hours in a microwave reactor. The reaction was monitored by LC-MS. The mixture was cooled and purified by reversed-phase column chromatography using 5mM $NH_4HCO_3$ solution or 0.2% TFA solution and MeCN as eluent to give the desired product.

**General Procedure H2**– *$SN_{Ar}$ II*

To the solution of the appropriate pyrimidinone derivative (1.0 eq.) in MeCN (20.0 mL/mmol) DBU (3.0 eq.) was added, and the mixture was stirred for 15 minutes, then BOP (1.5 eq.) was added, and the mixture was stirred at 80 °C for 16 hours. The reaction was monitored by LC-MS. The mixture cooled and purified by reversed-phase column chromatography using 5mM $NH_4HCO_3$ solution or 0.2% TFA solution and MeCN as eluent to give the desired product.

**General Procedure H3**– *$SN_{Ar}$ III*

A microwave tube was charged with the solution of the appropriate pyrimidinone derivative (1.0 eq.) in the mixture of MeCN (12.0 mL/mmol) and DMSO (3.0 mL/mmol) and the appropriate amine derivative (2.0 eq.) and TEA (2.0 eq.) were added. The sealed tube was stirred at 140 °C for 3.5 hours in a microwave reactor. The reaction was monitored by LC-MS. The mixture was cooled, concentrated and the residue was poured into 10 mL water. The mixture was extracted with EtOAc (4x), dried, concentrated and purified by reversed-phase column chromatography using 5mM $NH_4HCO_3$ solution and MeCN as eluent to give the desired product.

**General Procedure I** – *Boc removal*

The appropriate Boc derivative (1.0 eq.) in dioxane (20.0 mL/mmol) was treated with HCl (20.0 eq.) and stirred for 4 hours. The reaction was monitored by LC-MS. The reaction mixture was concentrated, and the desired product was used without further purification.

**General Procedure J** – *Basic hydrolysis*

To the solution of the appropriate methyl ester (1.0 eq.) in the mixture of water (6 mL/mmol) and THF (25.0 mL/mmol), LiOH (2.0 eq.) was added, and it was stirred at 30 °C for 2 hours. The reaction was monitored by LC-MS. The mixture was concentrated, and the residue was acidified using 1M HCl. If there was precipitation the precipitate was collected by filtration and dried in vacuum. In other cases, the residue was purified by reversed-phase column chromatography using 5mM $NH_4HCO_3$ solution and MeCN as eluent to give the desired product.

**General Procedure K** – *Nitro reduction (full)*

To the solution of the appropriate nitro derivative (1.0 eq.) in the mixture of EtOH (8.2 mL/mmol) and water (8.2 mL/mmol), Fe (4.0 eq.) and $NH_4Cl$ (5.0 eq.) were added, and it was stirred at 60 °C for 6 hours. The reaction was monitored by LC-MS. The mixture was filtered through a celite pad and concnetrated. The residue was diluted with water, extracted with EtOAc (3x), washed with brine (3x), dried and concentrated to give the desired product, which was used without further purification.

**General Procedure L1** – *SN Ar IV*

A microwave tube was charged with the appropriate amine derivative (1.0 eq.), 2-benzyl-4,6-dichloro-pyrimidine (1.0 eq.), acetone (10.0 mL/mmol) and HCl (0.6 eq.) were added. The sealed tube was stirred at 130 °C for 1 hour in a microwave reactor. The reaction was monitored by LC-MS. The mixture was cooled, the precipitate was collected by filtration, washed with cold acetone and dried in vacuum to give the desired product, which was used without further purification.

**General Procedure L2** – *SN Ar V*

A microwave tube was charged with the appropriate chloride derivative (1.0 eq.), tert-butyl N-(4-aminocyclohexyl)carbamate (2.5 eq.), MeCN (12.0 mL/mmol), DMSO (3.0 mL/mmol) and DIPEA (2.0 eq.) were added. The sealed tube was stirred at 140 °C for 52 hours in a microwave reactor. The reaction was monitored by LC-MS. The mixture was purified by reversed-phase column chromatography using 5mM $NH_4HCO_3$ solution and MeCN as eluent to give the desired product.

**General Procedure M** – *Suzuki coupling*

A microwave tube was charged with the appropriate bromide derivative (1.0 eq.), the appropriate boronic acid (1.5-5.0 eq.), $(A\text{-}taPhos)_2PdCl_2$ (5.0 mol%), $Cs_2CO_3$ (3.0 eq.) dioxane (20.0 mL/mmol) and water (6.8 mL/mmol) were added under inert atmosphere. The sealed tube was stirred at 130 °C for 1.5 hours in a microwave reactor. The reaction was monitored by LC-MS. The mixture was concentrated, and the residue was purified by reversed-phase column chromatography using water and MeCN as eluent to give the desired product.

**General Procedure N** – *Nitrile-amide transformation*

The mixture of the appropriate nitrile (1.0 eq.), (1E)-acetaldehyde oxime (8.0 eq.), and copper(II) molecular sieve catalyst (100 mg/mmol) in MeOH (3.8 mL/mmol) was stirred at 65 °C for 3 days. The reaction was monitored by LC-MS. The mixture was concentrated and purified by normal-phase column chromatography using DCM and MeOH as eluent to give the desired product.

**Description of intermediates and final molecules**

*N4-[2-benzyl-7-(1-methylpyrazol-4-yl)-9H-pyrimido[4,5-b]indol-4-yl]cyclohexane-1,4-diamine (**2**)*

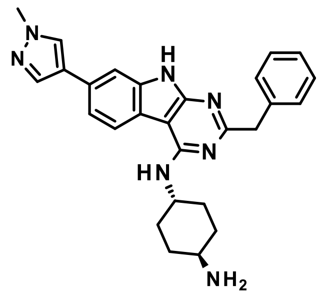

Title compound was synthesized according to the *General procedure I* using tert-butyl N-[4-[[2-benzyl-7-(1-methylpyrazol-4-yl)-9H-pyrimido[4,5-b]indol-4-yl]amino]cyclohexyl]carbamate (**21**, 77 mg, 0.14 mmol) to give the desired product (42 mg, 67%).

1H NMR (500 MHz, DMSO-$d_6$): δ ppm 11.71 (brs, 1 H), 8.22 (d, J = 8.2 Hz, 1 H), 8.17 (s, 1 H), 7.88 (d, J =0.8 Hz, 1 H), 7.48 (d, J = 1.1 Hz,1 H), 7.40 (dd, J = 6.8, 1.6 Hz, 1 H), 7.38 (d, J = 6.7 Hz, 2 H), 7.28 (t, J = 7.5 Hz, 2 H), 7.18 (t, J = 7.3 Hz, 1 H), 6.49 (d, J = 8.6 Hz, 1 H), 4.21 (m, 1 H), 4.02 (s, 2 H), 3.87 (s, 3 H), 2.57 (m, 1 H), 1.91/1.55 (d+dd, J = 10.1/22.7, 10.1

Hz, 4 H), 1.82/1.17 (d+dd, J = 12.0/24.0, 10.7 Hz, 4 H), 1.66 (brs, 2 H); 13C NMR (125 MHz, DMSO-$d_6$): δ 165.3, 157.1, 156.3, 140.0, 137.3, 136.4, 129.7, 129.1, 128.4, 128.2, 126.4, 123.2, 122.0, 118.3, 118.0, 107.2, 94.0, 50.5, 49.5, 46.2, 39.1, 36.1, 31.5; UV/Vis: $\lambda_{max}$ 210 nm; HRMS (m/z): [M+H]+ calcd. for $C_{27}H_{29}N_7$, 452.2562; found, 452.2558.

| tR | Peak type | % | Product type | MW | Ions | Comment |
|---|---|---|---|---|---|---|
| 1.90 | principal | **98.4** | **Product** | ***451.2484*** | [M+H]+=452.2558 (δ=0.2 ppm) | |

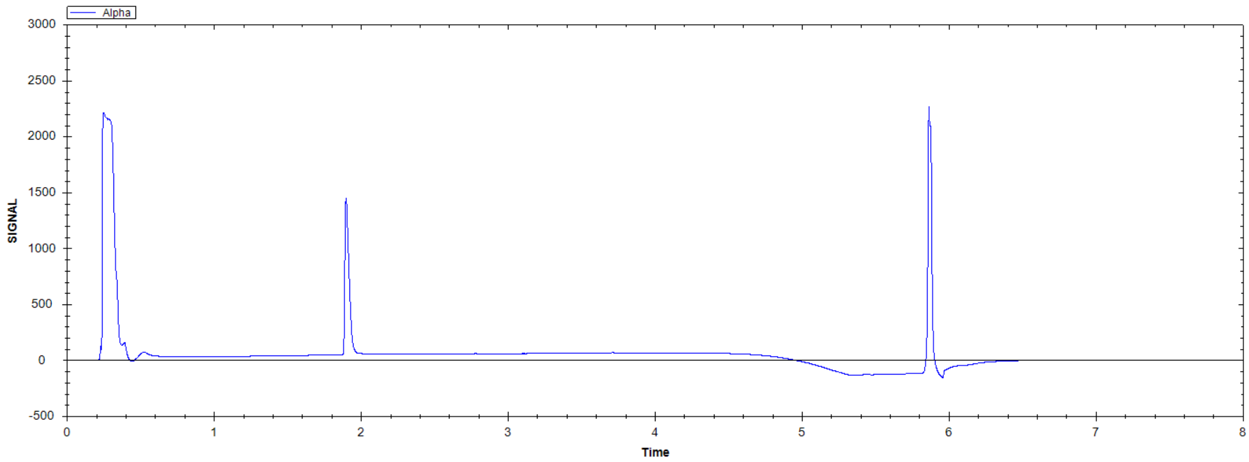


N4-[2-benzyl-7-(1-methylpyrazol-4-yl)-9H-pyrimido[4,5-b]indol-4-yl]cyclohexane-1,4-diamine

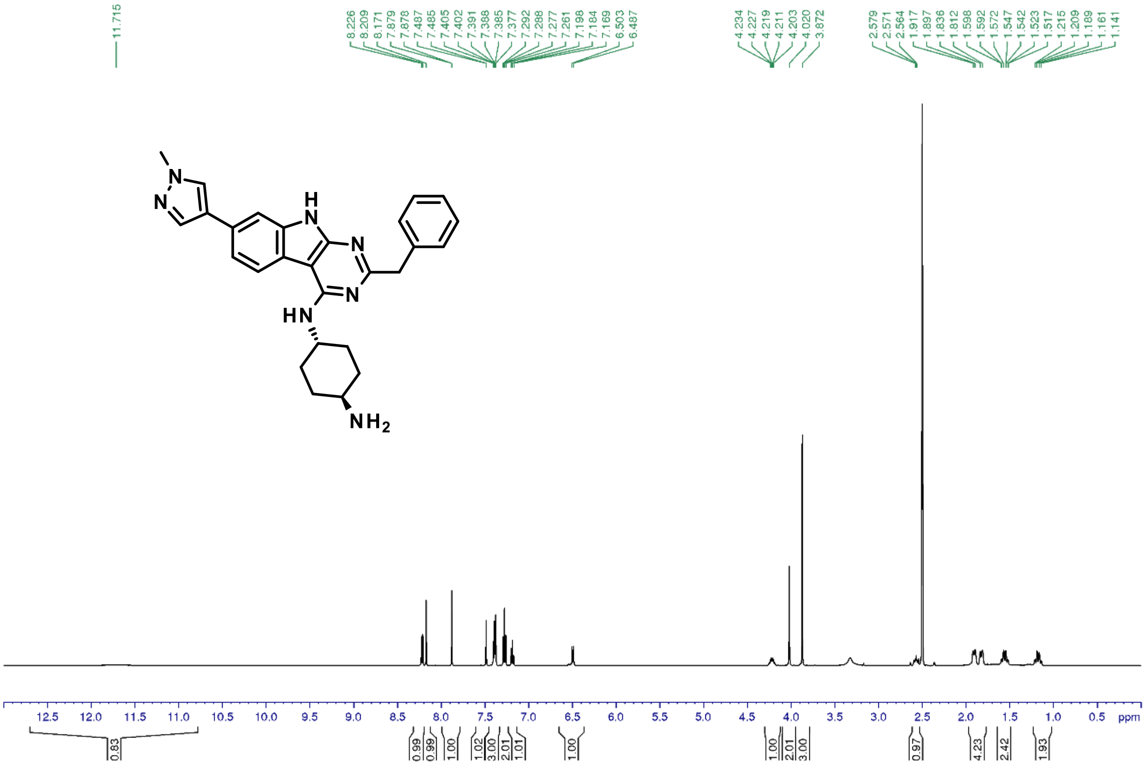

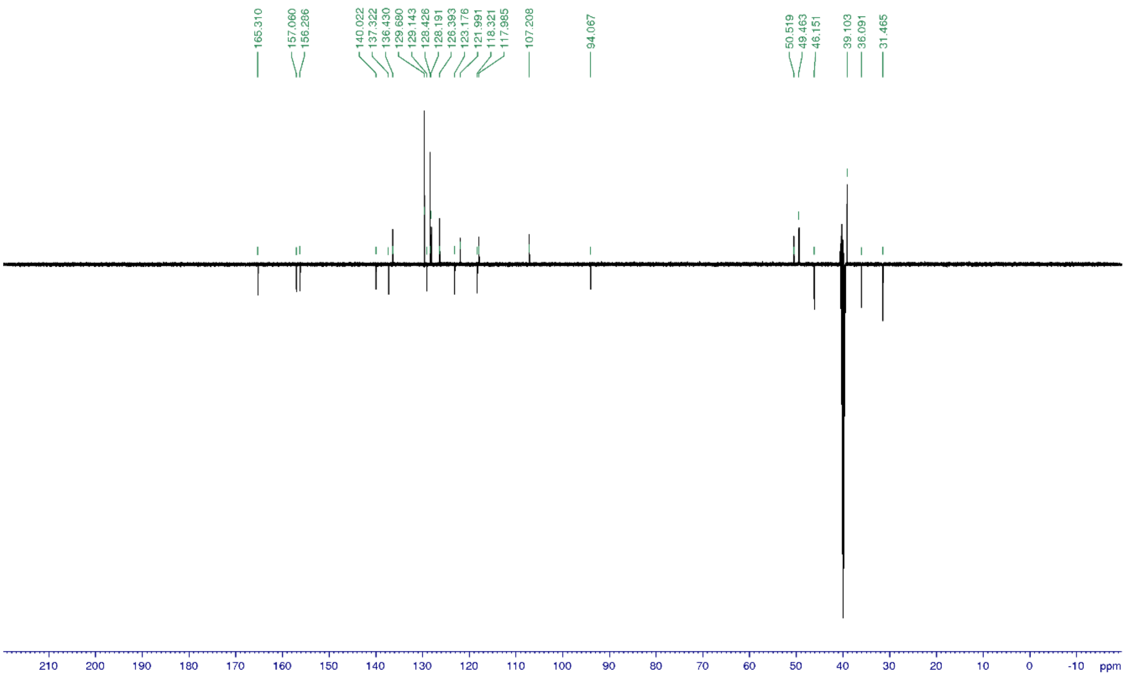

*N4-[2-benzyl-7-(3,5-dimethylisoxazol-4-yl)-9H-pyrimido[4,5-b]indol-4-yl]cyclohexane-1,4-diamine (**3**)*

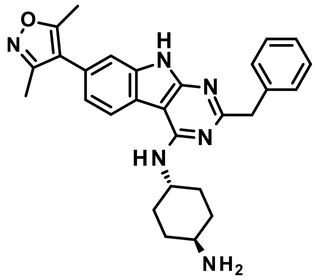

Title compound was synthesized according to the *General procedure I* using tert-butyl N-[4-[[2-benzyl-7-(3,5-dimethylisoxazol-4-yl)-9H-pyrimido[4,5-b]indol-4-yl]amino]cyclohexyl]carbamate (**22**, 19 mg, 0.03 mmol) to give the desired product (10 mg, 64%).

1H NMR (500 MHz, DMSO-$d_6$): δ ppm 11.80 (brs, 1 H), 8.35 (d, J = 8.3 Hz, 1 H), 7.39 (d, J = 7.6 Hz, 2 H), 7.32 (d, J = 1.1 Hz, 1 H), 7.28 (t, J = 7.3 Hz, 2 H), 7.19 (t, J = 7.4 Hz, 1 H), 7.17 (dd, J = 8.2, 1.6 Hz, 1 H), 6.61 (d, J = 8.1 Hz, 1 H), 4.22 (m, 1 H), 4.03 (s, 2 H), 2.57 (m, 1 H), 2.43 (s, 3 H), 2.25 (s, 3 H), 1.90/1.55 (d+dd, J = 10.6/24.0, 12.4 Hz, 4 H), 1.82/1.17 (d+dd, J = 12.4/24.2, 11.7 Hz, 4 H), 1.64 (brs, 2 H); 13C NMR (125 MHz, DMSO-$d_6$): δ 165.9, 165.2, 158.7, 157.2, 156.6, 139.9, 137.0, 129.7, 128.4, 126.4, 125.8, 121.9, 121.3, 119.4, 117.1, 111.4, 93.8, 50.5, 49.5, 46.1, 36.1, 31.4, 11.9, 11.1; UV/Vis: $\lambda_{max}$ 249 nm; HRMS (m/z): [M+H]+ calcd. for $C_{28}H_{30}N_6O$, 467.2559; found, 467.2557.

| tR | Peak type | % | Product type | MW | Ions | Comment |
|---|---|---|---|---|---|---|
| 2.17 | principal | **98.3** | **Product** | ***466.2481*** | [M+H]+=467.2557 (δ=0.7 ppm) | |

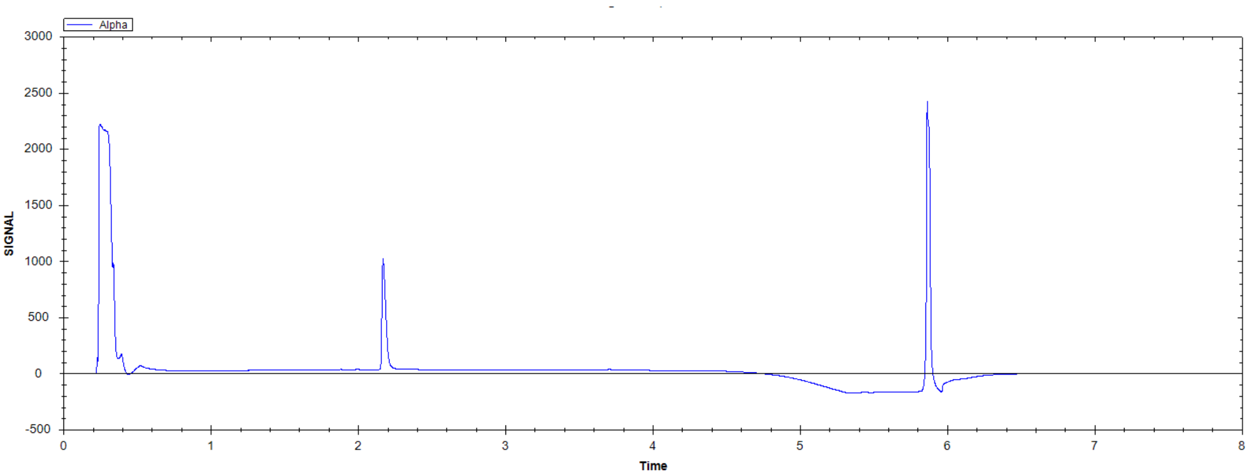


N4-[2-benzyl-7-(3,5-dimethylisoxazol-4-yl)-9H-pyrimido[4,5-b]indol-4-yl]cyclohexane-1,4-diamine

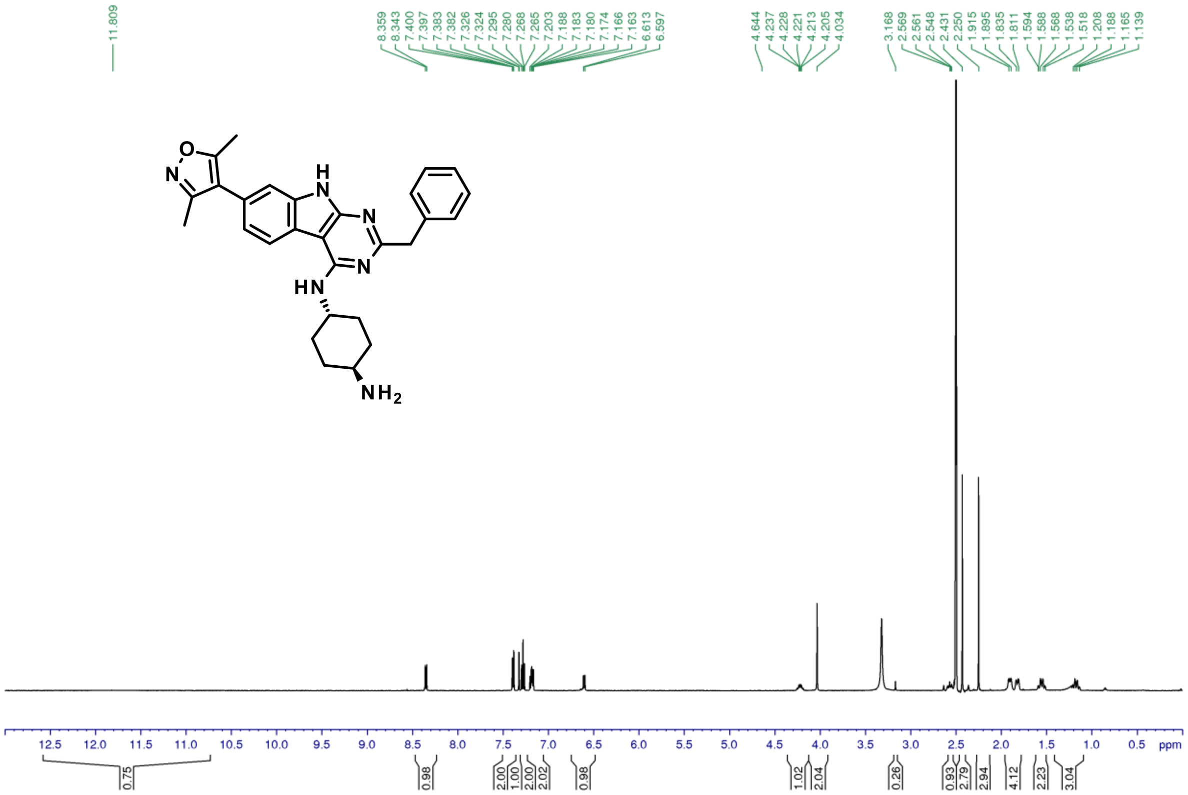

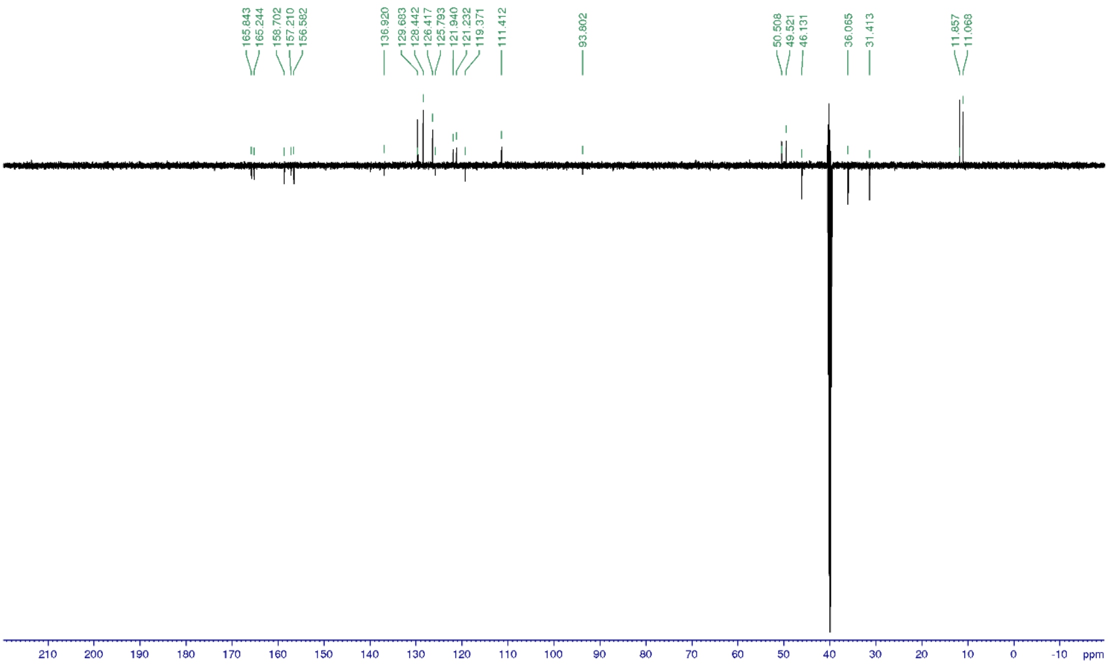


*Methyl 2-benzyl-4-[[4-(dimethylamino)cyclohexyl]amino]-9H-pyrimido[4,5-b]indole-7-carboxylate (**4**)*

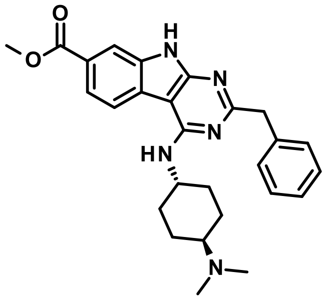


Title compound was synthesized according to the *General procedure H1* using methyl 2-benzyl-4-chloro-9H-pyrimido[4,5-b]indole-7-carboxylate (**29**, 45 mg, 0.1 mmol), N4,N4-dimethylcyclohexane-1,4-diamine (2.2 eq.) and DIPEA (15.0 eq.) to give the desired product (6 mg, 13%).

1H NMR (500 MHz, DMSO-$d_6$): δ ppm 12.13 (s, 1 H), 9.42 (brm, 1 H), 8.39 (d, J = 7.8 Hz, 1 H), 8.00 (d, J = 1.4 Hz,1 H), 7.82 (dd, J = 8.3, 1.5 Hz,1 H), 7.40-7.18 (m, 5 H), 7.01 (d, J = 7.8 Hz, 1 H), 4.26 (m, 1 H), 4.06 (s, 2 H), 3.88 (s, 3 H), 3.18 (m, 1 H), 2.81 (d, J = 5.0 Hz, 6 H), 2.15-1.54 (m, 8 H); 13C NMR (125 MHz, DMSO-$d_6$): δ 167.1, 121.4, 121.1, 112.5, 63.8, 52.5, 48.9, 45.9, 39.8, 30.1/25.4; UV/Vis: $\lambda_{max}$ 247 nm; HRMS (m/z): [M+H]+ calcd. for $C_{27}H_{31}N_5O_2$, 458.2556; found, 458.2553.

| tR | Peak type | % | Product type | MW | Ions | Comment |
|---|---|---|---|---|---|---|
| 1.89 | minor | **2.1** | | | | |
| 1.93 | principal | **96.6** | **Product** | ***457.2478*** | [M+H]+=458.2554 (δ=0.8 ppm) | |
| 2.02 | minor | **1.0** | | | | |

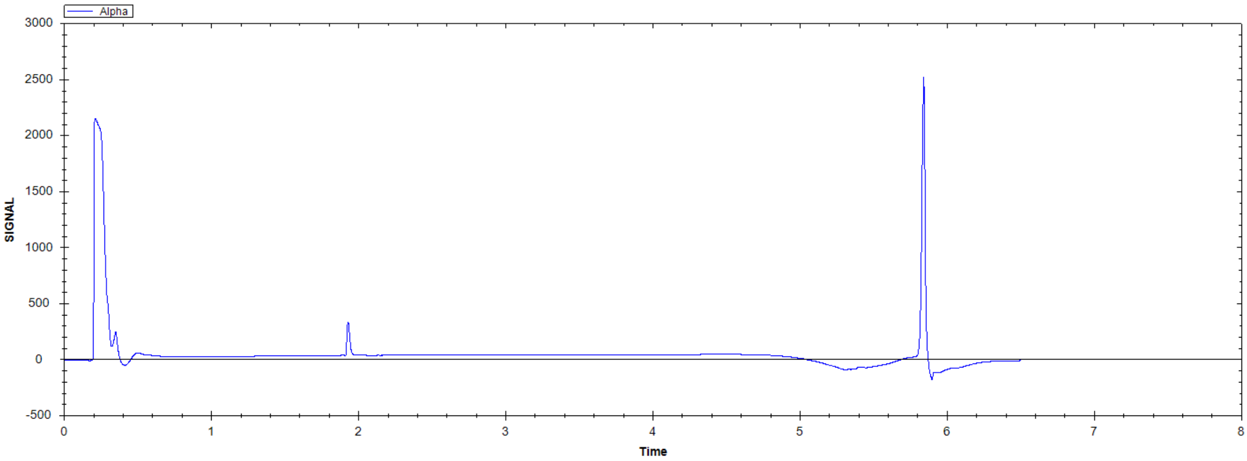


methyl 2-benzyl-4-[[4-(dimethylamino)cyclohexyl]amino]-9H-pyrimido[4,5-b]indole-7-carboxylate

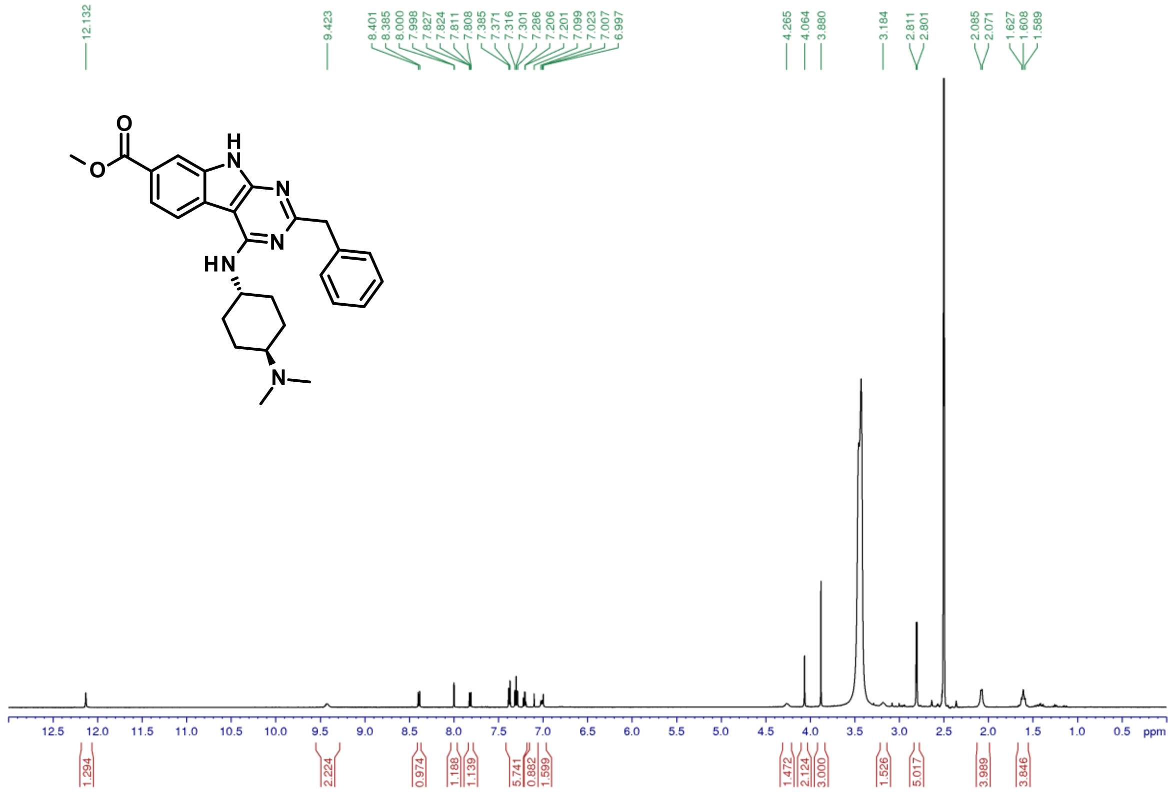

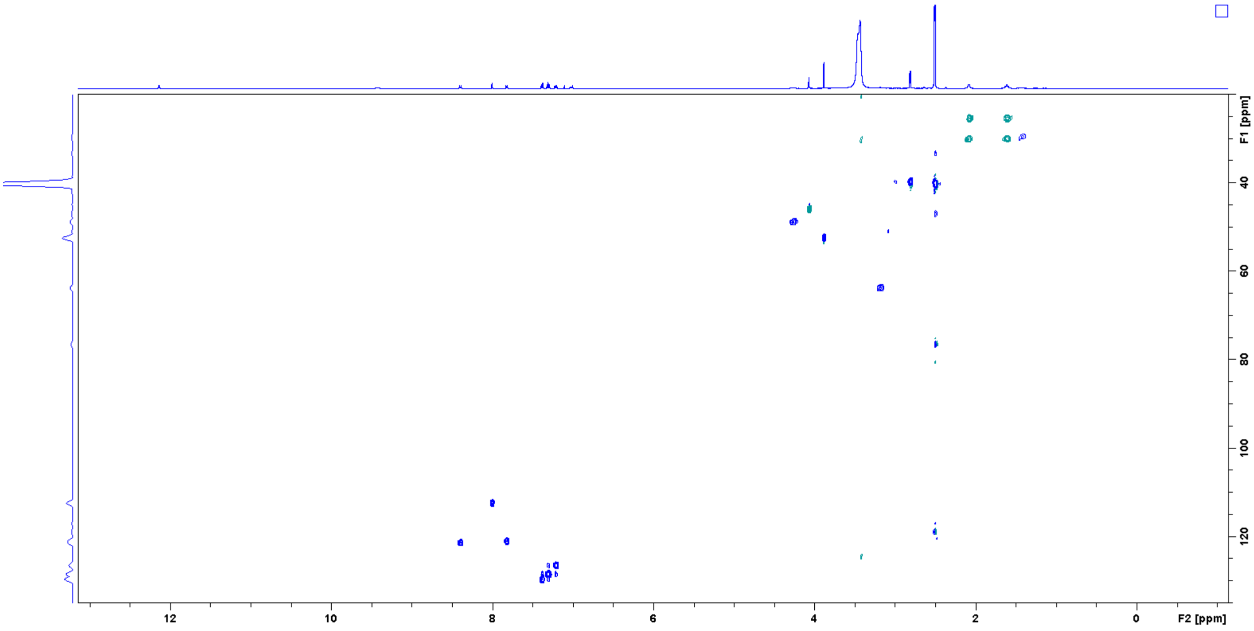


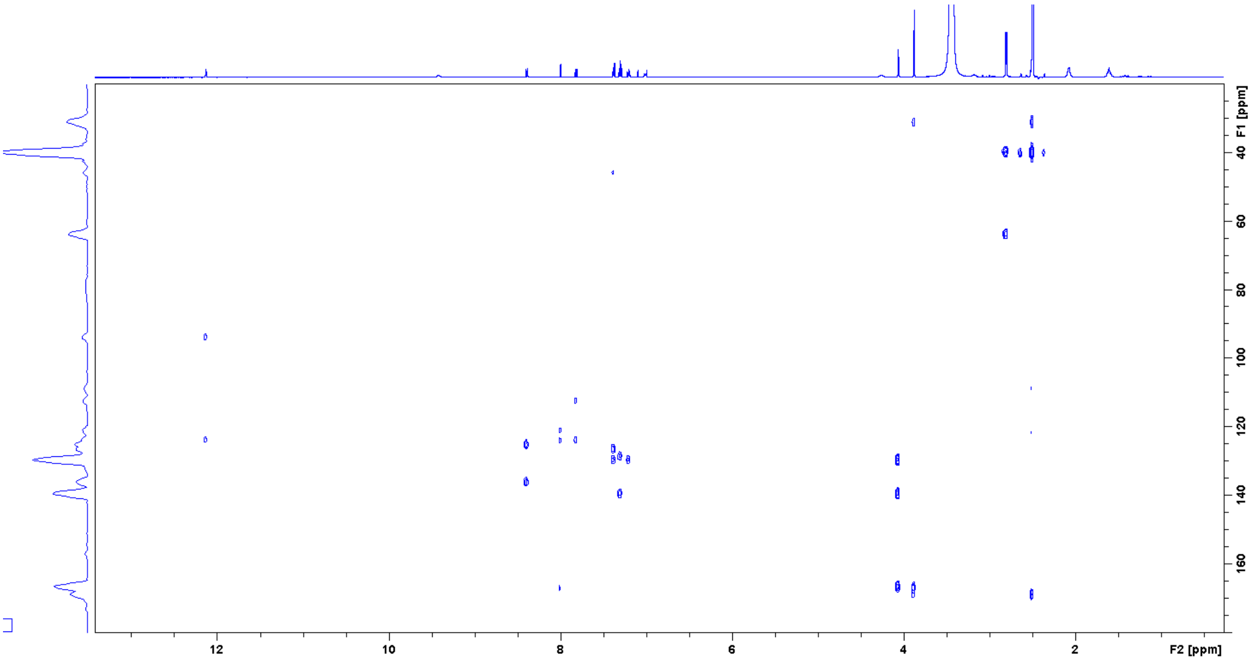


*2-Benzyl-4-[[4-(dimethylamino)cyclohexyl]amino]-9H-pyrimido[4,5-b]indole-7-carboxylic acid (**5**)*

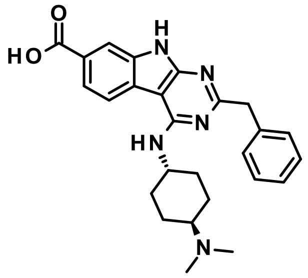

Title compound was synthesized according to the *General procedure J* using methyl 2-benzyl-4-[[4-(dimethylamino)cyclohexyl]amino]-9H-pyrimido[4,5-b]indole-7-carboxylate (**4**, 7 mg, 0.015 mmol) to give the desired product (4 mg, 58%).

1H NMR (500 MHz, DMSO-$d_6$): δ ppm 11.98 (s, 1 H), 8.35 (d, J = 8.4 Hz, 1 H), 7.97 (d, J = 1.0 Hz,1 H), 7.79 (dd, J = 8.3, 1.1 Hz, 1 H), 7.37 (d, J = 7.3 Hz, 2 H), 7.28 (t, J = 7.3 Hz, 2 H), 7.19 (t, J = 7.4Hz, 1 H), 6.8 (d, J = 7.8 Hz, 1 H), 4.21 (m, 1 H), 4.04 (s, 2 H), 2.30 (m, 1 H), 2.28 (s, 6 H), 2.00/1.56 (m+m, 4 H), 1.89/1.34 (m+m, 4 H); 13C NMR (125 MHz, DMSO-$d_6$): δ 168.4, 158.0, 156.8, 139.8, 129.7, 128.5, 126.4, 121.3, 121.1, 112.5, 63.1, 49.7, 46.1, 41.6, 31.3, 27.3; UV/Vis: $\lambda_{max}$ 248 nm; HRMS (m/z): [M+H]+ calcd. for $C_{26}H_{29}N_5O_2$, 444.2399; found, 444.2394.

| tR | Peak type | % | Product type | MW | Ions | Comment |
|---|---|---|---|---|---|---|
| 1.83 | principal | **96.9** | **Product** | ***443.2321*** | [M+H]+=444.2396 (δ=0.4 ppm) | |
| 1.99 | minor | **1.4** | | | | |

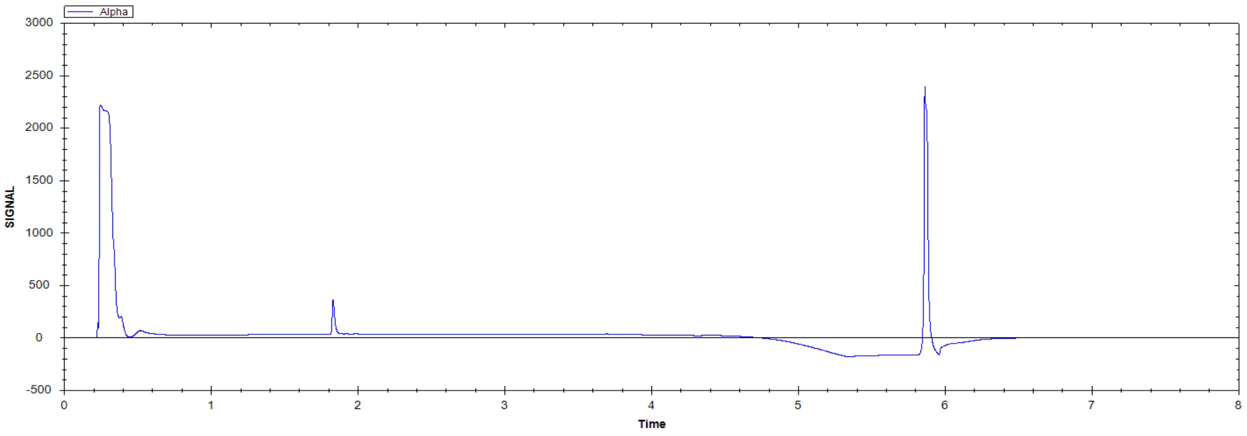

2-benzyl-4-[[4-(dimethylamino)cyclohexyl]amino]-9H-pyrimido[4,5-b]indole-7-carboxylic acid

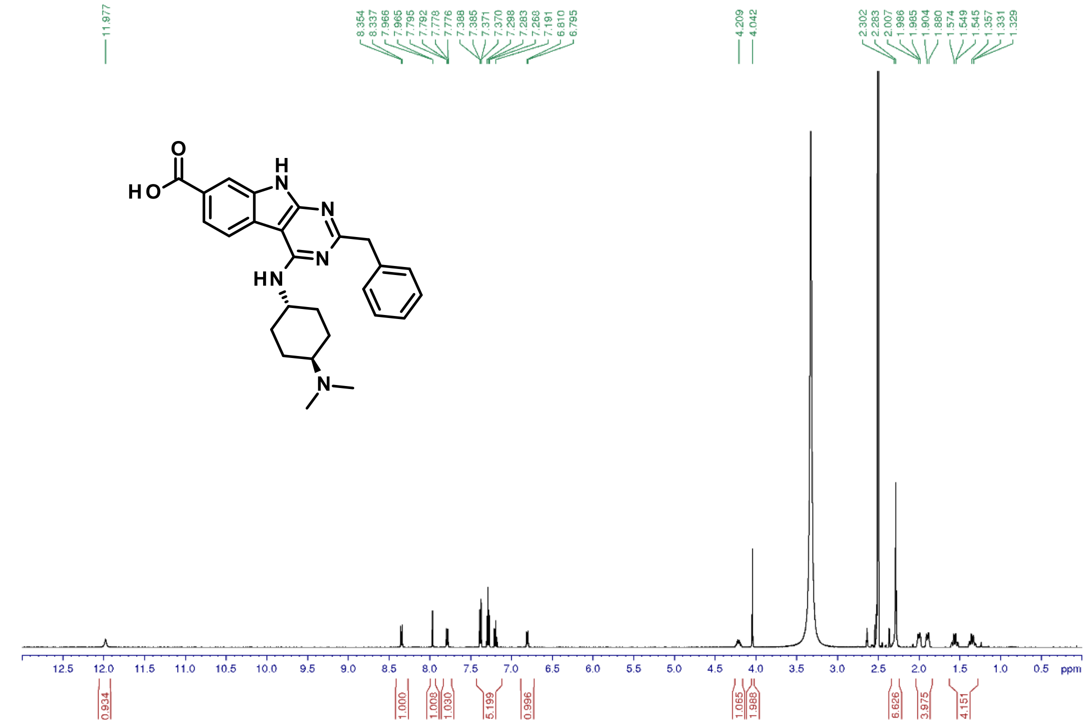


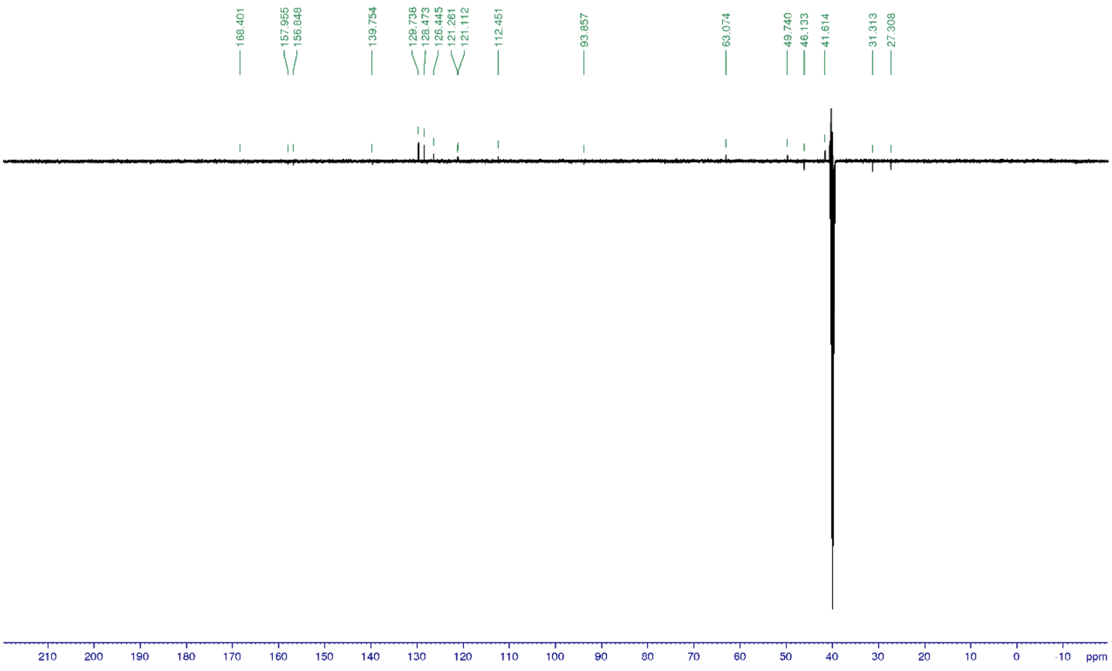


*2-Benzyl-4-[3-(1-piperidyl)propylamino]-9H-pyrimido[4,5-b]indole-7-carbonitrile* ***(6)***

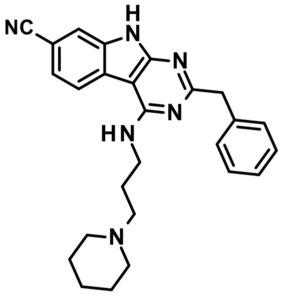


Title compound was synthesized according to the *General procedure H3* using 2-benzyl-4-chloro-9H-pyrimido[4,5-b]indole-7-carbonitrile (**45**, 36 mg, 0.11 mmol) and 3-(1-piperidyl)propan-1-amine (2.0 eq.) to give the desired product (43 mg, 89%).

1H NMR (500 MHz, DMSO-$d_6$): δ ppm 12.18 (brs, 1 H), 8.43 (d, J = 8.0 Hz, 1 H), 7.79 (d, J = 1.4 Hz, 1 H), 7.61 (dd, J = 8.2, 1.5 Hz, 1 H), 7.46 (t, J = 5.3 Hz, 1 H), 7.36 (d, J = 7.6 Hz, 2 H), 7.27 (t, J = 7.3 Hz, 2 H), 7.18 (t, J = 7.4 Hz, 1 H), 4.03 (s, 2 H), 3.62 (m, 2 H), 2.31 (t, J = 7.0 Hz, 2 H), 2.30 (brs, 4 H), 1.78 (m, 2 H), 1.48 (m, 4 H), 1.37 (m, 2 H); 13C NMR (125 MHz, DMSO-$d_6$): δ 167.6, 157.3, 135.7, 123.8, 123.6, 121.8, 120.5, 115.0, 105.5, 93.6, 57.0, 54.5, 46.3, 39.6, 26.9, 26.1, 24.6; UV/Vis: $\lambda_{max}$ 230 nm; HRMS (m/z): [M+H]+ calcd. for $C_{26}H_{28}N_6$, 425.2453; found, 425.2450.

| tR | Peak type | % | Product type | MW | Ions | Comment |
|---|---|---|---|---|---|---|
| 1.98 | principal | **96.8** | **Product** | ***424.2375*** | [M+H]+=425.245 (δ=0.4 ppm) | |
| 2.06 | minor | **1.7** | | | | |

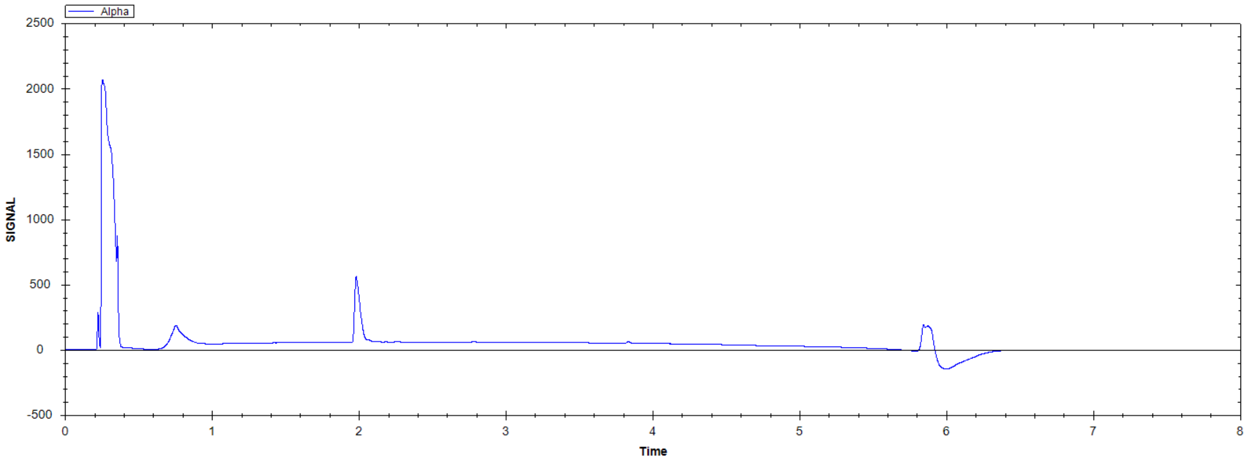

2-Benzyl-4-[3-(1-piperidyl)propylamino]-9H-pyrimido[4,5-b]indole-7-carbonitrile

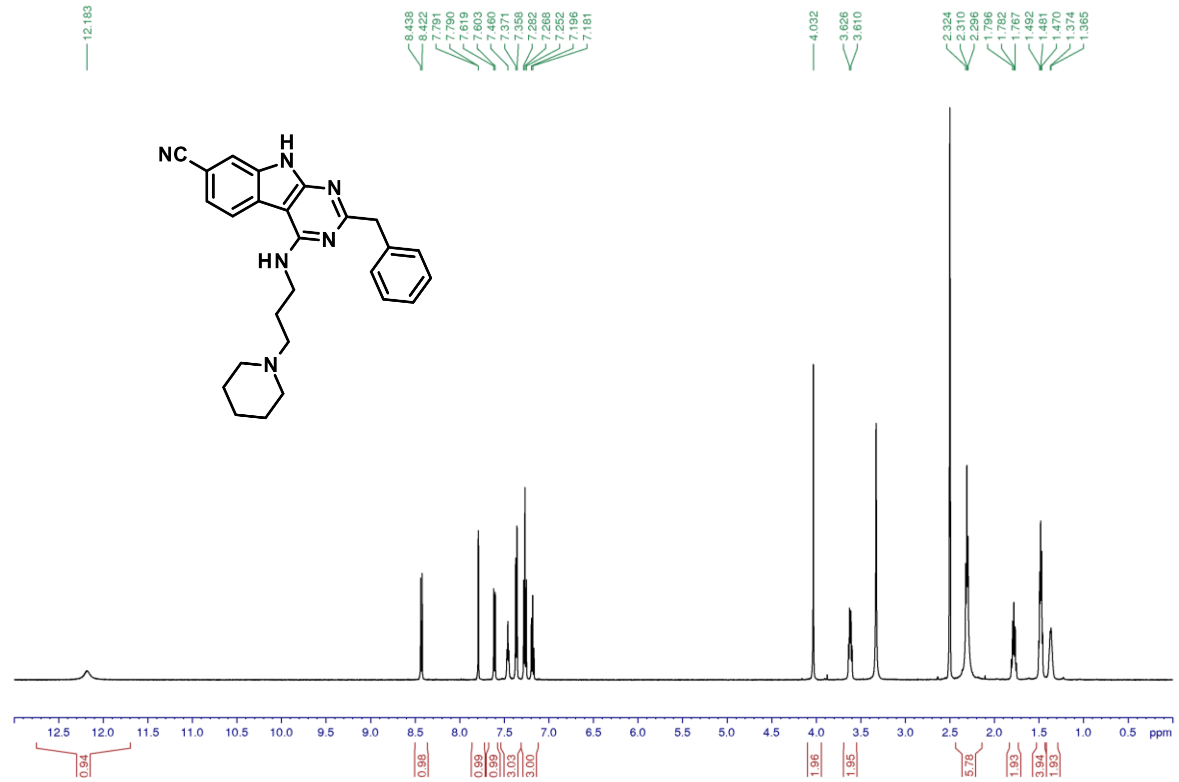

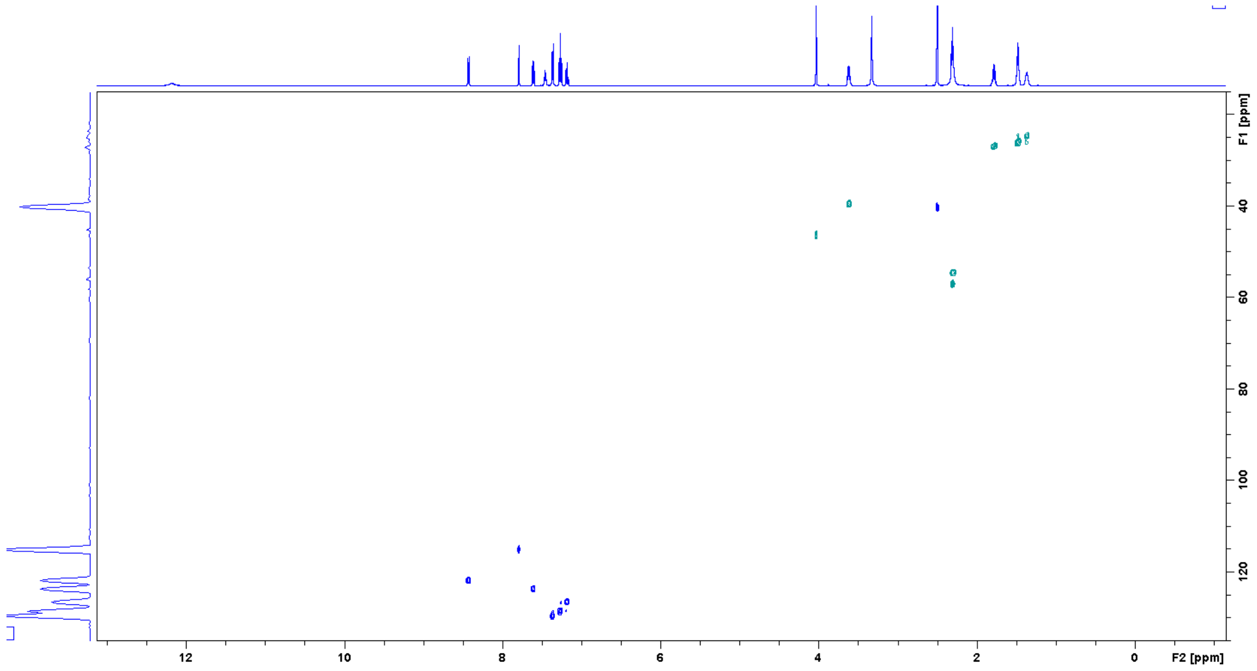

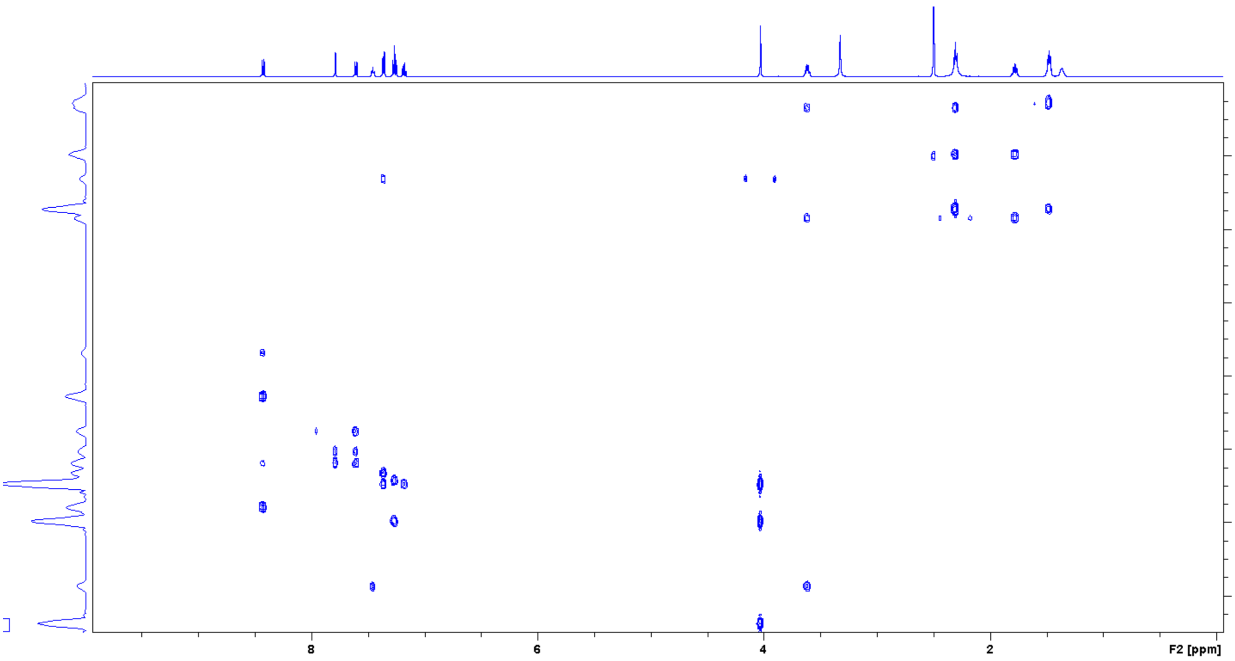


*Methyl 4-[(4-aminocyclohexyl)amino]-2-benzyl-9H-pyrimido[4,5-b]indole-6-carboxylate* (**7**)

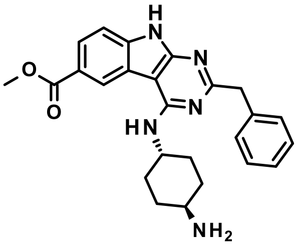


Title compound was synthesized according to the *General procedure I* using methyl 2-benzyl-4-[[4-(tert-butoxycarbonylamino)cyclohexyl]amino]-9H-pyrimido[4,5-b]indole-6-carboxylate (**35**, 68 mg, 0.13 mmol) to give the desired product (55 mg, 99%).

1H NMR (500 MHz, DMSO-$d_6$): δ ppm 8.88 (d, J = 1.1 Hz,1 H), 7.94 (dd, J = 8.4, 1.5 Hz, 1 H), 7.45 (d, J = 8.5 Hz, 1 H), 7.38 (d, J = 7.1 Hz, 2 H), 7.28 (t, J = 7.4 Hz, 2 H), 7.19 (t, J = 7.3 Hz, 1 H), 7.04 (d, J = 8.0 Hz, 1 H), 4.22 (m, 1 H), 4.03 (s, 2 H), 3.88 (s, 3 H), 2.58 (m, 1 H), 1.90/1.60 (m+m, 4 H), 1.83/1.17 (m+m, 4 H); 13C NMR (125 MHz, DMSO-$d_6$): δ 167.6, 166.3, 157.8, 156.7, 139.8, 129.7, 128.5, 126.5, 125.9, 123.6, 121.8, 120.0, 111.0, 94.2, 52.3, 50.5, 49.7, 49.5, 46.1, 36.1, 31.2; UV/Vis: $\lambda_{max}$ 255 nm; HRMS (m/z): [M+H]+ calcd. for $C_{25}H_{27}N_5O_2$, 430.2243; found, 430.2239.

| tR | Peak type | % | Product type | MW | Ions | Comment |
|---|---|---|---|---|---|---|
| 1.97 | principal | >99 | Product | 429.2165 | [M+H]+=430.2239 (δ=0.3 ppm) | |

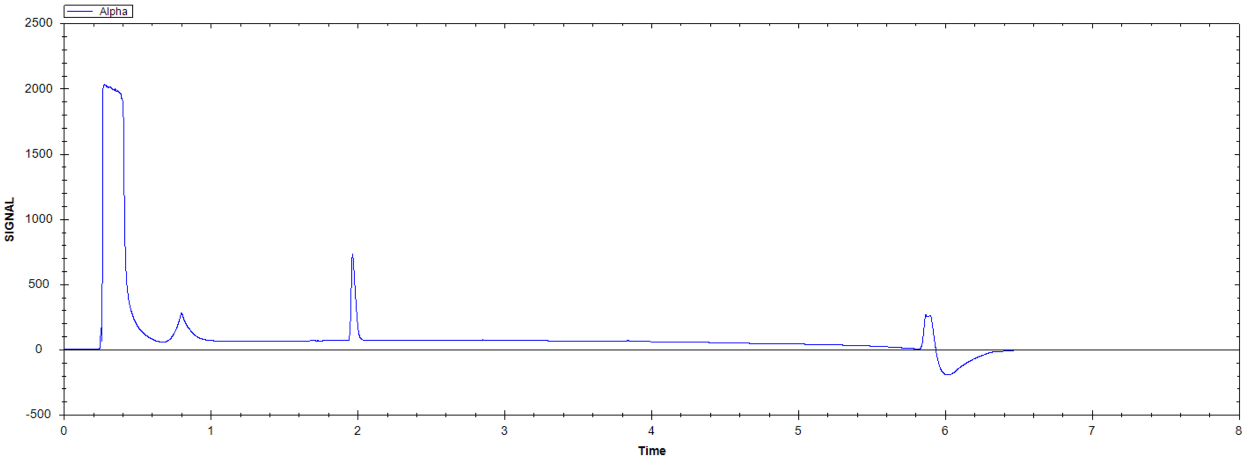


Methyl 4-[(4-aminocyclohexyl)amino]-2-benzyl-9H-pyrimido[4,5-b]indole-6-carboxylate

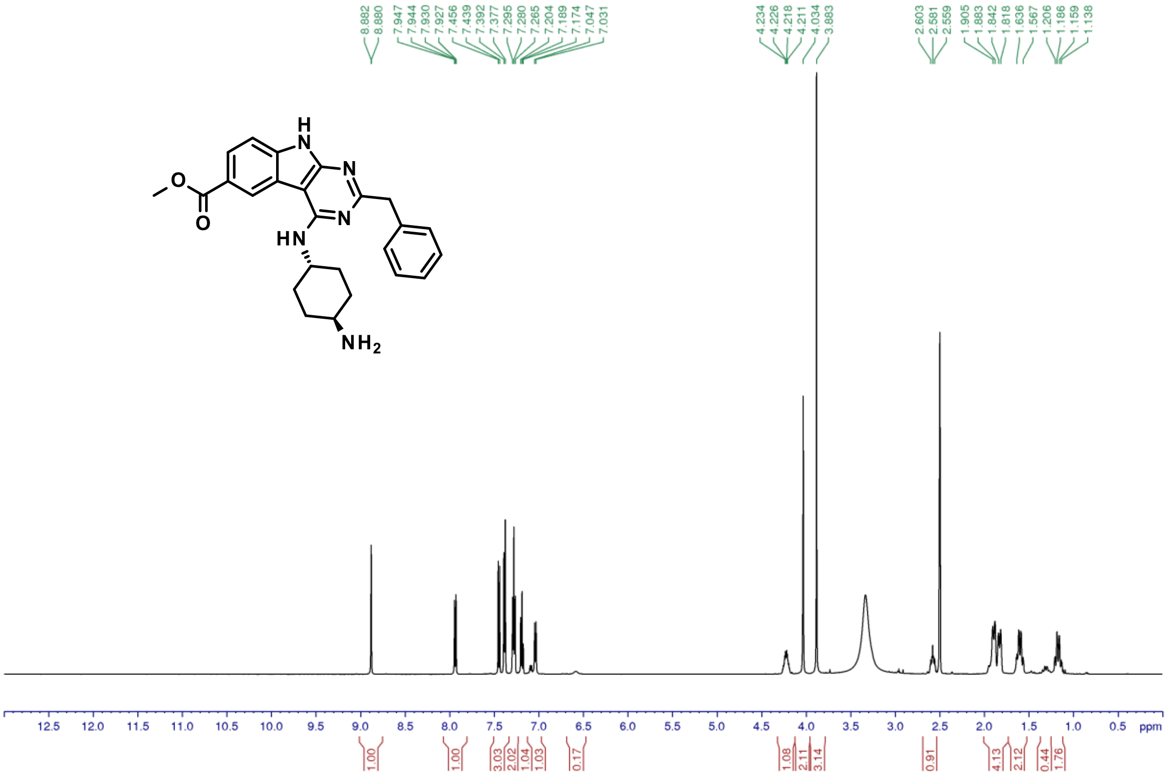

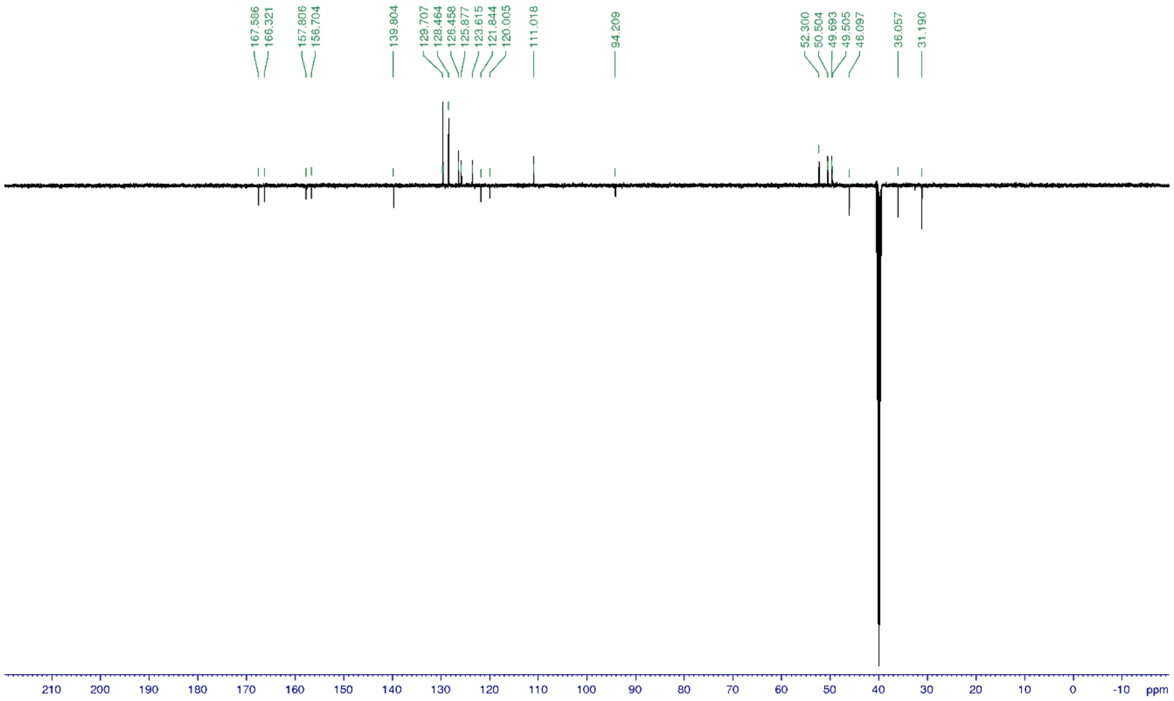


*N4,N4-dimethyl-N1-[7-(2-methyltetrazol-5-yl)-9H-pyrimido[4,5-b]indol-4-yl]cyclohexane-1,4-diamine (**8**)*

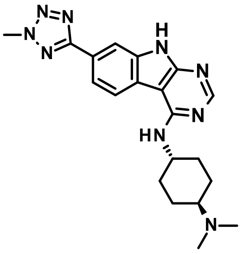

Title compound was synthesized according to the *General procedure H1* using 4-chloro-7-(2-methyltetrazol-5-yl)-9H-pyrimido[4,5-b]indole (**42**, 35 mg, 0.12 mmol), N4,N4-dimethylcyclohexane-1,4-diamine (3.0 eq.) and DBU (15.0 eq.) to give the desired product (14 mg, 29%).

1H NMR (500 MHz, DMSO-$d_6$): δ ppm 12.28 (s, 1 H), 9.59 (brm, 1 H), 8.54 (d, J = 8.3 Hz, 1 H), 8.42 (s, 1 H), 8.15 (d, J = 1.1 Hz, 1 H), 7.96 (dd, J = 8.3, 1.3 Hz, 1 H), 7.10 (brs, 1 H), 4.45 (s, 3 H), 4.35 (m, 1 H), 3.21 (m, 1 H), 2.79 (d, J = 5.0 Hz, 6 H), 2.18-1.57 (m, 8 H); 13C NMR (125 MHz, DMSO-$d_6$): δ 154.9, 122.6, 118.6, 109.4, 63.8, 48.7, 40.2, 39.8, 30.3/25.4; UV/Vis: $\lambda_{max}$ 255 nm; HRMS (m/z): [M+H]+ calcd. for $C_{20}H_{25}N_9$, 392.2311; found, 392.2309.

| tR | Peak type | % | Product type | MW | Ions | Comment |
|---|---|---|---|---|---|---|
| 1.36 | secondary | **20.9** | **By-Product** | | [M+H]+ m/z=266.22 | |
| 1.40 | minor | **2.6** | | | | |
| 1.43 | minor | **1.6** | | | | |
| 1.53 | minor | **1.5** | | | | |
| 1.56 | principal | **67.7** | **Product** | ***391.2233*** | [M+H]+=392.2309 (δ=0.8 ppm) | |
| 1.59 | minor | **2.2** | | | | |
| 1.82 | minor | **2.4** | | | | |

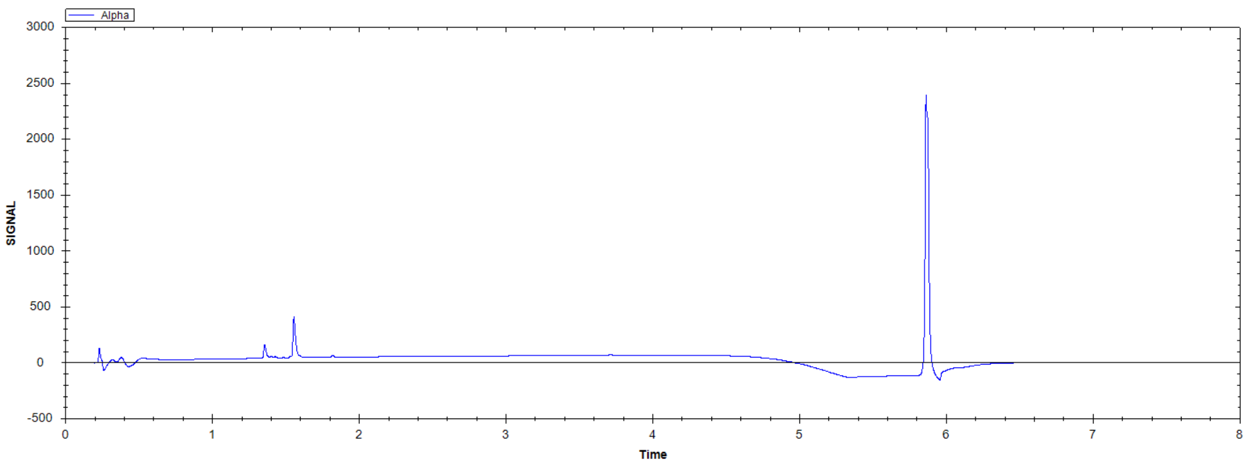


l4,N4-dimethyl-N1-[7-(2-methyltetrazol-5-yl)-9H-pyrimido[4,5-b]indol-4-yl]cyclohexane-1,4-diamine

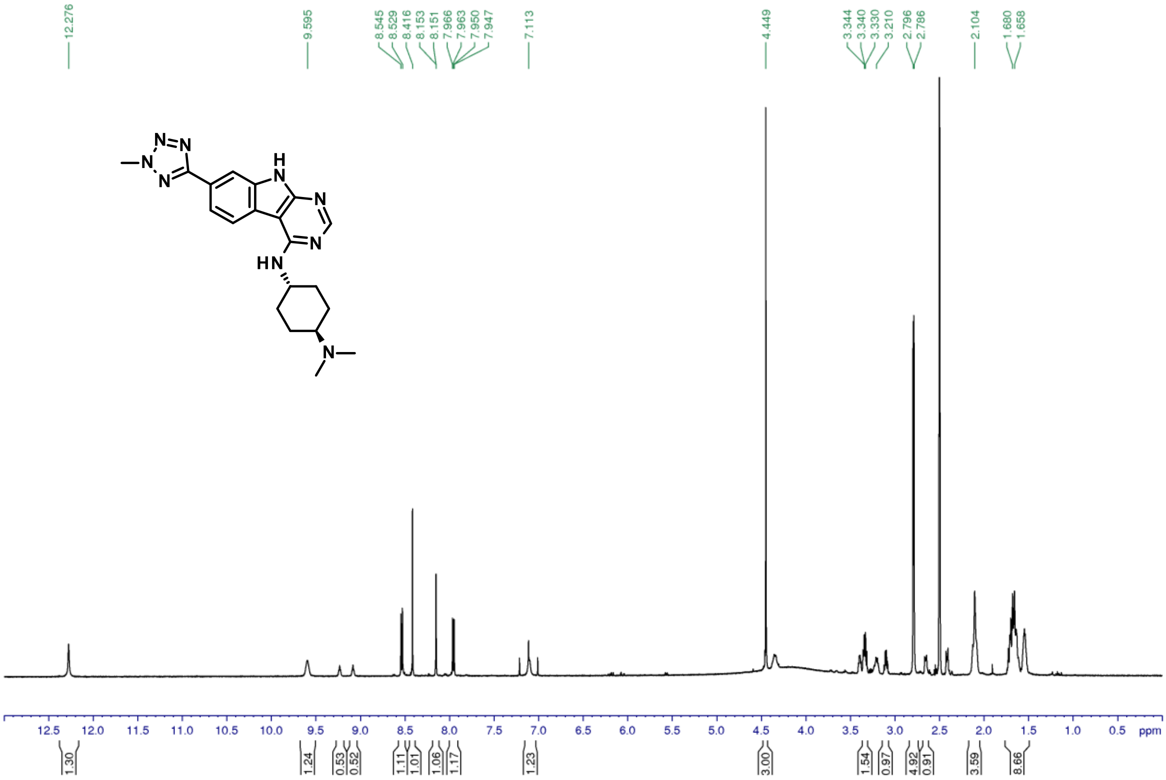

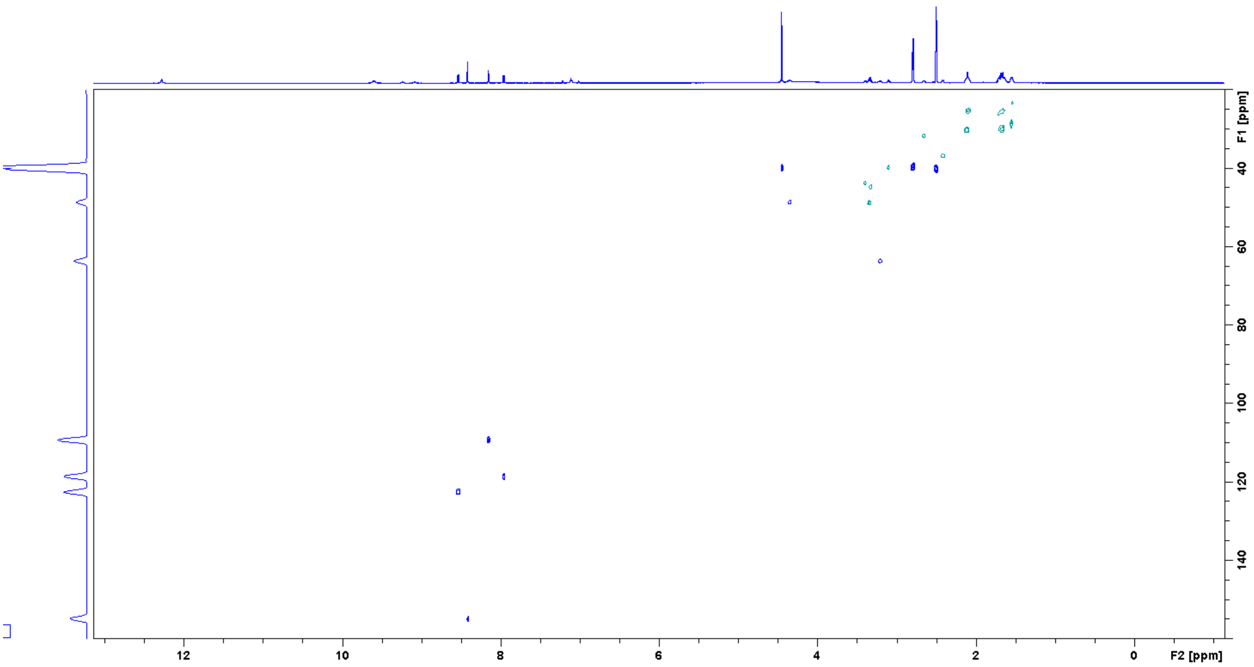


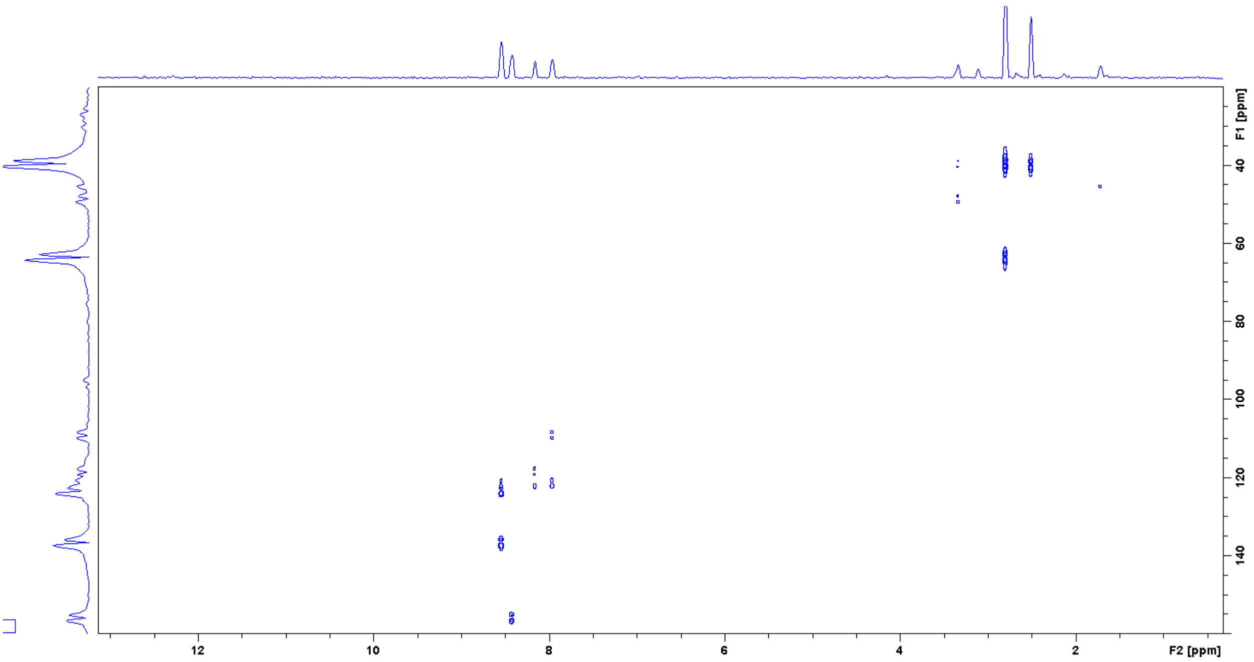


*N1-[2-benzyl-7-(2-methyltetrazol-5-yl)-9H-pyrimido[4,5-b]indol-4-yl]-N4,N4-dimethyl-cyclohexane-1,4-diamine (**9**)*

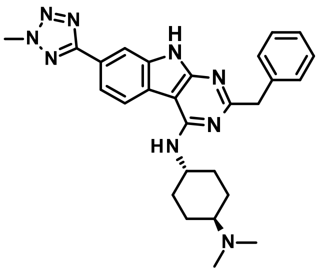

Title compound was synthesized according to the *General procedure H1* using 2-benzyl-4-chloro-7-(2-methyltetrazol-5-yl)-9H-pyrimido[4,5-b]indole (**40**, 20 mg, 0.04 mmol), 3-(1-

piperidyl)propan-1-amine (3.0 eq.) and DIPEA (15.0 eq.) to give the desired product (4 mg, 21%).

1H NMR (500 MHz, DMSO-$d_6$): δ ppm 12.09 (s, 1 H), 9.94 (brm, 1 H), 8.45 (d, J = 8.1 Hz, 1 H), 8.09 (d, J = 1.1 Hz,1 H), 7.91 (dd, J = 8.3, 1.7 Hz, 1 H), 7.39 (d, J = 7.1 Hz, 2 H), 7.31 (t, J = 7.6 Hz, 2 H), 7.21 (t, J = 7.6 Hz, 1 H), 6.93 (d, J = 8.3 Hz, 1 H), 4.44 (s, 3 H), 4.29 (brm, 1 H), 4.07 (s, 2 H), 3.19 (m, 1 H), 2.82 (d, J = 4.9 Hz, 6 H), 2.11/1.64 (m+m, 4 H), 2.09/1.61 (m+m, 4 H); 13C NMR (125 MHz, DMSO-$d_6$): δ 165.3, 139.7, 136.8, 129.7, 128.6, 126.5, 123.0, 122.3, 121.8, 118.3, 109.1, 63.8, 48.8, 46.0, 40.1, 39.9, 30.2, 25.5; UV/Vis: $\lambda_{max}$ 234 nm; HRMS (m/z): [M+H]+ calcd. for $C_{27}H_{31}N_9$, 482.2780; found, 482.2777.

| tR | Peak type | % | Product type | MW | Ions | Comment |
|---|---|---|---|---|---|---|
| 1.83 | secondary | **15.1** | **By-Product** | | [M+H]+ m/z=287.17 | |
| 1.99 | minor | **1.5** | | | | |
| 2.14 | principal | **73.9** | **Product** | ***481.2702*** | [M+H]+=482.2777 (δ=0.4 ppm) | |
| 2.18 | minor | **1.8** | | | | |
| 2.48 | minor | **3.8** | **By-Product** | | [M+H]+ m/z=315.21 | |
| 2.98 | minor | **1.5** | | | | |

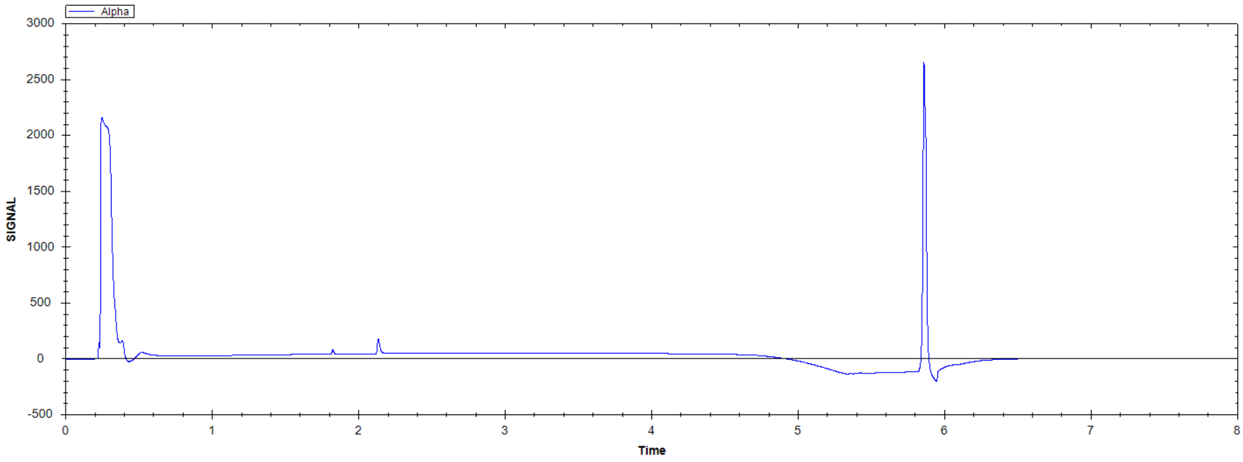

N1-[2-benzyl-7-(2-methyltetrazol-5-yl)-9H-pyrimido[4,5-b]indol-4-yl]-N4,N4-dimethyl-cyclohexane-1,4-diamine

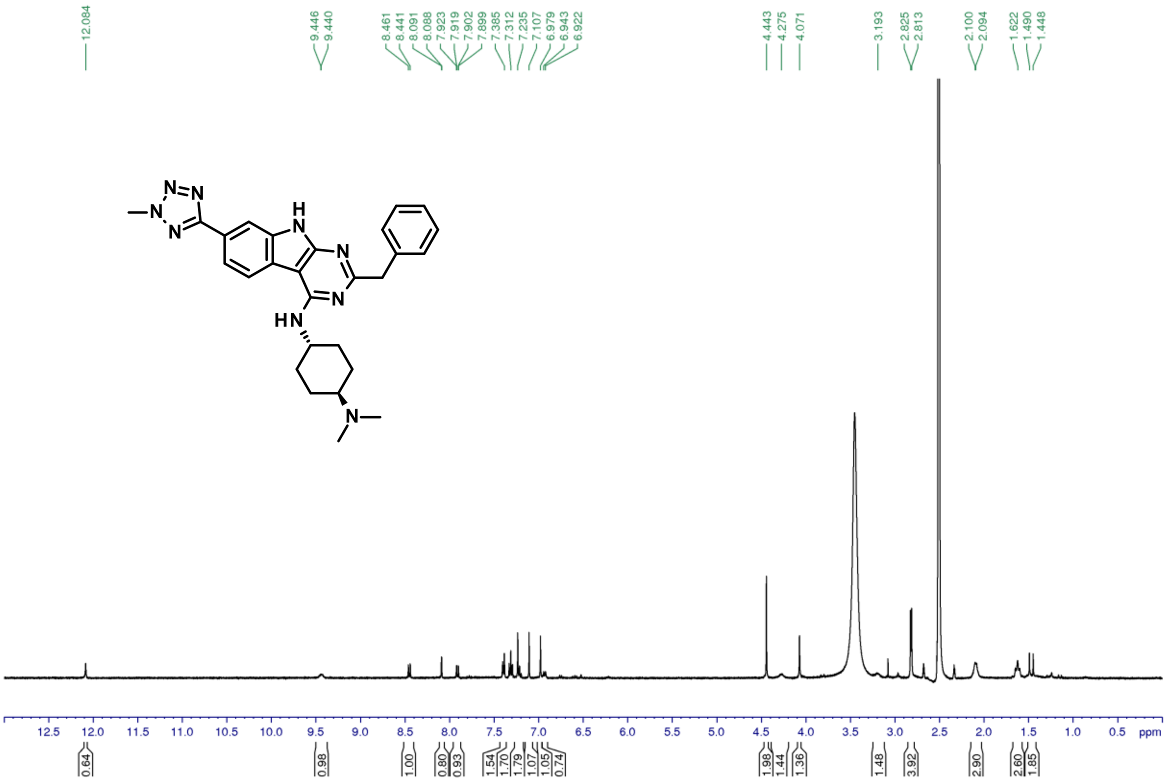


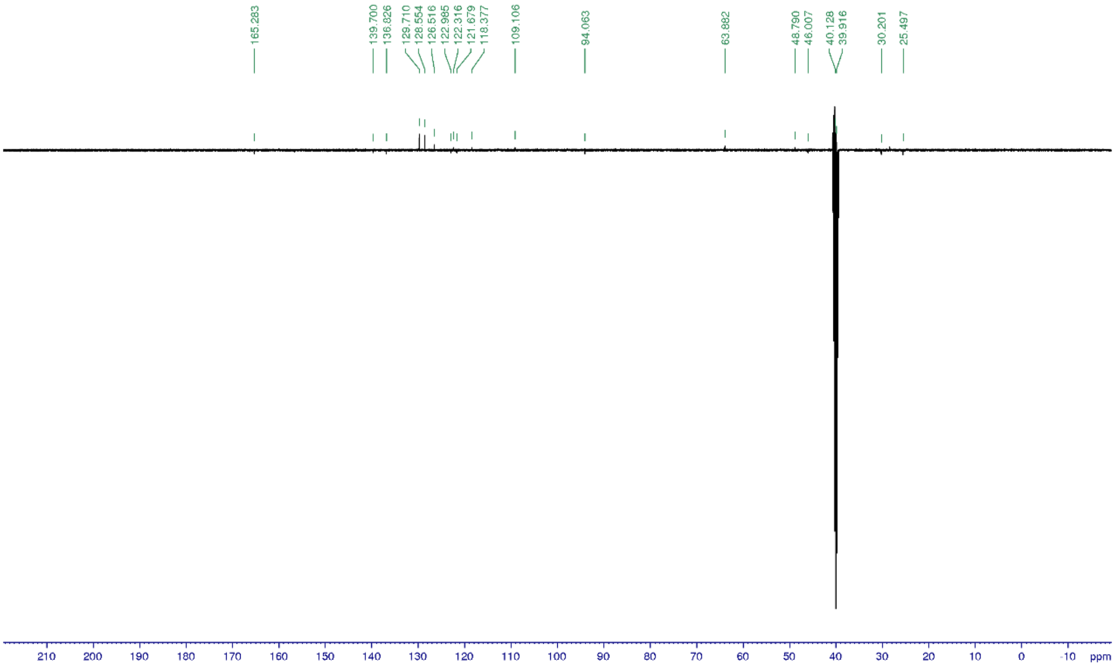


*2-Benzyl-N,N-dimethyl-7-(2-methyltetrazol-5-yl)-9H-pyrimido[4,5-b]indol-4-amine (**10**)*

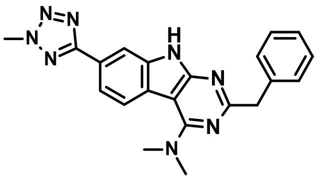

Title compound was synthesized according to the *General procedure H1* using 2-benzyl-4-chloro-7-(2-methyltetrazol-5-yl)-9H-pyrimido[4,5-b]indole (**40**, 20 mg, 0.04 mmol) and DIPEA (15.0 eq.) to give the desired product (4 mg, 24%).

1H NMR (500 MHz, DMSO-$d_6$): δ ppm 12.27 (s, 1 H), 8.11 (d, J = 1.3 Hz, 1 H), 8.05 (d, J = 8.7 Hz, 1 H), 7.92 (dd, J = 8.3, 1.5 Hz, 1 H), 7.39 (d, J = 7.4 Hz, 2 H), 7.29 (t, J = 7.4 Hz, 2 H), 7.2 (t, J = 7.4 Hz, 1 H), 4.44 (s, 3 H), 4.10 (s, 2 H), 3.32 (s, 6 H); 13C NMR (125 MHz, DMSO-$d_6$): δ 165.1, 164.5, 160.0, 139.2, 137.3, 129.6, 128.7, 126.6, 123.6, 123.1, 121.9, 119.0, 109.4, 95.5, 45.5, 40.2, 40.1; UV/Vis: $\lambda_{max}$ 236 nm; HRMS (m/z): [M+H]+ calcd. for $C_{21}H_{20}N_8$, 385.1889; found, 385.1885.

| tR | Peak type | % | Product type | MW | Ions | Comment |
|---|---|---|---|---|---|---|
| 2.37 | secondary | **11.7** | **By-Product** | | [M+H]+ m/z=286.06 | |
| 2.47 | minor | **2.6** | | | | |
| 3.17 | principal | **82.4** | **Product** | ***384.1811*** | [M+H]+=385.1885 (δ=0.3 ppm) | |

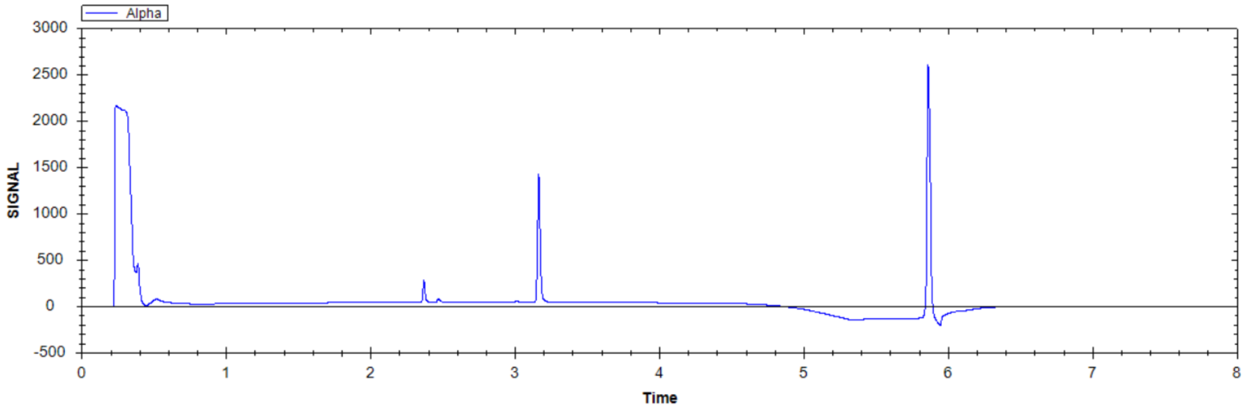

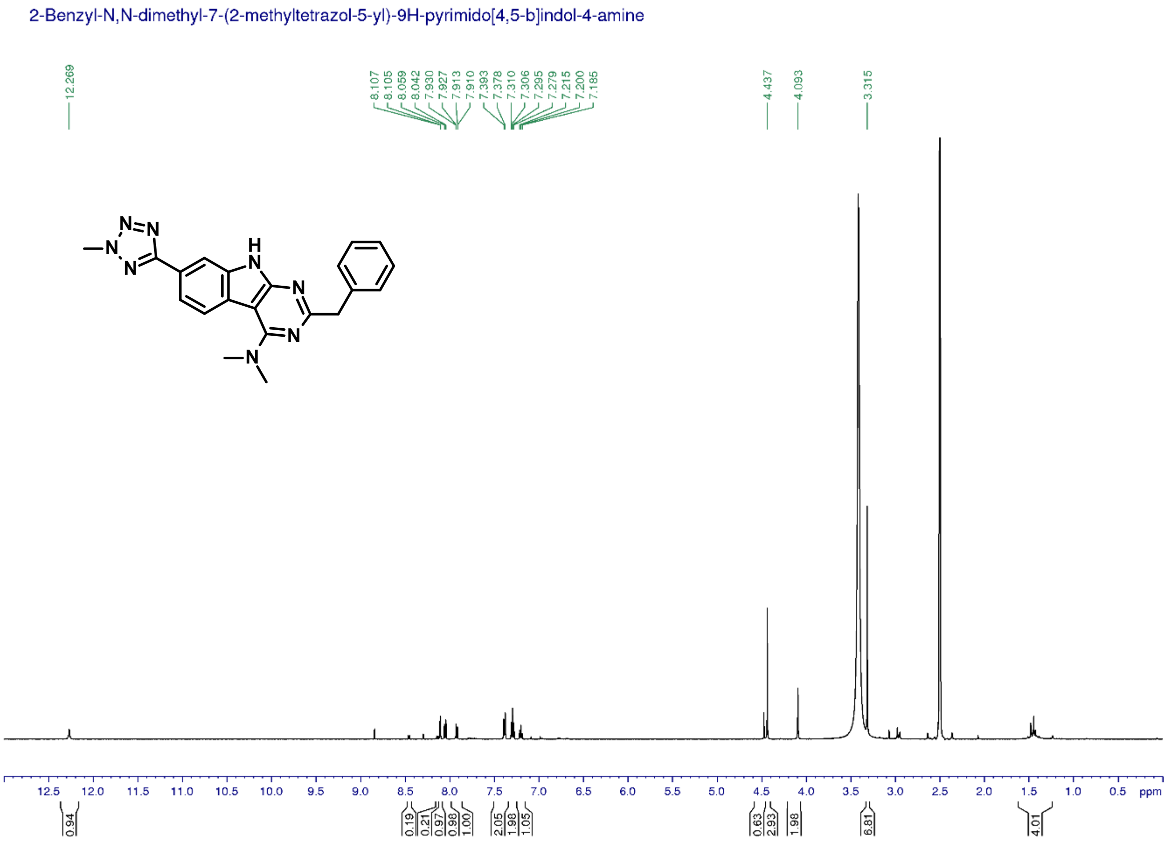


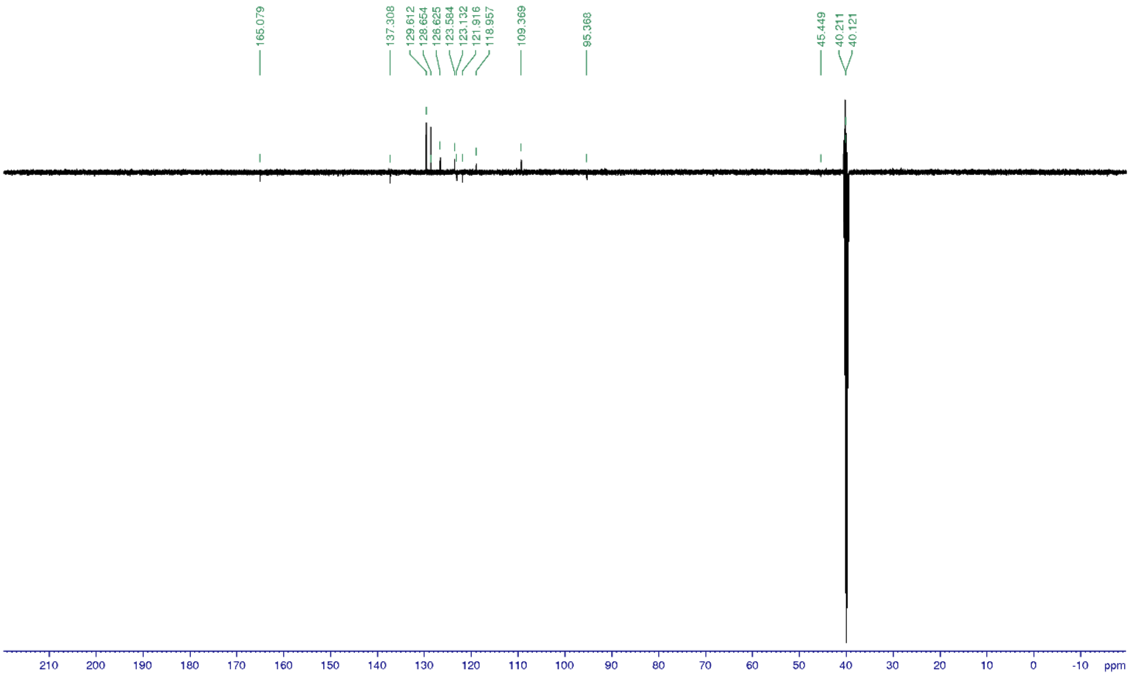


*2-Benzyl-7-(2-methyltetrazol-5-yl)-N-(3-pyrrolidin-1-ylpropyl)-9H-pyrimido[4,5-b]indol-4-amine (**11**)*

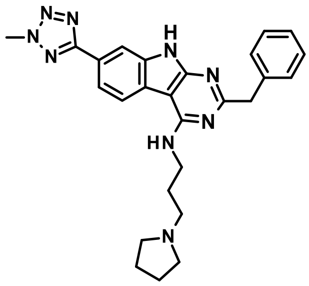

Title compound was synthesized according to the *General procedure H1* using 2-benzyl-4-chloro-7-(2-methyltetrazol-5-yl)-9H-pyrimido[4,5-b]indole (**40**, 49 mg, 0.13 mmol), 3-pyrrolidin-1-ylpropan-1-amine (3.5 eq.) and DIPEA (15.0 eq.) to give the desired product (22 mg, 36%).

1H NMR (500 MHz, DMSO-$d_6$): δ ppm 12.13 (s, 1 H), 9.44 (brs, 1 H), 8.42 (d, J = 8.2 Hz, 1 H), 8.10 (d, J = 1.0 Hz, 1 H), 7.93 (dd, J = 8.2, 1.5 Hz, 1 H), 7.47 (t, J = 5.4 Hz, 1 H), 7.38 (d, J = 7.6 Hz, 2 H), 7.31 (t, J = 7.4 Hz, 2 H), 7.21 (t, J = 7.4 Hz, 1 H), 4.44 (s, 3 H), 4.09 (s, 2 H), 3.70 (q, J = 6.0 Hz, 2 H), 3.52/2.91 (br+br, 4 H), 3.18 (m, 2 H), 2.02 (qn, J = 6.7 Hz, 2 H), 2.00/1.84 (br+br, 4 H); 13C NMR (125 MHz, DMSO-$d_6$): δ 166.1, 165.3, 158.3, 157.0, 139.6, 136.9, 129.6, 128.7, 126.6, 123.1, 121.9, 121.7, 118.6, 109.3, 94.1, 53.7, 52.6, 46.0, 40.1, 37.9, 26.6, 23.0; UV/Vis: $\lambda_{max}$ 234 nm; HRMS (m/z): [M+H]+ calcd. for $C_{26}H_{29}N_9$, 468.2624; found, 468.2621.

| tR | Peak type | % | Product type | MW | Ions | Comment |
|---|---|---|---|---|---|---|
| 2.15 | principal | >99 | **Product** | ***467.2546*** | [M+H]+=468.2621 (δ=0.5 ppm) | |

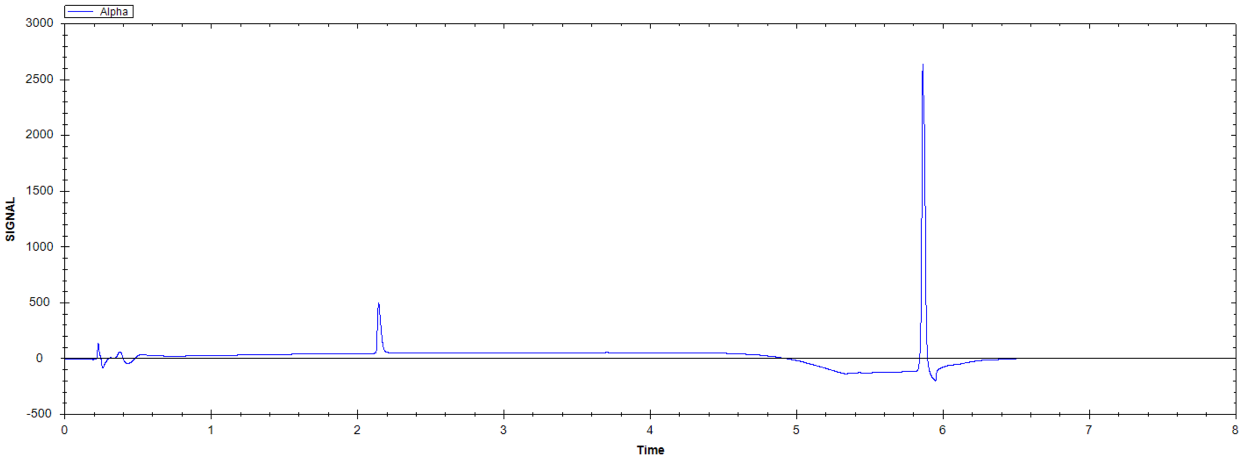

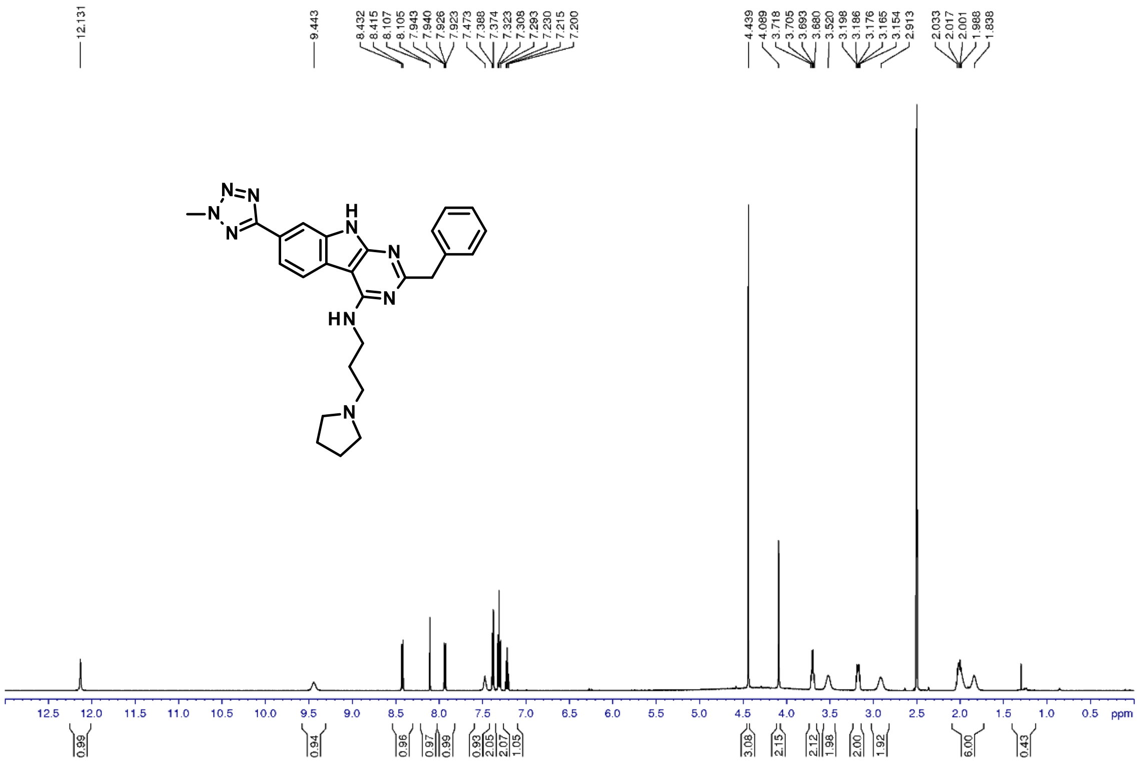


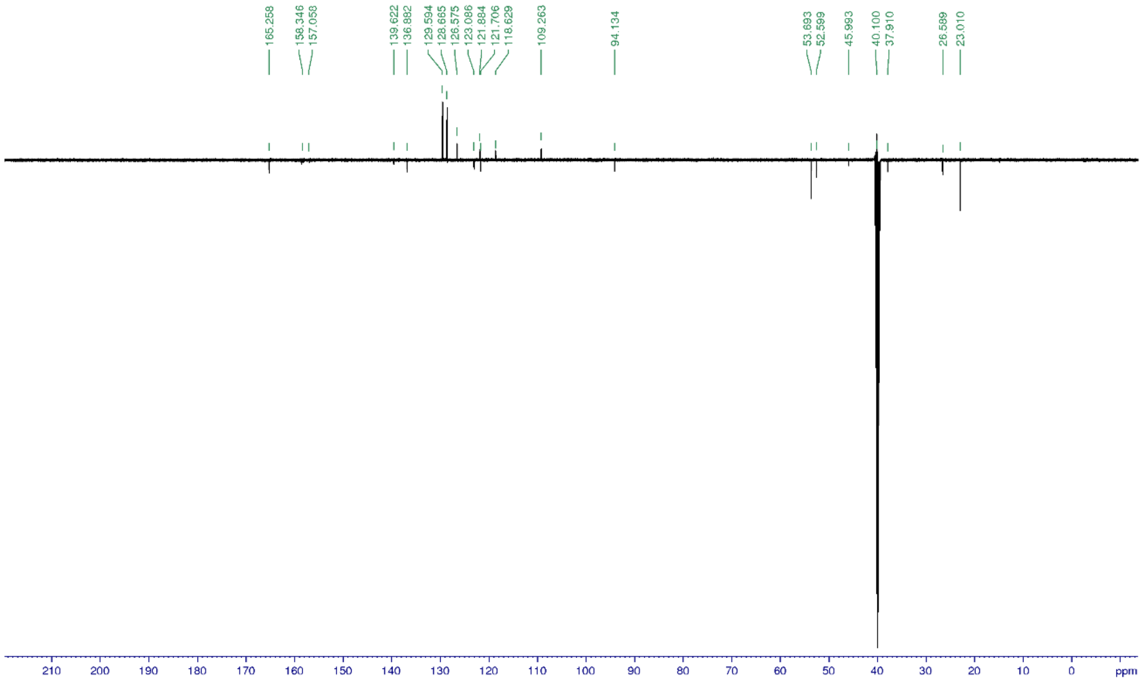


*2-Benzyl-7-(2-methyltetrazol-5-yl)-N-[3-(1-piperidyl)propyl]-9H-pyrimido[4,5-b]indol-4-amine (**12**)*

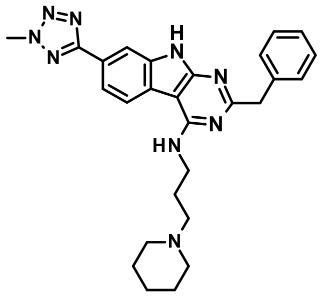

Title compound was synthesized according to the *General procedure H1* using 2-benzyl-4-chloro-7-(2-methyltetrazol-5-yl)-9H-pyrimido[4,5-b]indole (**40**, 11 mg, 0.03 mmol), 3-(1-piperidyl)propan-1-amine (3.5 eq.) and DIPEA (15.0 eq.) to give the desired product (4 mg, 26%).

1H NMR (500 MHz, DMSO-$d_6$): δ ppm 12.16 (s, 1 H), 8.96 (brs, 1 H), 8.42 (d, J = 8.0 Hz, 1 H), 8.10 (d, J = 0.9 Hz, 1 H), 7.93 (dd, J = 8.3, 1.5 Hz, 1 H), 7.46 (t, J = 5.2 Hz, 1 H), 7.38 (d, J = 7.2 Hz, 2 H), 7.30 (t, J = 7.6 Hz, 2 H), 7.21 (t, J = 7.6 Hz, 1 H), 4.44 (s, 3 H), 4.08 (s, 2 H), 3.69 (q, J = 6.5 Hz, 2 H), 3.39/2.81 (d+dd, J = 12.1/21.1, 9.5 Hz, 4 H), 3.09 (m, 2 H), 2.03 (qn, J = 7.4 Hz, 2 H), 1.79/1.60 (d+m, J = 14.1 Hz, 4 H), 1.68/1.35 (m+m, 2 H); 13C NMR (125 MHz, DMSO-$d_6$): δ 166.2, 165.3, 158.2, 157.1, 139.7, 136.9, 129.6, 128.7, 126.6, 123.1, 121.9, 121.7, 118.6, 109.3, 94.1, 54.6, 52.5, 46.0, 40.0, 38.0, 24.6, 23.1, 21.7; UV/Vis: $\lambda_{max}$ 233 nm; HRMS (m/z): [M+H]+ calcd. for $C_{27}H_{31}N_9$, 482.2780; found, 482.2779.

| tR | Peak type | % | Product type | MW | Ions | Comment |
|---|---|---|---|---|---|---|
| 1.92 | minor | 2.3 | | | | |
| 2.23 | principal | 92.4 | Product | *481.2702* | [M+H]+=482.2779 (δ=0.8 ppm) | |
| 2.27 | minor | 4.1 | By-Product | | [M+H]+ m/z=458.25 | |

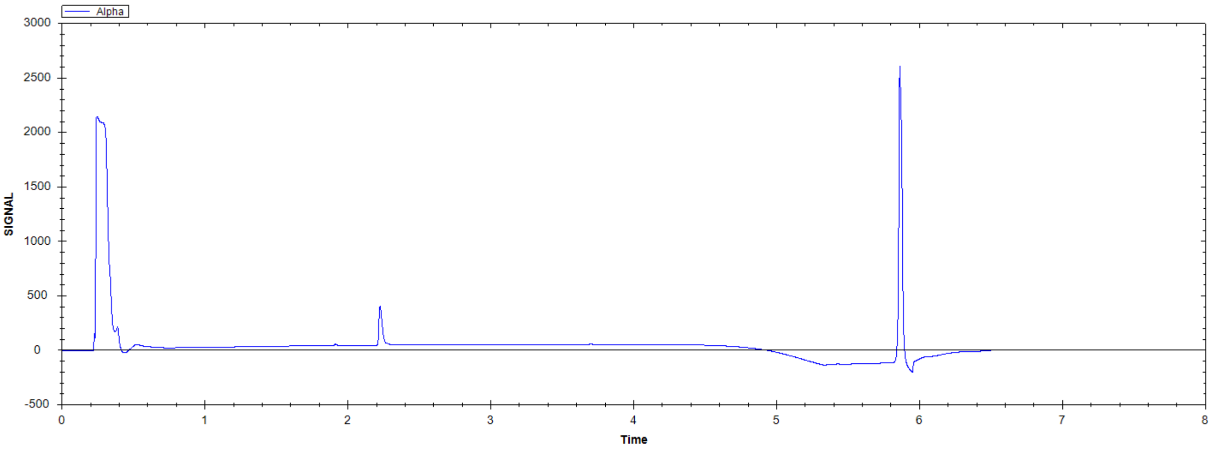

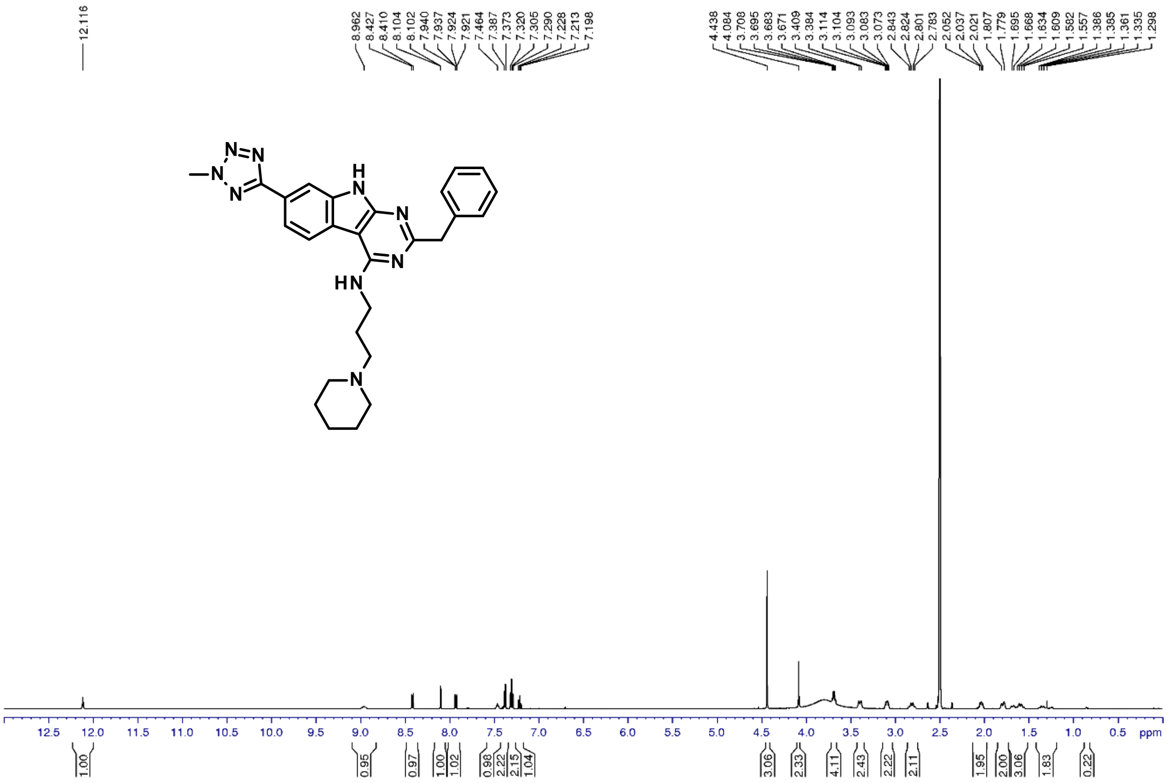
12.116
8.962 8.427 8.410 8.104 8.102 7.940 7.937 7.924 7.921 7.464 7.387 7.373 7.320 7.305 7.290 7.228 7.213 7.198
4.438 4.084 3.708 3.695 3.683 3.671 3.409 3.384 3.114 3.104 3.093 3.083 3.073 2.843 2.824 2.801 2.783 2.052 2.037 2.021 1.807 1.779 1.695 1.668 1.634 1.609 1.582 1.557 1.386 1.385 1.361 1.335 1.298
HN
12.5 12.0 11.5 11.0 10.5 10.0 9.5 9.0 8.5 8.0 7.5 7.0 6.5 6.0 5.5 5.0 4.5 4.0 3.5 3.0 2.5 2.0 1.5 1.0 0.5 ppm
1.00 0.95 0.97 1.00 1.02 0.98 2.22 2.15 1.04 3.06 2.33 4.11 2.43 2.22 2.11 1.95 2.00 3.06 1.83 0.22

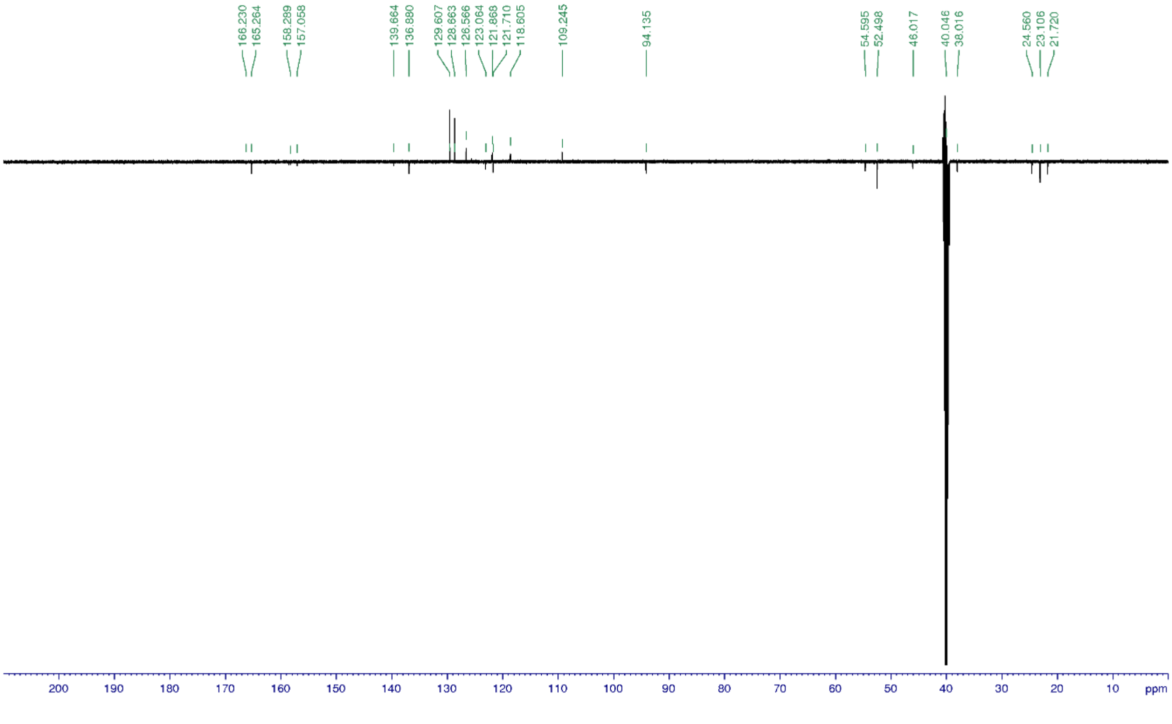
166.230 165.264 158.289 157.058 139.664 136.880 129.607 128.663 126.566 123.064 121.868 121.710 118.605 109.245 94.135 54.595 52.498 48.017 40.046 38.016 24.560 23.106 21.720
200 190 180 170 160 150 140 130 120 110 100 90 80 70 60 50 40 30 20 10 ppm

*Methyl 2-benzyl-4-(3-pyrrolidin-1-ylpropylamino)-9H-pyrimido[4,5-b]indole-7-carboxylate (**13**)*

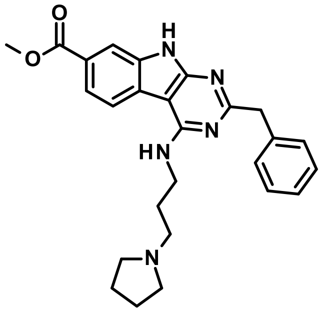

Title compound was synthesized according to the *General procedure H1* using methyl 2-benzyl-4-chloro-9H-pyrimido[4,5-b]indole-7-carboxylate (**29**, 35 mg, 0.08 mmol), 3-pyrrolidin-1-ylpropan-1-amine (3.5 eq.) and DIPEA (15.0 eq.) to give the desired product (15 mg, 42%).

1H NMR (500 MHz, DMSO-$d_6$): δ ppm 12.17 (s, 1 H), 9.46 (brs, 1 H), 8.37 (d, J = 8.3 Hz, 1 H), 8.01 (d, J = 1.5 Hz, 1 H), 7.84 (dd, J = 8.3, 1.5 Hz, 1 H), 7.53 (t, J = 5.2 Hz, 1 H), 7.37 (d, J = 7.3 Hz, 2 H), 7.30 (t, J = 7.3 Hz, 2 H), 7.21 (t, J = 7.3 Hz, 1 H), 4.08 (s, 2 H), 3.88 (s, 3 H), 3.69 (q, J = 6.1 Hz, 2 H), 3.51 (brm, 2 H), 3.16 (m, 2 H), 2.90 (brm, 2 H), 2.01 (qn, J = 6.5 Hz, 2 H), 1.98/1.83 (brm+brm, 4 H); 13C NMR (125 MHz, DMSO-$d_6$): δ 167.1, 166.8, 158.5, 157.3, 139.6, 136.2, 129.6, 128.7, 126.6, 125.4, 123.9, 121.2, 121.0, 112.6, 94.1, 53.7, 52.6, 52.6, 46.0, 37.9, 26.6, 23.0; UV/Vis: $\lambda_{max}$ 227 nm; HRMS (m/z): [M+H]+ calcd. for $C_{26}H_{29}N_5O_2$, 444.2399; found, 444.2397.

| tR | Peak type | % | Product type | MW | Ions | Comment |
|---|---|---|---|---|---|---|
| 1.87 | minor | **2.5** | | | | |
| 1.92 | principal | **94.1** | **Product** | ***443.2321*** | [M+H]+=444.2397 (δ=0.7 ppm) | |
| 1.96 | minor | **3.0** | | | m/z=457.2717 | |

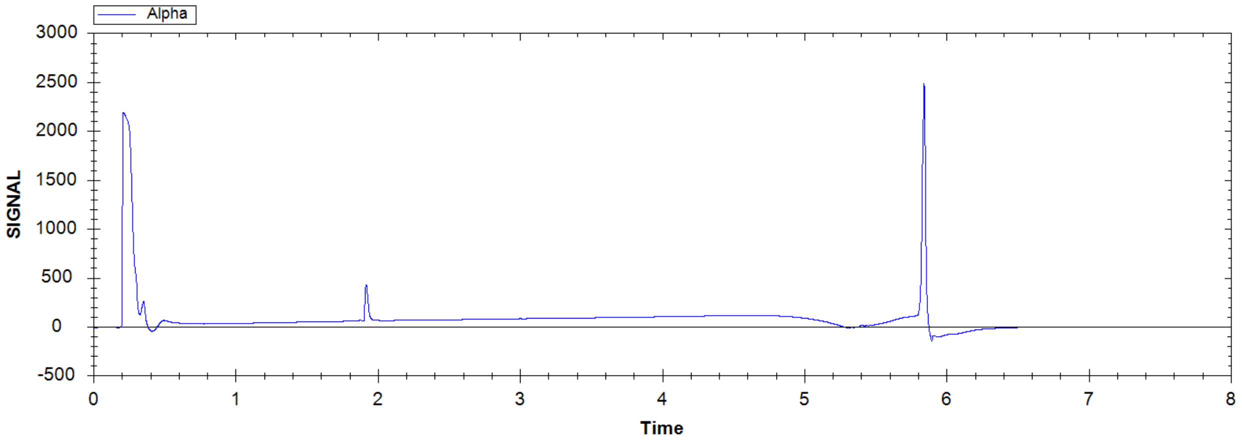

methyl 2-benzyl-4-(3-pyrrolidin-1-ylpropylamino)-9H-pyrimido[4,5-b]indole-7-carboxylate

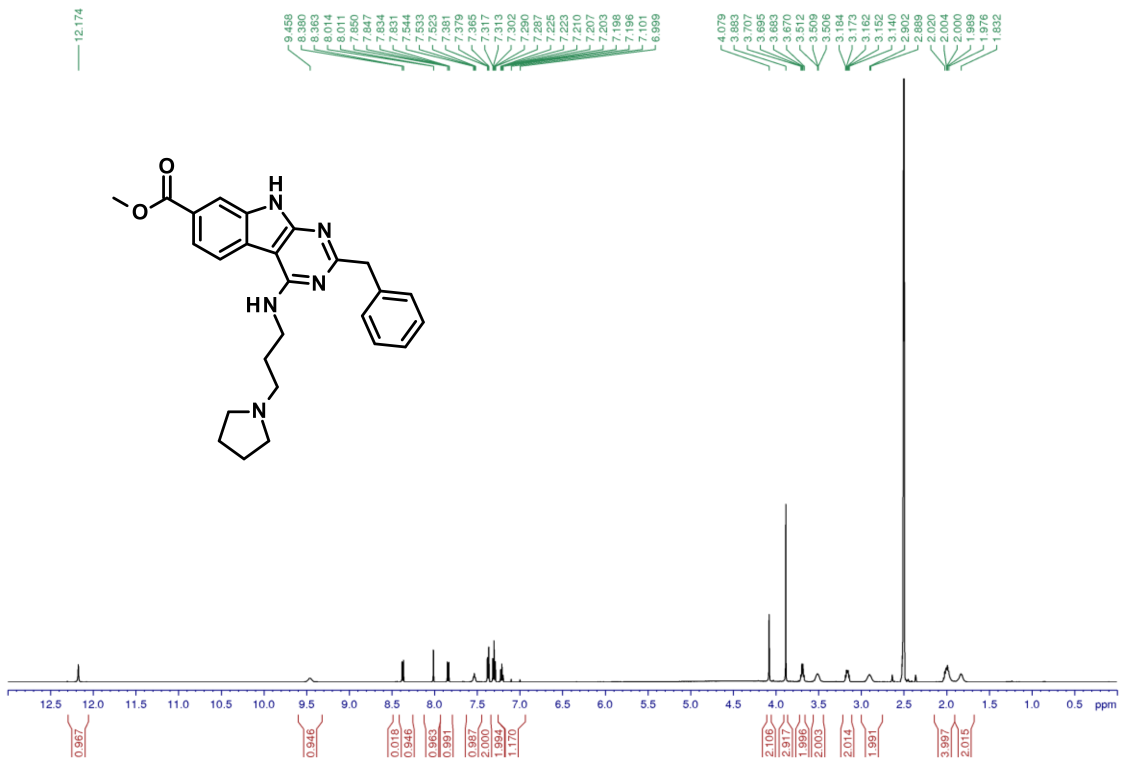


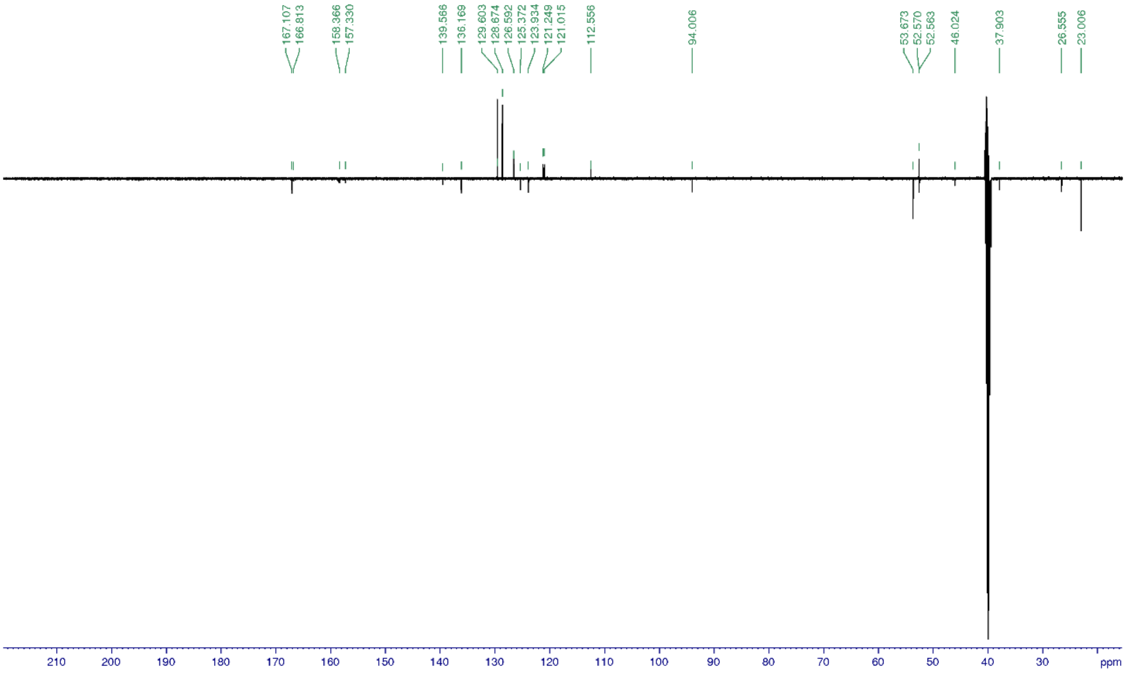


*Methyl 2-benzyl-4-[3-(1-piperidyl)propylamino]-9H-pyrimido[4,5-b]indole-7-carboxylate (**14**)*

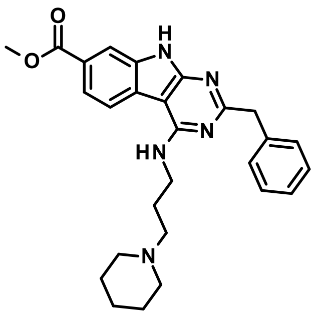

Title compound was synthesized according to the *General procedure H1* using methyl 2-benzyl-4-chloro-9H-pyrimido[4,5-b]indole-7-carboxylate (**29**, 40 mg, 0.09 mmol), 3-(1-piperidyl)propan-1-amine (3.5 eq.) and DIPEA (15.0 eq.) to give the desired product (14 mg, 34%).

1H NMR (500 MHz, DMSO-$d_6$): δ ppm 12.16 (s, 1 H), 8.37 (d, J = 7.8 Hz, 1 H), 8.01 (d, J = 1.3 Hz, 1 H), 7.84 (m, 1 H), 7.53 (t, J = 5.8 Hz, 1H), 7.37 (d, J = 7.6 Hz, 2 H), 7.30 (t, J = 7.5 Hz, 2 H), 7.21 (m, 1 H), 4.08 (s, 2 H), 3.88 (s, 3 H), 3.68 (q, J = 6.2 Hz, 2 H), 3.39 (m, 2 H), 3.08 (m, 2 H), 2.81 (m, 2 H), 2.03 (m, 2 H), 1.79 (m, 2 H), 1.67 (m, 1 H), 1.59 (m, 2 H), 1.34 (m, 1 H); 13C NMR (125 MHz, DMSO-$d_6$): δ 167.1, 166.8, 157.6, 139.6, 136.1, 129.6, 128.7, 126.6, 125.4, 123.9, 121.3, 121.0, 112.6, 94.0, 54.6, 52.6, 52.5, 52.5, 46.1, 38.0, 24.5, 23.1, 21.7; UV/Vis: $\lambda_{max}$ 226 nm; HRMS (m/z): [M+H]+ calcd. for $C_{27}H_{31}N_5O_2$, 458.2556; found, 458.2553.

| tR | Peak type | % | Product type | MW | Ions | Comment |
|---|---|---|---|---|---|---|
| 2.24 | secondary | **19.0** | **By-Product** | | [M+H]+ m/z=425.24 | |
| 2.30 | principal | **79.0** | **Product** | ***457.2478*** | [M+H]+=458.2553 (δ=0.5 ppm) | |

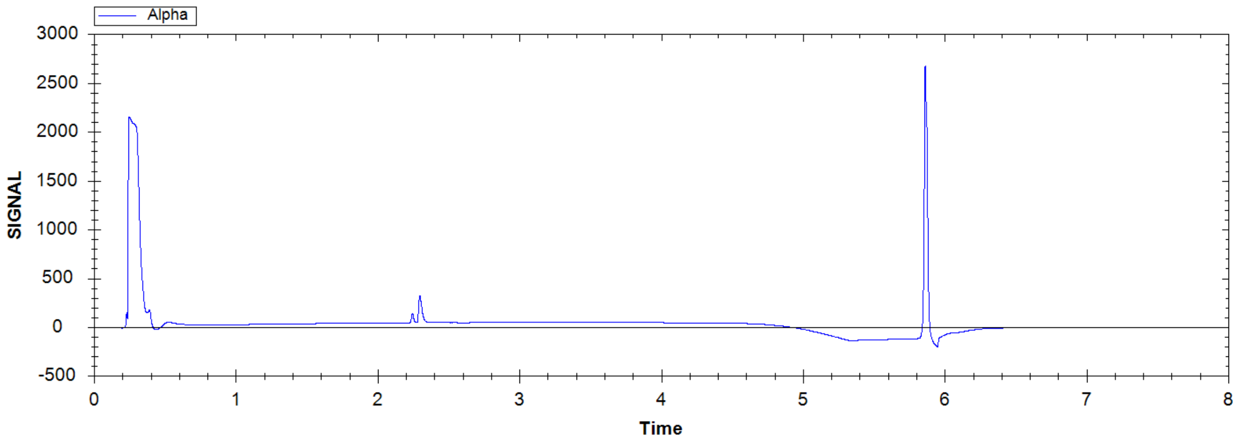

methyl 2-benzyl-4-[3-(1-piperidyl)propylamino]-9H-pyrimido[4,5-b]indole-7-carboxylate

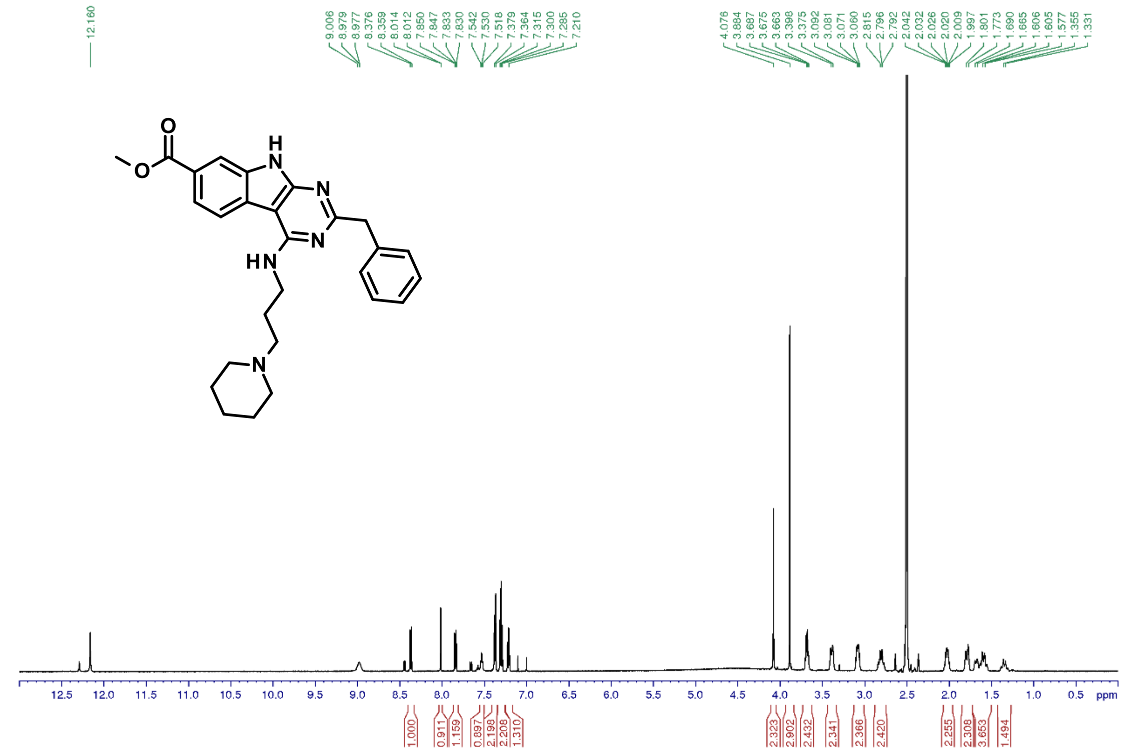


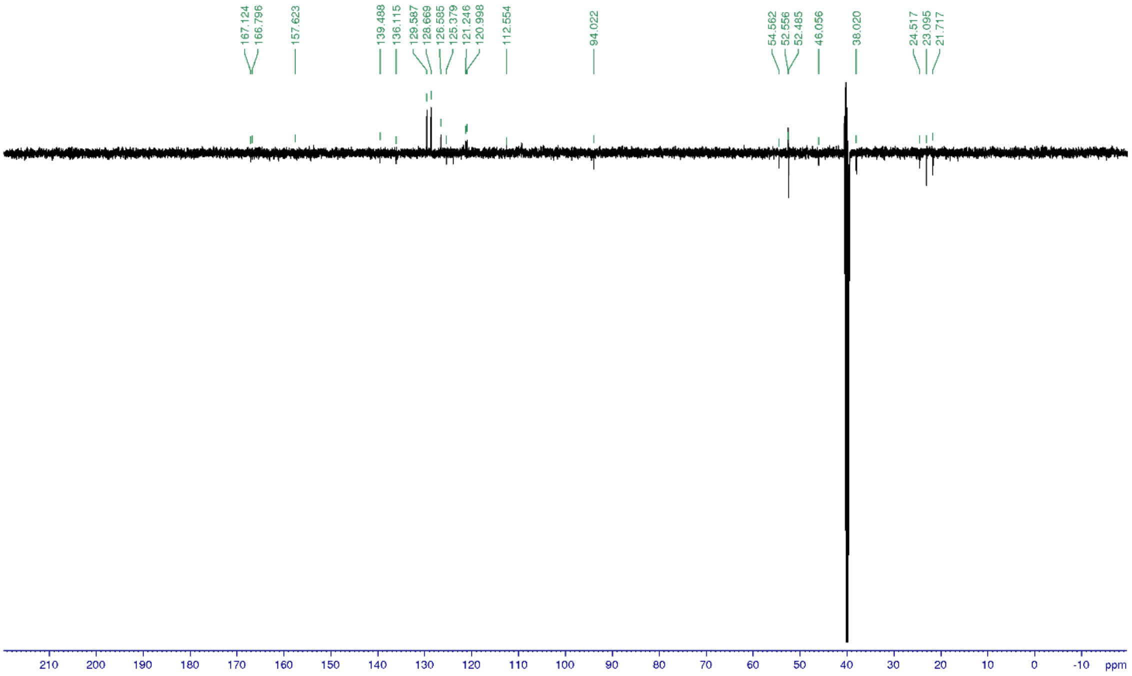


*N4-(4-aminocyclohexyl)-2-benzyl-N6-[3-(2-methyltetrazol-5-yl)phenyl]pyrimidine-4,6-diamine (**15**)*

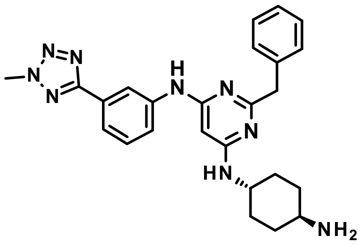


Title compound was synthesized in a two-step procedure according to the *General procedure L2* using 2-benzyl-6-chloro-N-[3-(2-methyltetrazol-5-yl)phenyl]pyrimidin-4-amine (**44**, 40 mg, 0.10 mmol) to give tert-butyl N-[4-[[2-benzyl-6-[3-(2-methyltetrazol-5-yl)anilino]pyrimidin-4-yl]amino]cyclohexyl]carbamate intermediate, which was not characterized, it was used without purification as starting material according to *General procedure I* to give the desired product (3 mg, 6%).

1H NMR (500 MHz, DMSO-$d_6$): δ ppm 9.10 (brs, 1 H), 8.39 (brs, 1 H), 7.68 (d, J = 8.1 Hz, 1 H), 7.55 (d, J = 7.9 Hz, 1 H), 7.40 (d, J = 8.0 Hz, 2 H), 7.36 (t, J = 7.8 Hz, 1 H), 7.27 (t, J = 7.5 Hz, 2 H), 7.13 (t, J = 7.5 Hz, 1 H), 6.70 (d, J = 8.1 Hz, 1 H), 5.61 (s, 1 H), 4.43 (s, 3 H), 3.82 (s, 2 H), 3.17 (m, 1 H), 2.5 (m, 1 H), 1.94-1.02 (m, 8 H); 13C NMR (125 MHz, DMSO-$d_6$): δ 129.9, 121.1, 119.0, 117.0, 50.3, 49.5, 46.1, 40.1; UV/Vis: $\lambda_{max}$ 245 nm; HRMS (m/z): [M+H]+ calcd. for $C_{25}H_{29}N_9$, 456.2624; found, 456.2620.

| tR | Peak type | % | Product type | MW | Ions | Comment |
|---|---|---|---|---|---|---|
| 2.00 | principal | >99 | **Product** | ***455.2546*** | [M+H]+=456.262 (δ=0.3 ppm) | |

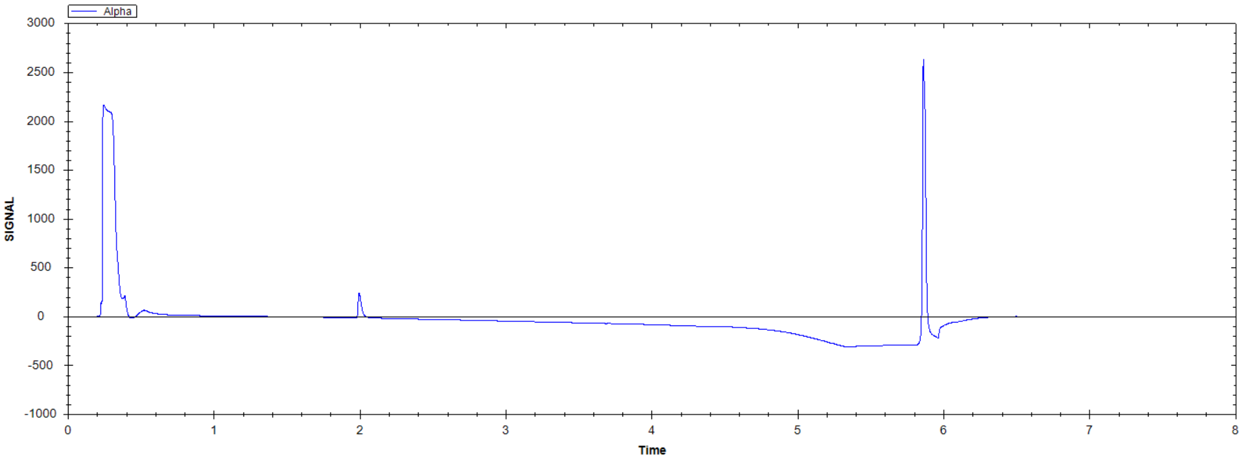

N4-(4-aminocyclohexyl)-2-benzyl-N6-[3-(2-methyltetrazol-5-yl)phenyl]pyrimidine-4,6-diamine

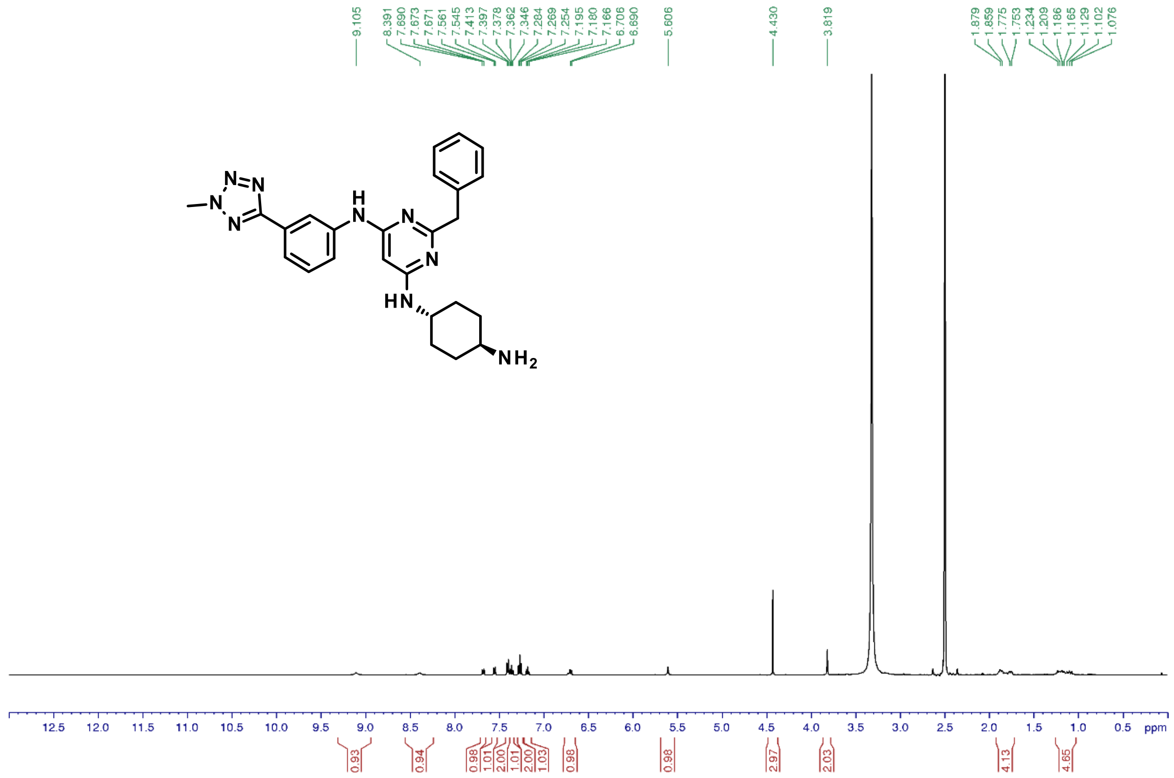

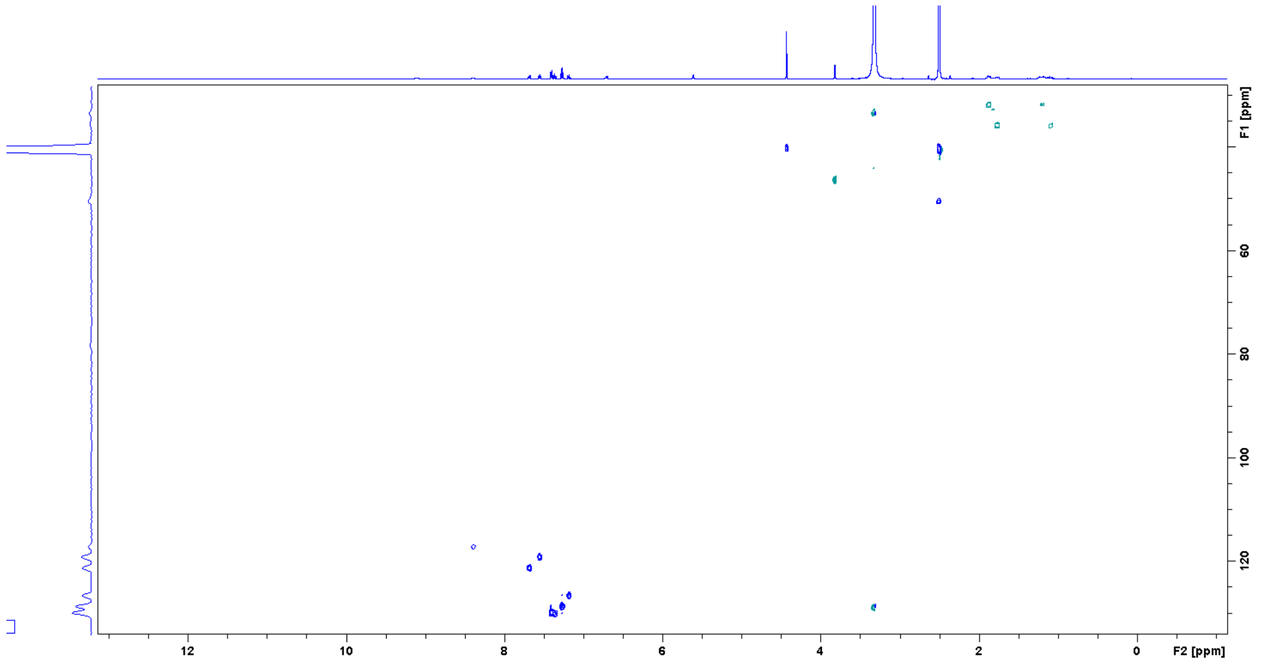

*N-(5-bromo-2-iodo-phenyl)-2,2,2-trifluoro-acetamide (**16**)*

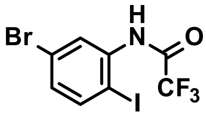

Title compound was synthesized according to the *General procedure B1* using 5-bromo-2-iodo-aniline (8.5 g, 29 mmol) and TFA anhydride (1.1 eq., 4.4 mL) to give the desired product (10.9 g, 98%).

1H NMR (500 MHz, DMSO-$d_6$): δ ppm 11.36 (s, 1 H), 7.89 (d, J = 8.4 Hz, 1 H), 7.68 (d, J = 2.3 Hz, 1 H), 7.36 (dd, J = 8.5, 2.3 Hz, 1 H); 13C NMR (125 MHz, DMSO-$d_6$): δ 155.9, 141.2, 139.2, 133.0, 131.7, 121.9, 98.0; UV/Vis: $\lambda_{max}$ 198 nm; HRMS (m/z): [M+H]+ calcd. for $C_8H_4NOF_3BrI$, 392.8467; found, 392.8475.

N-(5-bromo-2-iodo-phenyl)-2,2,2-trifluoro-acetamide

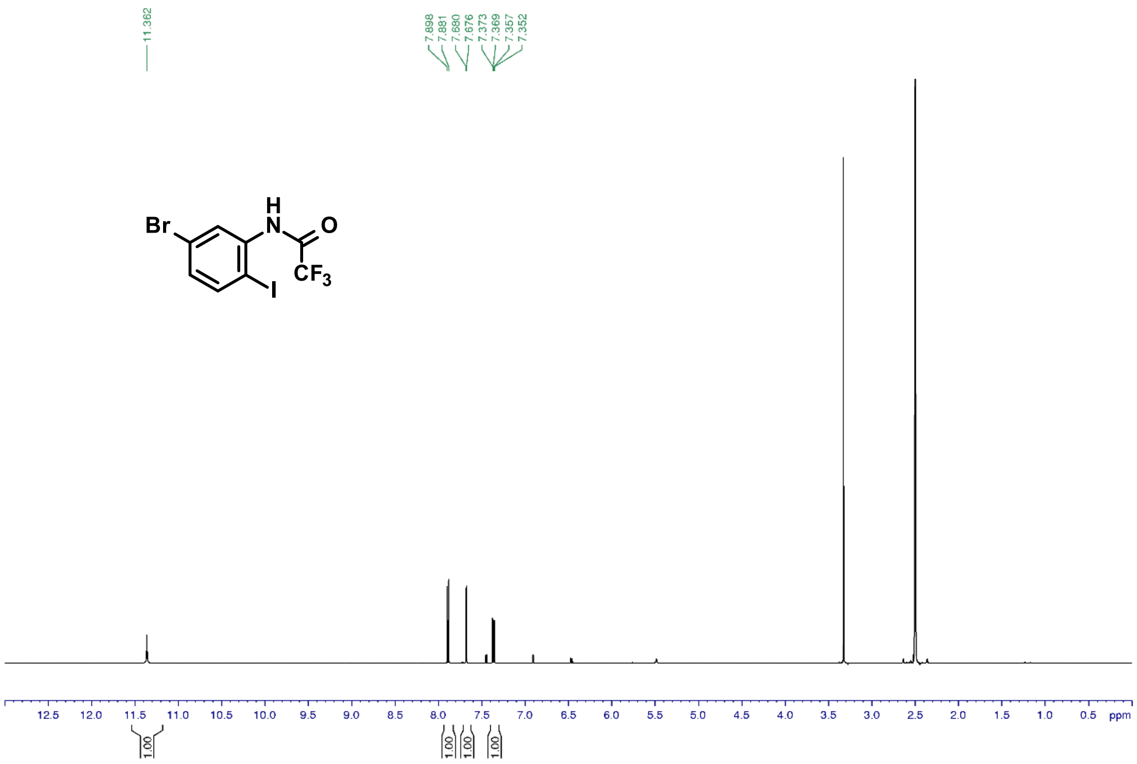

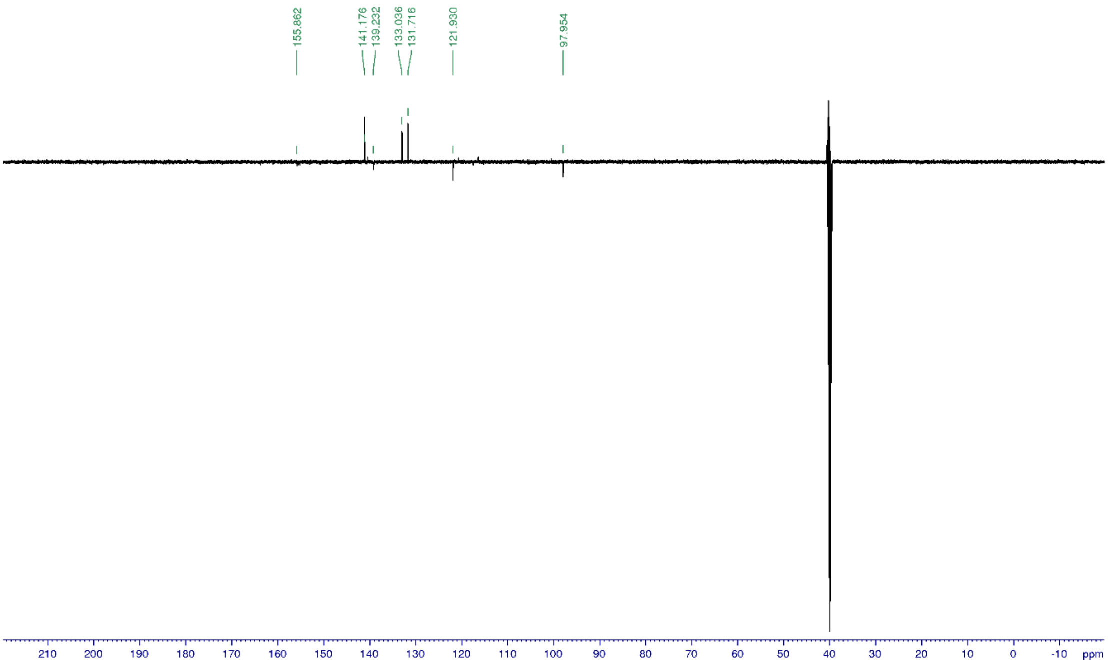


*2-Amino-6-bromo-1H-indole-3-carbonitrile (**17**)*

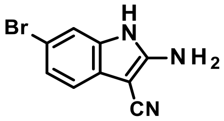

Title compound was synthesized according to the *General procedure C2* using N-(5-bromo-2-iodo-phenyl)-2,2,2-trifluoro-acetamide (**16**, 5.5 g, 14 mmol) to give the desired product (2.7 g, 80%).

1H NMR (500 MHz, DMSO-$d_6$): δ ppm 10.78 (s, 1 H), 7.29 (d, J = 1.5 Hz, 1 H), 7.09 (dd, J = 8.2, 1.7 Hz 1 H), 7.06 (d, J = 8.2 Hz, 1 H), 6.93 (s, 2 H); 13C NMR (125 MHz, DMSO-$d_6$): δ 154.8, 133.7, 128.0, 123.6, 117.7, 117.0, 113.3, 111.7, 62.1; UV/Vis: $\lambda_{max}$ 226 nm; HRMS (m/z): [M+Na]+ calcd. for $C_9H_6BrN_3$, 257.9642; found, 257.9638.

2-Amino-6-bromo-1H-indole-3-carbonitrile

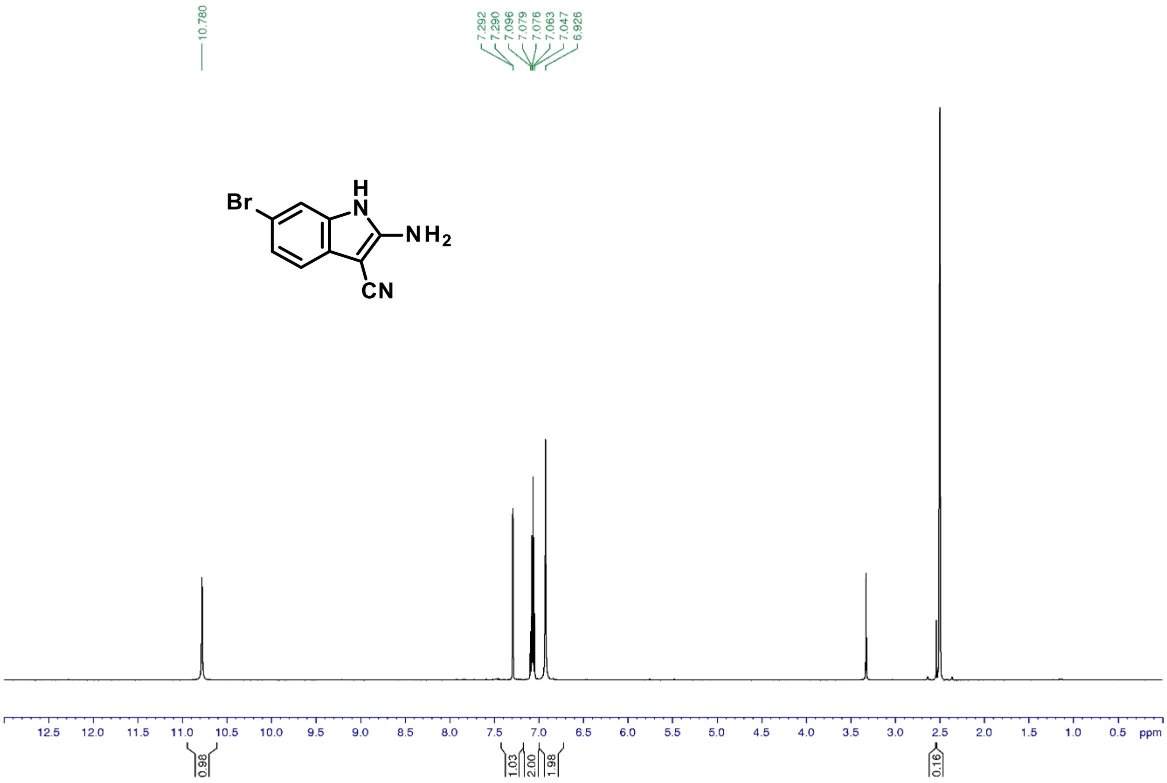


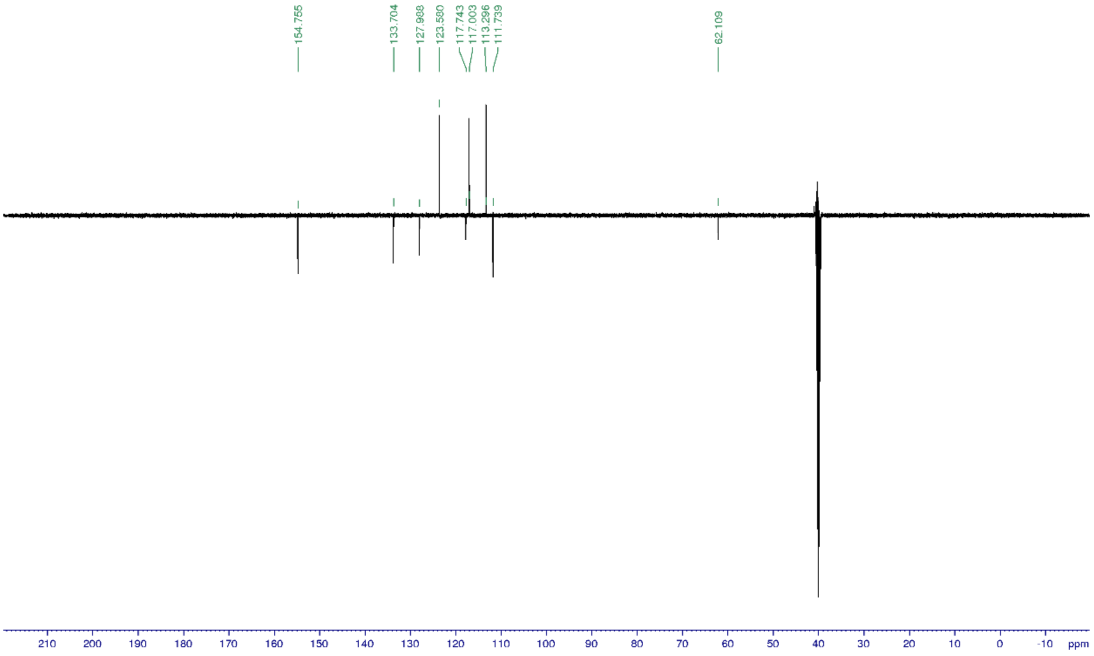


### *2-Amino-6-bromo-1H-indole-3-carboxamide (**18**)*

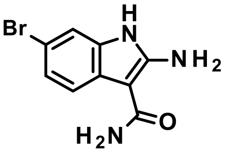

Title compound was synthesized according to the *General procedure N* using 2-amino-6-bromo-1H-indole-3-carbonitrile (**17**, 4.3 g, 18 mmol) to give the desired product (3.4 g, 73%).

1H NMR (500 MHz, DMSO-$d_6$): δ ppm 10.64 (s, 1 H), 7.48 (d, J = 8.3 Hz, 1 H), 7.26 (d, J = 1.8 Hz, 1 H), 7.03 (dd, J = 8.3, 2.0 Hz,1 H), 6.9 (s, 2 H), 6.51 (s, 2 H); 13C NMR (125 MHz, DMSO-$d_6$): δ 168.7, 153.7, 134.0, 125.3, 122.7, 118.4, 112.6, 110.7, 86.5; UV/Vis: $\lambda_{max}$ 234 nm; HRMS (m/z): [M+H]+ calcd. for $C_9H_8BrN_3O$, 253.9928; found, 253.9925.

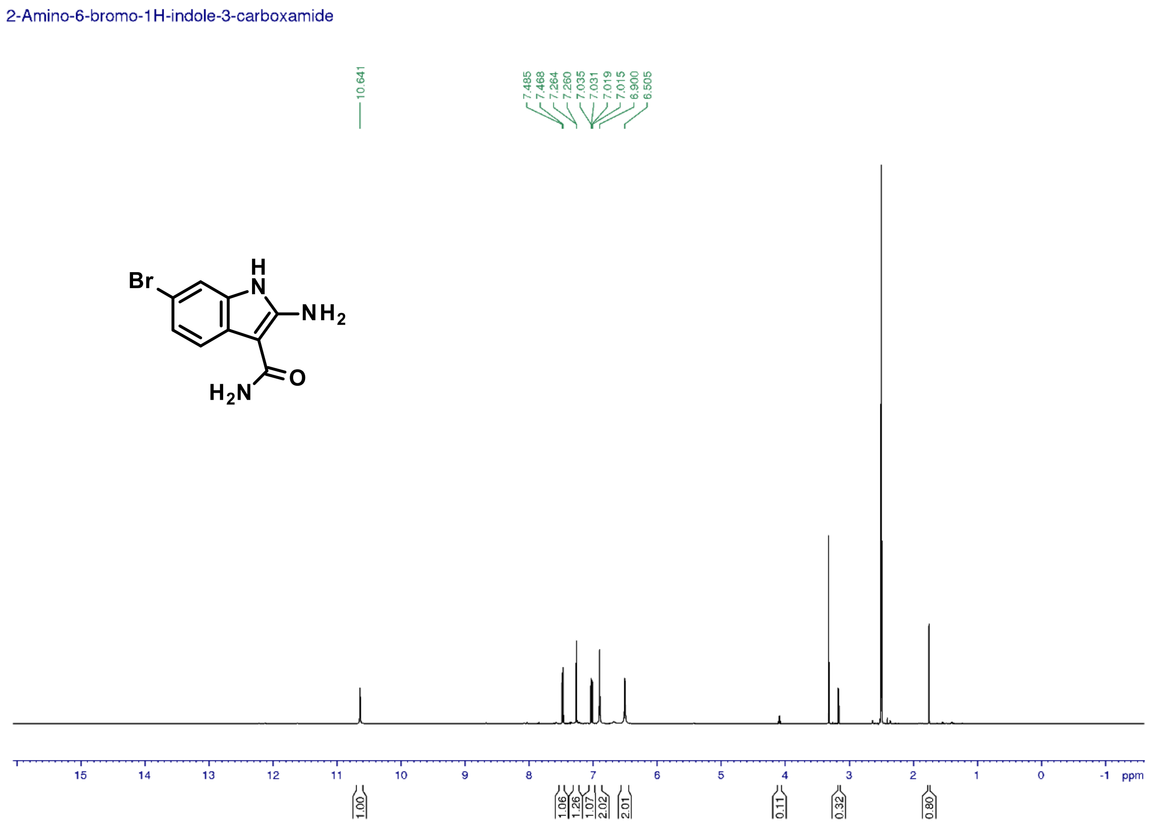

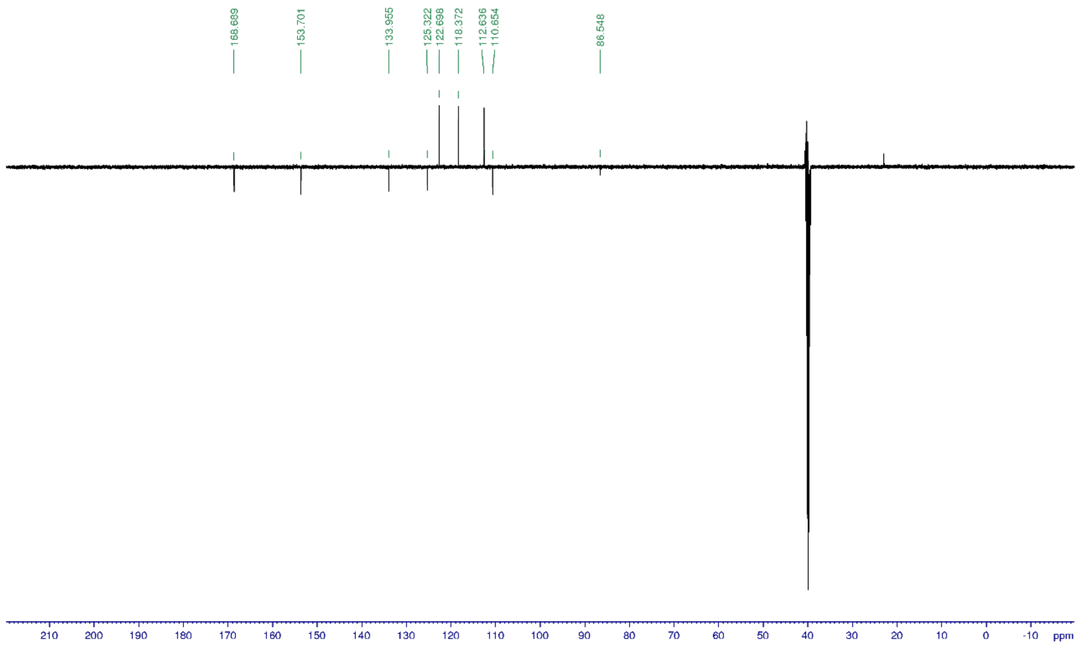


*2-Benzyl-7-bromo-9H-pyrimido[4,5-b]indol-4-ol (**19**)*

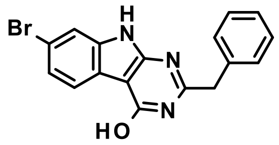


Title compound was synthesized according to the *General procedure D3* using 2-amino-6-bromo-1H-indole-3-carboxamide (**18**, 1.1 g, 4.3 mmol) to give the desired product (1.1 g, 73%).

1H NMR (500 MHz, DMSO-$d_6$): δ ppm 12.37 (brs, 2 H), 7.86 (d, J = 8.2 Hz, 1 H), 7.56 (d, J = 1.7 Hz, 1 H), 7.38 (d, J = 8.3, 1.4 Hz, 2 H), 7.35 (dd, J = 6.4, 1.9 Hz, 1 H), 7.33 (t, J = 6.3 Hz, 2 H), 7.25 (t, J = 7.3 Hz, 1 H), 4.01 (s, 2 H); 13C NMR (125 MHz, DMSO-$d_6$): δ 160.1, 159.3, 155.2, 137.0, 129.5, 129.0, 127.3, 124.3, 122.3, 121.6, 116.5, 114.6, 98.3, 41.0; UV/Vis: $\lambda_{max}$ 244 nm; HRMS (m/z): [M+H]+ calcd. for $C_{17}H_{12}BrN_3O$, 354.0241; found, 354.0237.

2-Benzyl-7-bromo-9H-pyrimido[4,5-b]indol-4-ol

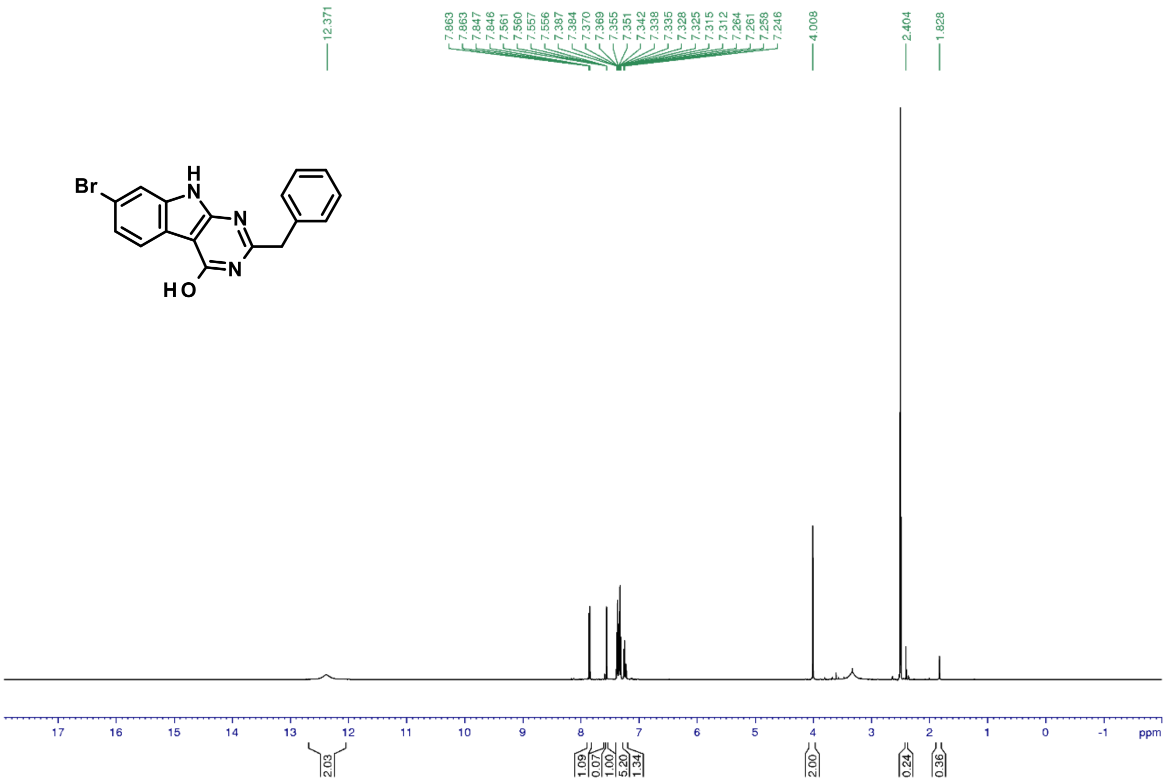


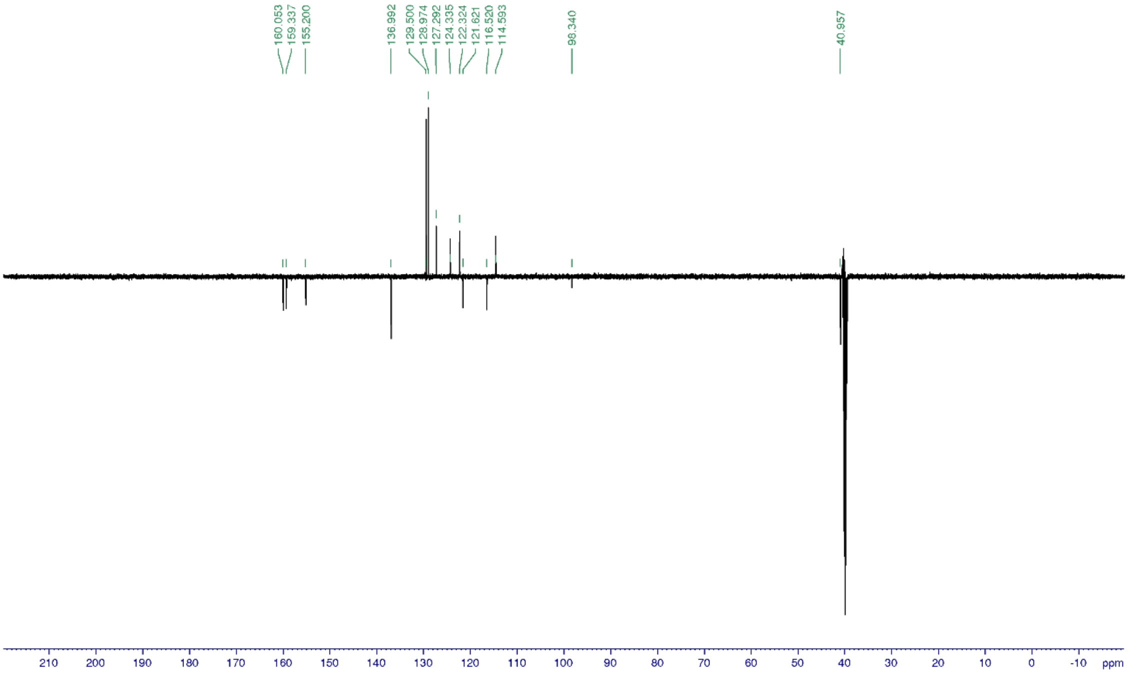


*Tert-butyl N-[4-[(2-benzyl-7-bromo-9H-pyrimido[4,5-b]indol-4-yl)amino]cyclohexyl]carbamate (**20**)*

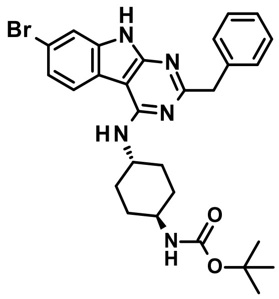

Title compound was synthesized according to the *General procedure H2* using 2-benzyl-7-bromo-9H-pyrimido[4,5-b]indol-4-ol (**19**, 30 mg, 0.08 mmol) to give the desired product (17 mg, 36%).

1H NMR (500 MHz, DMSO-d$_6$): δ ppm 11.88 (s, 1 H), 8.26 (d, J = 8.4 Hz, 1 H), 7.52 (d, J = 1.9 Hz, 1 H), 7.37 (d, J = 7.6 Hz, 2 H), 7.35 (dd, J = 8.3, 2.0 Hz, 1 H), 7.29 (t, J = 7.3 Hz, 2 H), 7.19 (t, J = 7.2 Hz, 1 H), 6.79 (d, J = 8.0 Hz, 1 H), 6.71 (d, J = 8.0 Hz, 1 H), 4.20 (m, 1 H), 4.03 (s, 2 H), 3.27 (m, 1 H), 1.93/1.56 (d+d, J = 10.7/12.7 Hz, 4 H), 1.85/1.32 (d+d, J = 11.4/12.0 Hz, 4 H), 1.41 (s, 9 H); 13C NMR (125 MHz, DMSO-d$_6$): δ 166.3, 157.2, 157.1, 156.4, 139.8, 137.7, 129.7, 128.5, 126.5, 123.3, 122.9, 119.2, 117.0, 113.8, 93.6, 77.9, 49.3, 49.2, 46.1, 32.2, 31.3, 28.8; UV/Vis: $\lambda_{max}$ 220 nm; HRMS (m/z): [M+H]+ calcd. for $C_{28}H_{32}BrN_5O_2$, 550.1817; found, 550.1812.

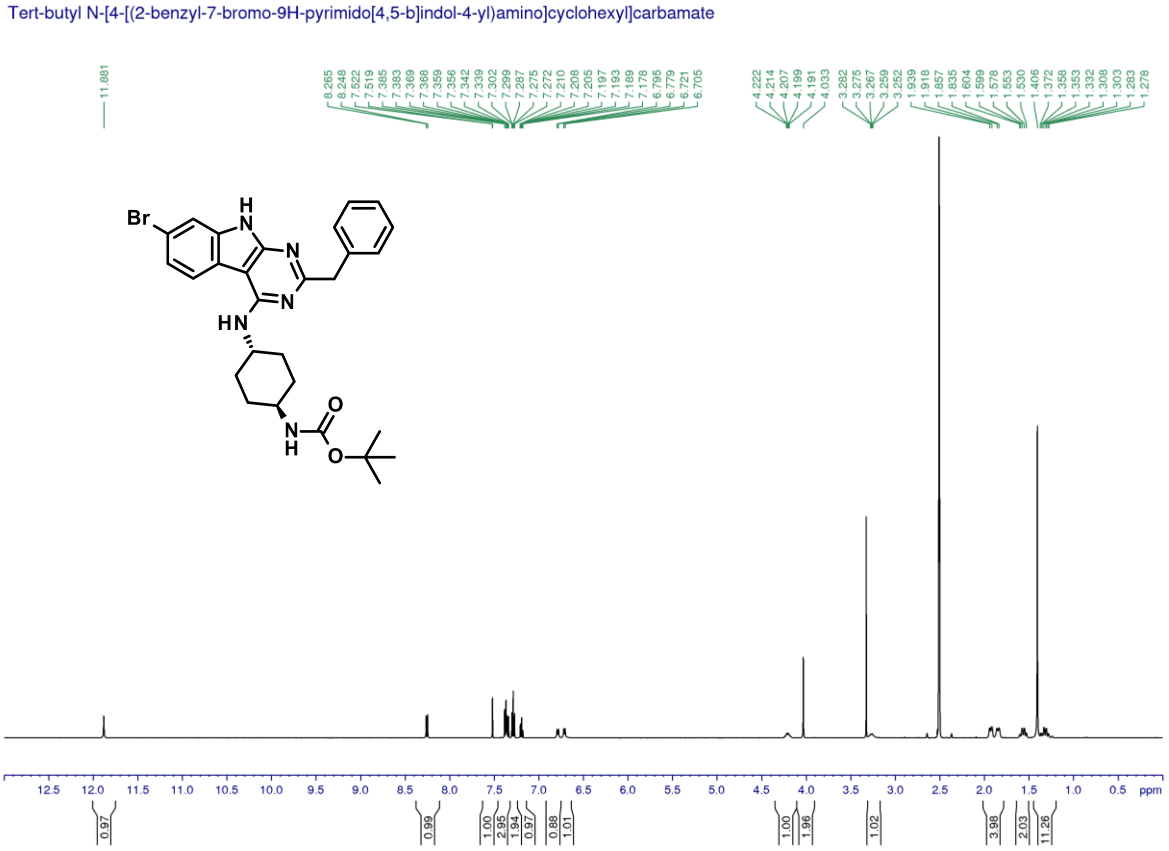

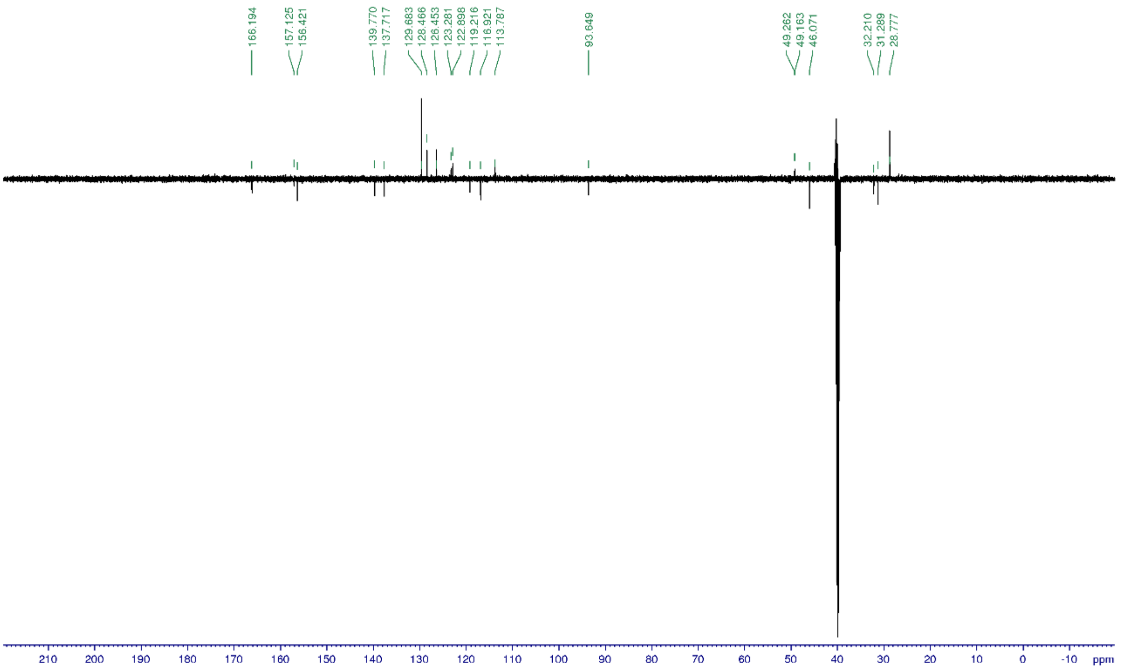


*Tert-butyl N-[4-[[2-benzyl-7-(1-methylpyrazol-4-yl)-9H-pyrimido[4,5-b]indol-4-yl]amino]cyclohexyl]carbamate (**21**)*

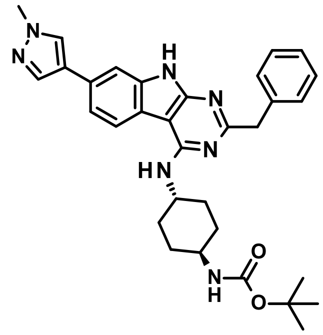

Title compound was synthesized according to the *General procedure M* using tert-butyl N-[4-[(2-benzyl-7-bromo-9H-pyrimido[4,5-b]indol-4-yl)amino]cyclohexyl] carbamate (**20**, 100 mg, 0.18 mmol) and 1-methyl-4-(4,4,5,5-tetramethyl-1,3,2-dioxaborolan-2-yl)pyrazole (5.0 eq., 189 mg) to give the desired product (80 mg, 80%).

1H NMR (500 MHz, DMSO-$d_6$): δ ppm 11.70 (s, 1 H), 8.20 (d, J = 8.2 Hz, 1 H), 8.17 (s, 1 H), 7.88 (d, J = 0.8 Hz, 1 H), 7.48 (d, J = 1.1 Hz, 1 H), 7.40 (dd, J = 8.2, 1.6 Hz, 1 H), 7.38 (d, J = 7.7 Hz, 2 H), 7.28 (t, J = 7.6 Hz, 2 H), 7.19 (t, J = 7.4 Hz, 1 H), 6.78 (d, J = 7.9 Hz, 1 H), 6.55 (d, J = 7.9 Hz, 1 H), 4.21 (m, 1 H), 4.02 (s, 2 H), 3.87 (s, 3 H), 3.27 (m, 1 H), 1.94/1.58 (d+dd, J = 11.2/23.6, 10.4 Hz, 4 H), 1.85/1.32 (d+dd, J = 10.8/23.6, 10.4 Hz, 4 H), 1.40 (s, 9 H); 13C NMR (125 MHz, DMSO-$d_6$): δ 165.3, 157.1, 157.0, 156.2, 140.0, 137.3, 136.4, 129.7, 129.2, 128.5, 128.2, 126.4, 123.2, 122.0, 118.3, 118.0, 107.2, 94.1, 77.9, 49.3, 49.1, 46.1, 39.1, 32.3, 31.4, 28.8; UV/Vis: $\lambda_{max}$ 238 nm; HRMS (m/z): [M+H]+ calcd. for $C_{32}H_{37}N_7O_2$, 552.3086; found, 552.3080.

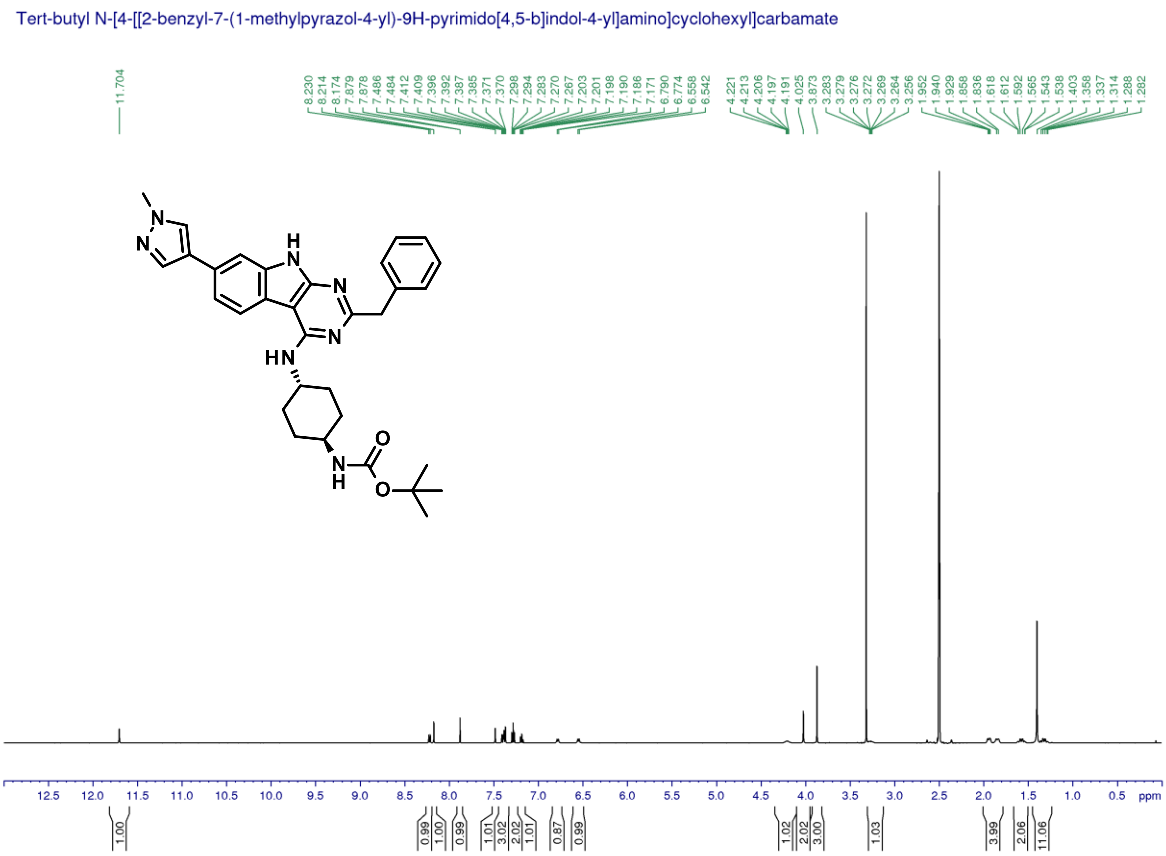


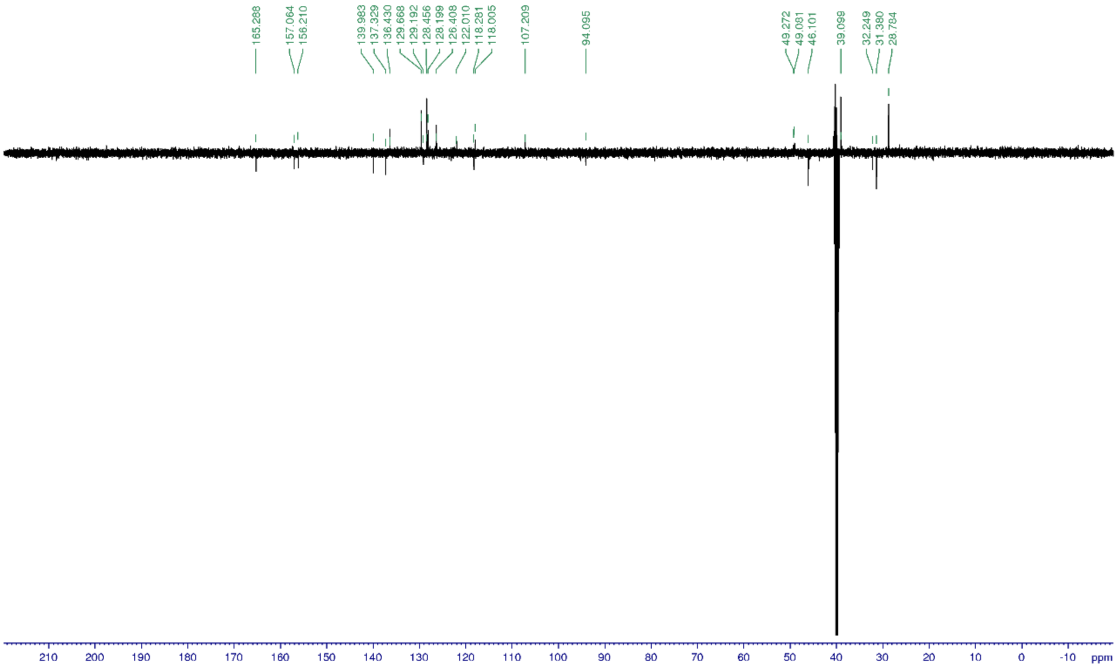

*Tert-butyl N-[4-[[2-benzyl-7-(3,5-dimethylisoxazol-4-yl)-9H-pyrimido[4,5-b]indol-4-yl]amino]cyclohexyl]carbamate (**22**)*

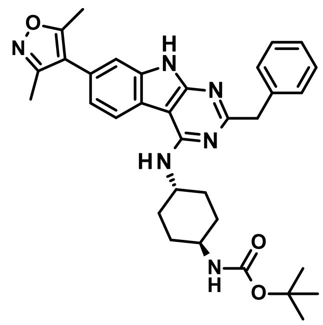

Title compound was synthesized according to the *General procedure M* using tert-butyl N-[4-[(2-benzyl-7-bromo-9H-pyrimido[4,5-b]indol-4-yl)amino]cyclohexyl] carbamate (**20**, 80 mg, 0.15 mmol) and (3,5-dimethylisoxazol-4-yl)boronic acid (1.5 eq., 31 mg) to give the desired product (20 mg, 24%).

1H NMR (500 MHz, DMSO-$d_6$): δ ppm 11.79 (s, 1 H), 8.36 (d, J = 8.4 Hz, 1 H), 7.38 (m, 2 H), 7.32 (d, J = 1.1 Hz,1 H), 7.29 (t, J = 7.6 Hz, 2 H), 7.19 (m, 1 H), 7.18 (dd, J = 7.8, 2.0 Hz, 1 H), 6.79 (br d, J = 8.0 Hz,1 H), 6.66 (br d, J = 7.9 Hz, 1 H), 4.21 (m, 1 H), 4.04 (s, 2 H), 3.27 (m, 1 H), 2.43 (s, 3 H), 2.25 (s, 3 H), 1.94 (m, 2 H), 1.85 (m, 2 H), 1.57 (m, 2 H), 1.40 (s, 9 H), 1.33 (m, 2 H); 13C NMR (125 MHz, DMSO-$d_6$): δ 165.8, 165.3, 158.7, 156.8, 139.8, 136.8, 129.6, 128.5, 126.4, 125.7, 121.9, 121.2, 119.3, 116.9, 111.4, 93.9, 77.9, 49.2, 49.1, 46.1, 32.2, 31.3, 28.7, 11.8, 11.0; UV/Vis: $\lambda_{max}$ 249 nm; HRMS (m/z): [M+H]+ calcd. for $C_{33}H_{38}N_6O_3$, 567.3083; found, 567.3075.

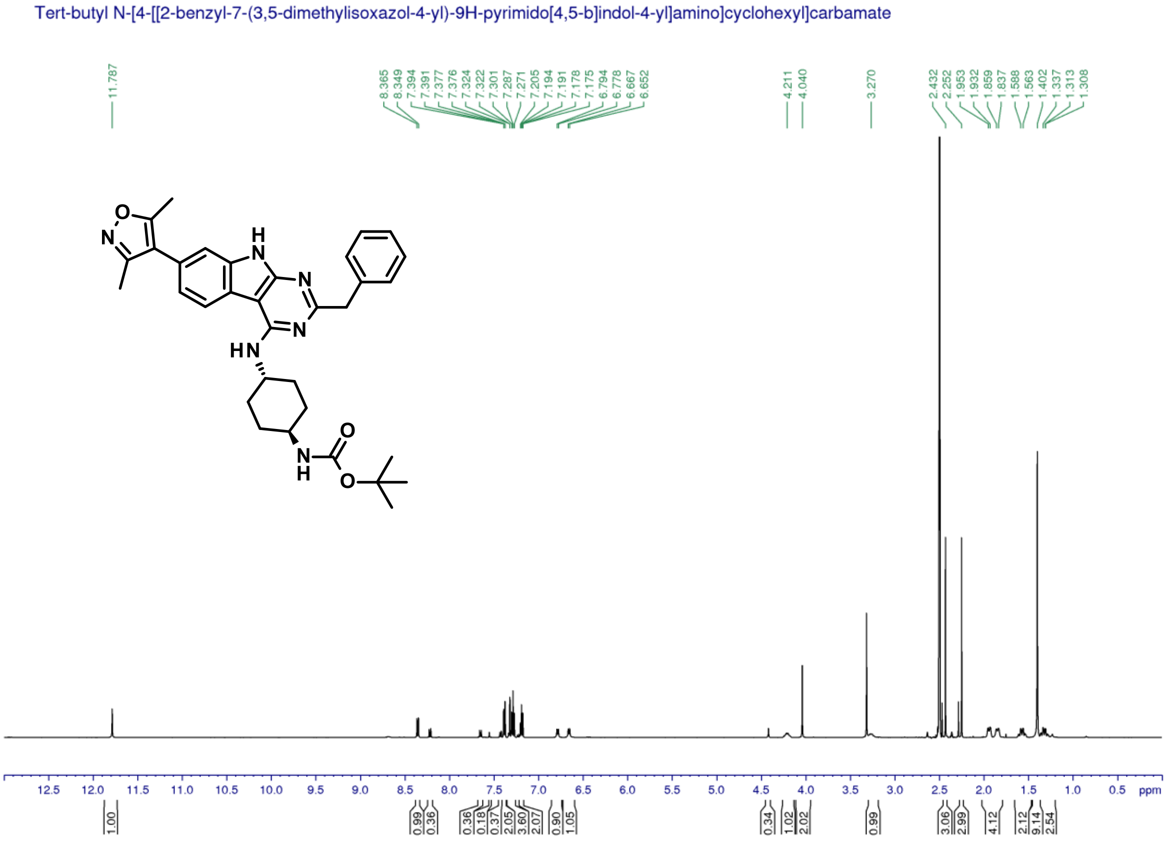

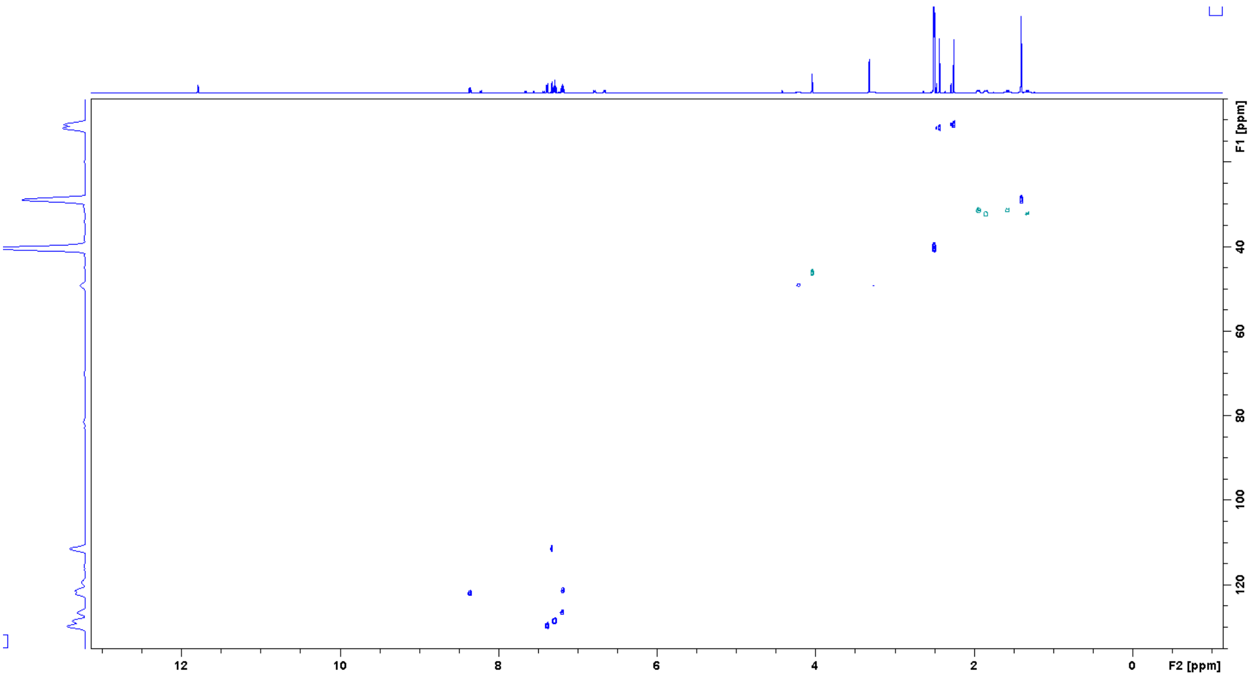


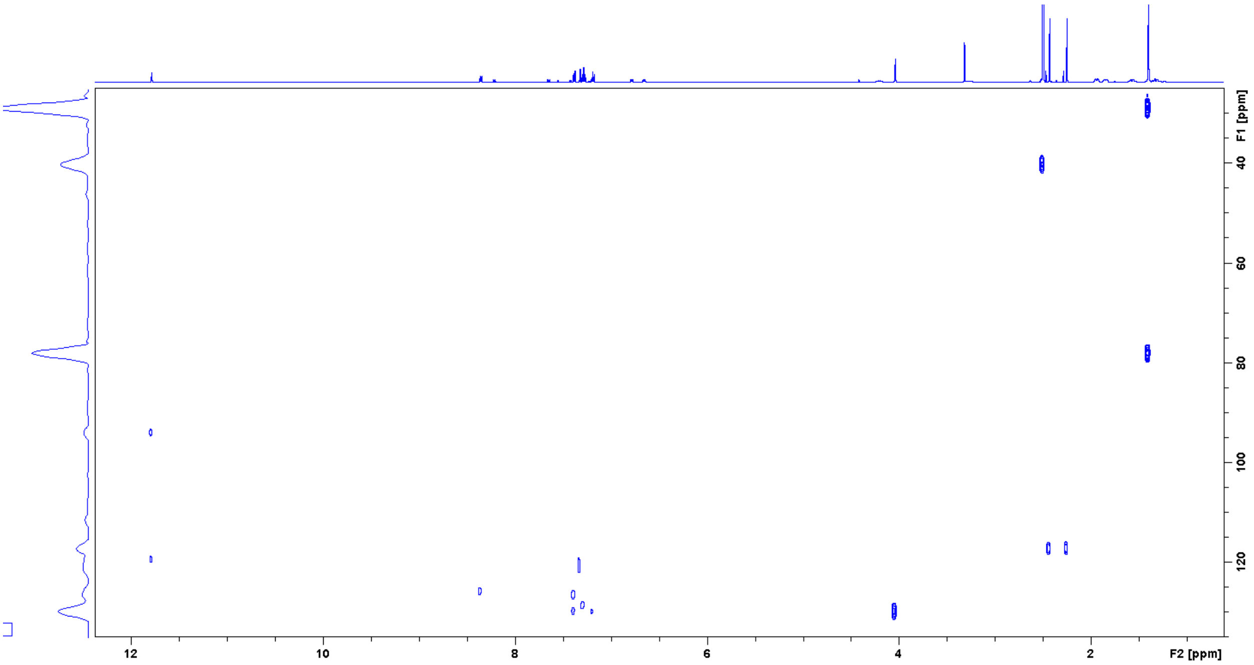


*3-(Hydroxyamino)benzonitrile (**23**)*

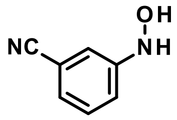


Title compound was synthesized according to the *General procedure A* using 3-nitrobenzonitrile (4.0 g, 27 mmol) to give the desired product (2.1 g, 59%).

1H NMR (500 MHz, DMSO-$d_6$): δ ppm 8.70 (d, J = 1.7 Hz, 1 H), 8.61 (d, J = 2.0 Hz, 1 H), 7.35 (t, J = 7.8 Hz,1 H), 7.15 (dt, J = 7.5, 1.3 Hz, 1 H), 7.11 (dd, J = 2.5, 1.8 Hz, 1 H), 7.09 (dd, J = 8.2, 2.1 Hz,1 H); 13C NMR (125 MHz, DMSO-$d_6$): δ 153.1, 130.3, 122.9, 119.7, 117.8, 115.3, 111.8; UV/Vis: $\lambda_{max}$ 218 nm; HRMS (m/z): [M+H]+ calcd. for $C_7H_6N_2O$, 135.0558; found, 135.0552.

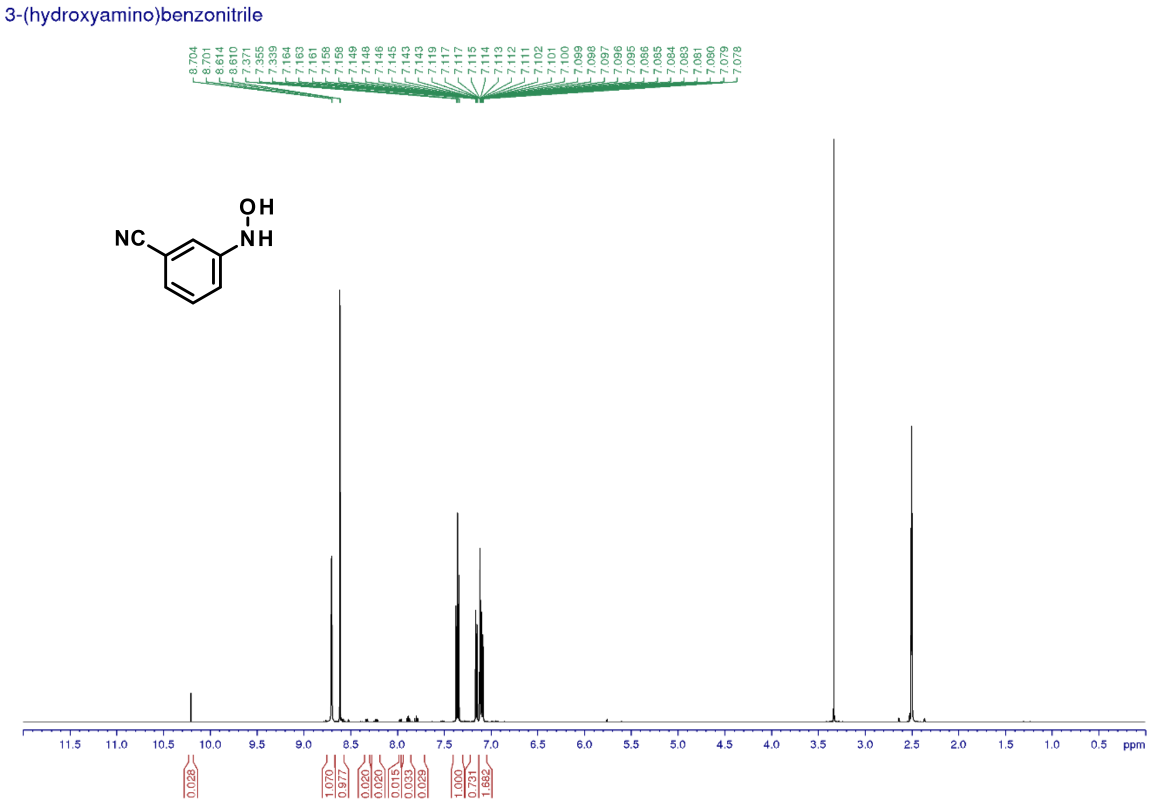


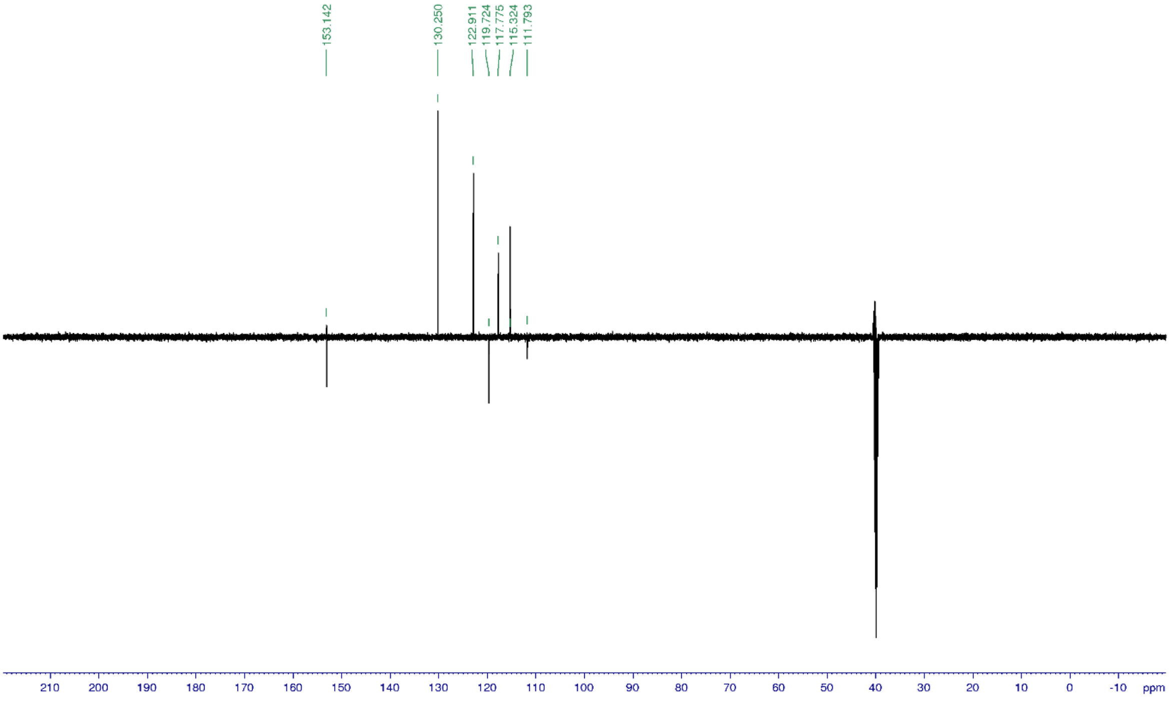


*N-(3-cyanophenyl)-N-hydroxy-acetamide (**24**)*

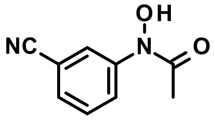

Title compound was synthesized according to the *General procedure B1* using 3-(hydroxyamino)benzonitrile (**23**, 1.5 g, 11 mmol) and acetyl chloride (888 uL, 1.1 eq.) to give the desired product (1.6 g, 82%).

1H NMR (500 MHz, DMSO-$d_6$): δ ppm 10.89 (s, 1 H), 8.07 (m, 1 H), 7.98 (m, 1 H), 7.59 (m, 1 H), 7.59 (m, 1 H), 2.25 (s, 3 H); 13C NMR (125 MHz, DMSO-$d_6$): δ 171.3, 142.6, 130.5, 128.2, 124.2, 122.6, 119.1, 111.8, 23.1; UV/Vis: $\lambda_{max}$ 194 nm; HRMS (m/z): [M+H]+ calcd. for $C_9H_8N_2O_2$, 177.0664; found, 177.0659.

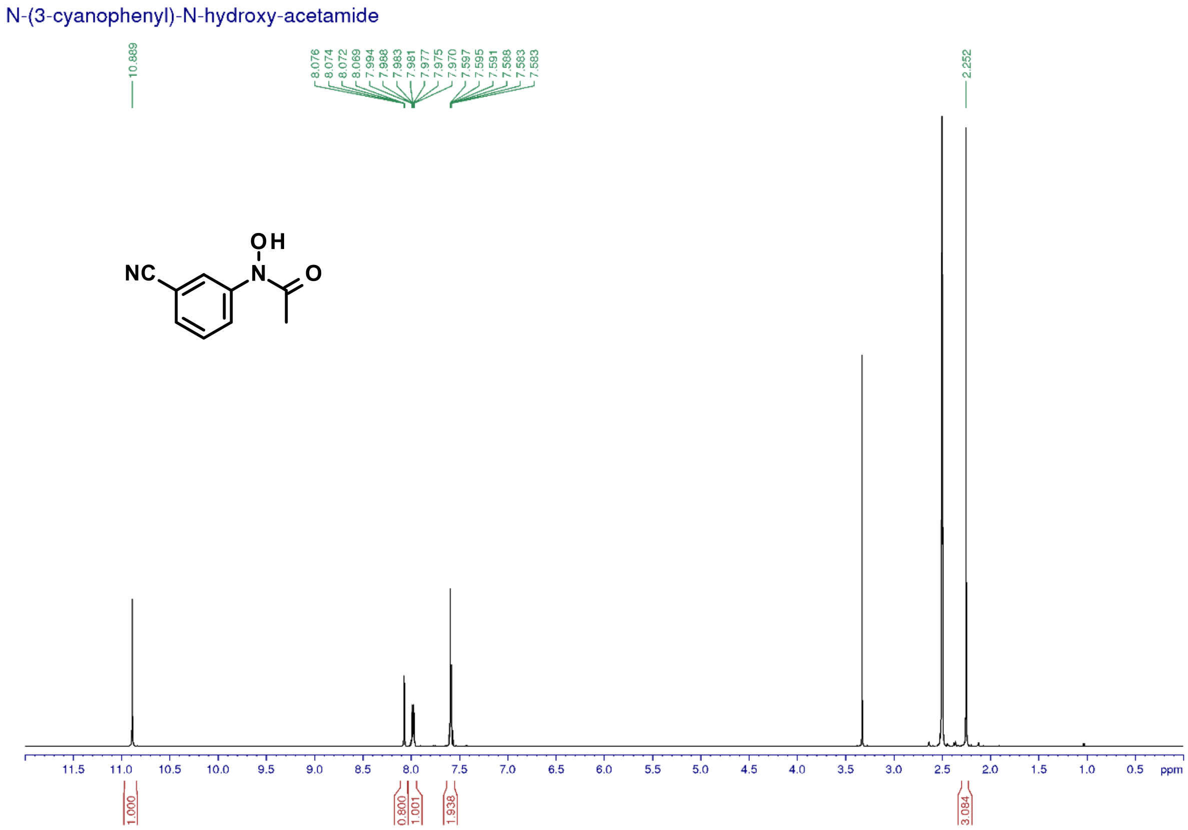

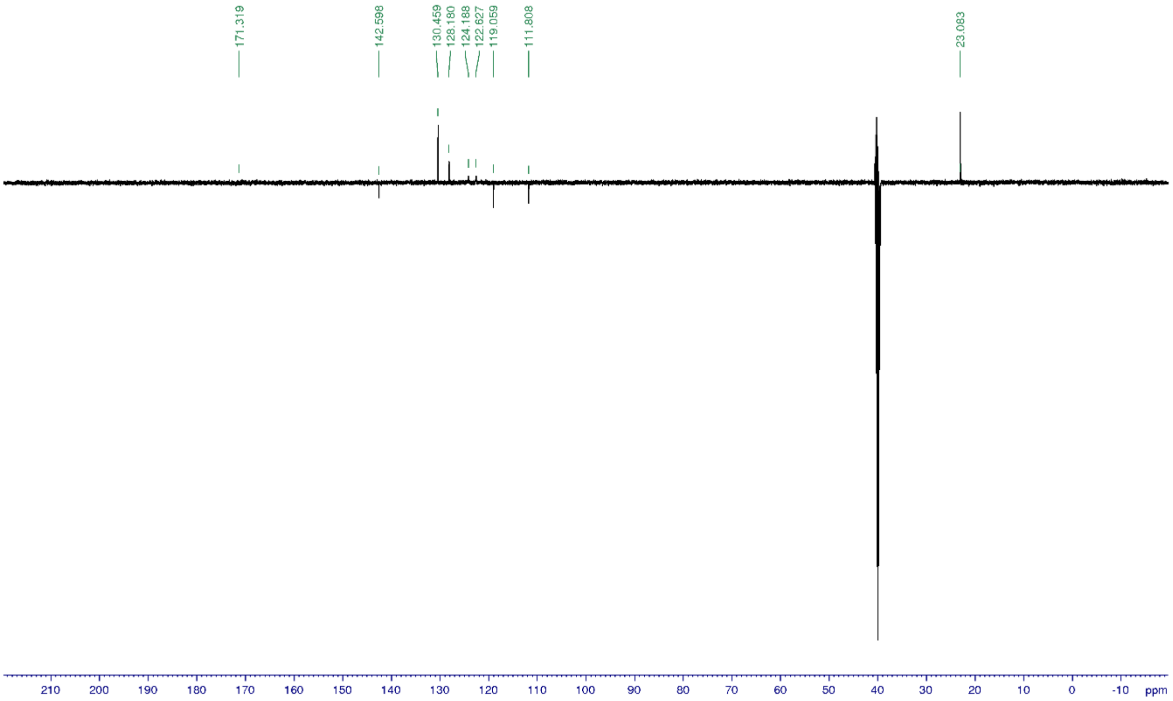


*2-Amino-6-cyano-1H-indole-3-carboxamide (**25**)*

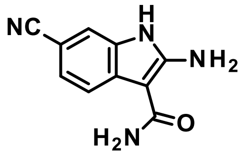


Title compound was synthesized according to the *General procedure C1* using *N*-(3-cyanophenyl)-*N*-hydroxy-acetamide (**24**, 1.6 g, 9.3 mmol) to give the desired product in mixture with 2-amino-4-cyano-1*H*-indole-3-carboxamide (1.0 g, 28%). The two isomers were not separated.

1H NMR (500 MHz, DMSO-d$_6$): δ ppm 10.96 (s, 1 H), 7.66 (d, J = 8.2 Hz, 1 H), 7.46 (d, J = 1.6 Hz,1 H), 7.28 (dd, J = 8.2, 1.6 Hz, 1 H), 7.18 (s, 2 H), 6.7 (s, 2 H); 13C NMR (125 MHz, DMSO-d$_6$): δ 168.4, 155.4, 132.1, 130.1, 124.3, 121.5, 117.1, 113.2, 99.1, 87.9; UV/Vis: $\lambda_{max}$ 236 nm; HRMS (m/z): [M+H]+ calcd. for $C_{10}H_8N_4O$, 201.0776; found, 201.0771.

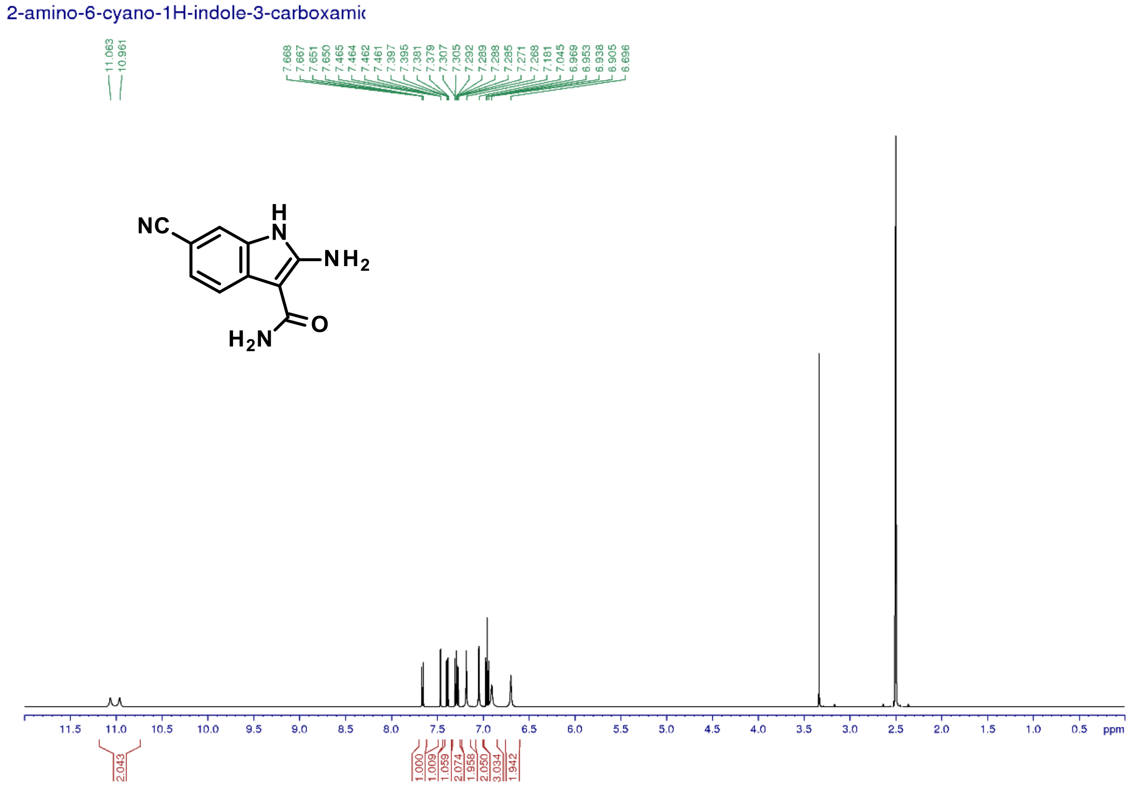


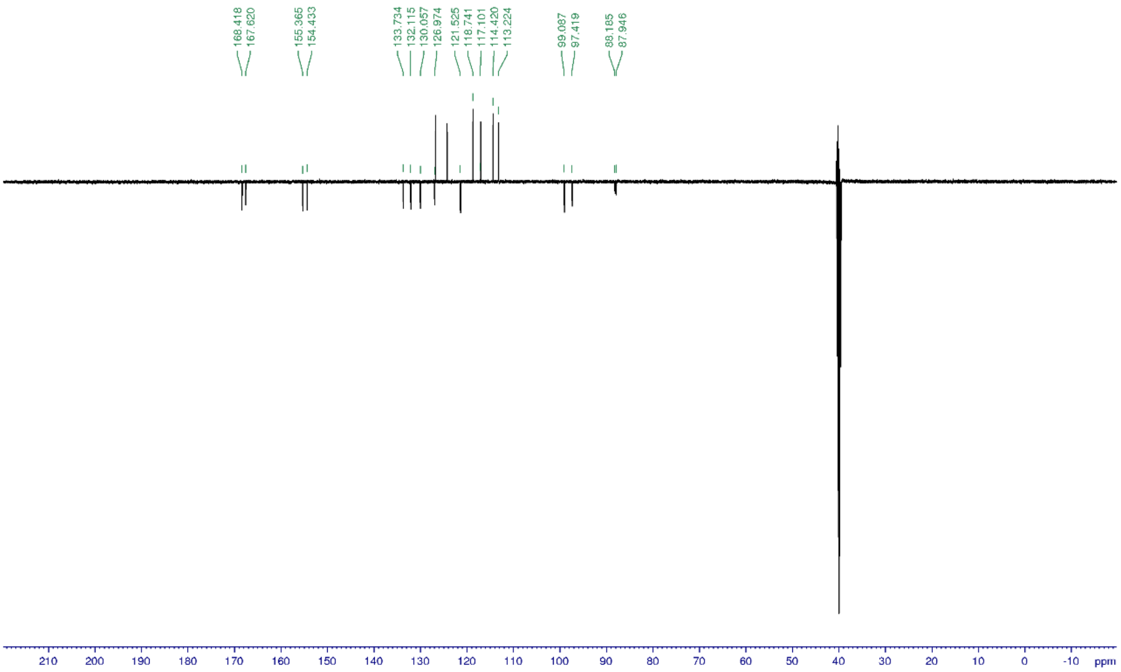


*2-Benzyl-4-hydroxy-9H-pyrimido[4,5-b]indole-7-carbonitrile* (***26***)

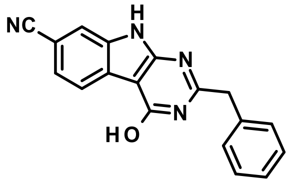


Title compound was synthesized according to the *General procedure D3* using the mixture of 2-amino-6-cyano-1*H*-indole-3-carboxamide (**25**) and 2-amino-4-cyano-1*H*-indole-3-carboxamide (481 mg, 2.4 mmol) to give the desired product (231 mg, 91%). The other isomer did not react.

1H NMR (500 MHz, DMSO-$d_6$): δ ppm 12.61 (brs, 2 H), 8.06 (dd, J = 8.1, 0.5 Hz, 1 H), 7.85 (dd, J = 1.3, 0.7 Hz, 1 H), 7.59 (dd, J = 8.3, 1.4 Hz, 1 H), 7.39 (d, J = 7.3 Hz, 2 H), 7.34 (t, J = 7.5 Hz, 2 H), 7.26 (tt, J = 7.3, 1.4 Hz, 1 H), 4.04 (s, 2 H); 13C NMR (125 MHz, DMSO-$d_6$): δ 161.3, 159.3, 156.6, 136.8, 135.1, 129.5, 129.0, 127.4, 126.2, 124.8, 121.4, 120.4, 116.2, 105.6, 98.6, 41.0; UV/Vis: $\lambda_{max}$ 248 nm; HRMS (m/z): [M+H]+ calcd. for $C_{18}H_{12}N_4O$, 301.1089; found, 301.1084.

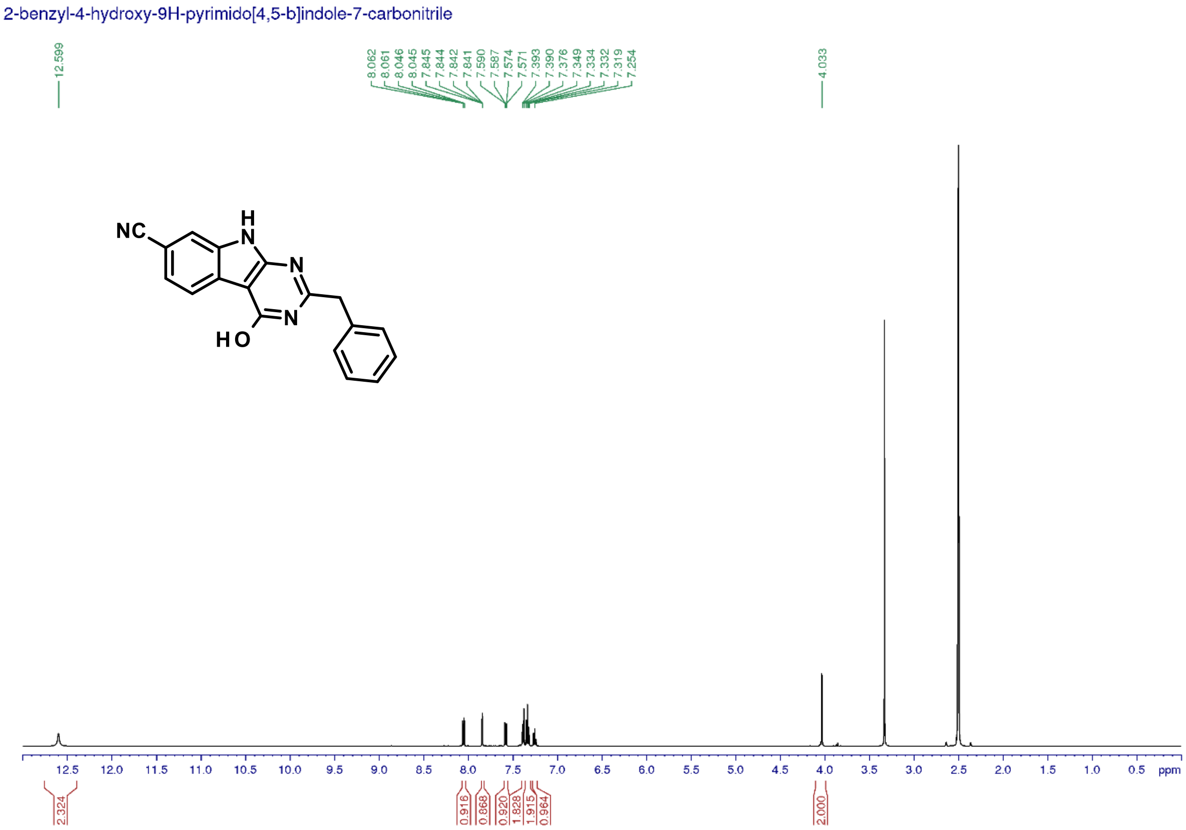

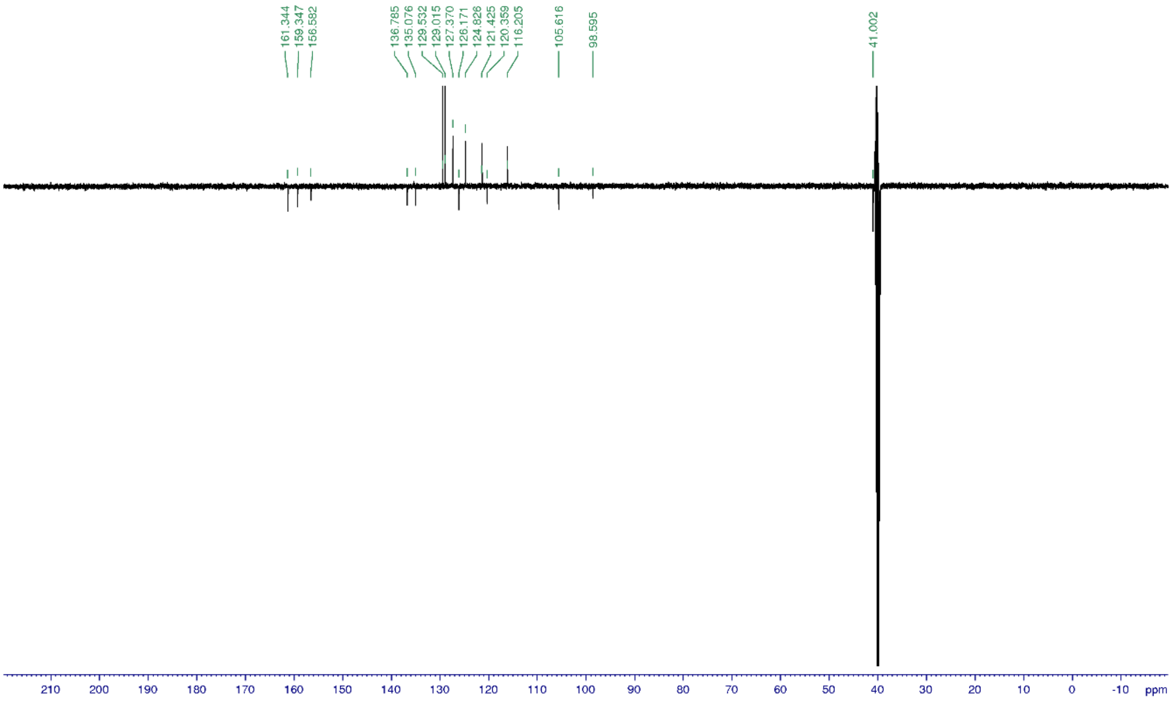


*2-Benzyl-4-hydroxy-9H-pyrimido[4,5-b]indole-7-carboxylic acid (**27**)*

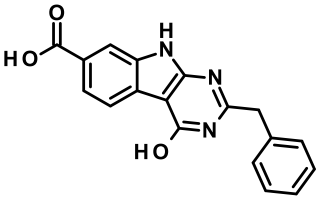


Title compound was synthesized according to the *General procedure E* using 2-benzyl-4-hydroxy-9H-pyrimido[4,5-b]indole-7-carbonitrile (**26**, 1.4 g, 4.8 mmol) to give the desired product (1.5 g, 99%).

1H NMR (400 MHz, DMSO-$d_6$): δ ppm 12.16 (br., 3 H), 7.99 (brs., 1 H), 7.96 (d, J = 8.4 Hz,1 H), 7.81 (dd, J = 8.2, 1.0 Hz, 1 H), 7.42-7.21 (m, 5 H), 4.02 (s, 2 H); 13C NMR (100 MHz, DMSO-$d_6$): δ 168.4, 159.5/156, 136.9, 135.4, 130.1, 122.7, 113.3, 40.9; UV/Vis: $\lambda_{max}$ 250 nm; HRMS (m/z): [M+H]+ calcd. for $C_{18}H_{13}N_3O_3$, 320.1035; found, 320.1031.

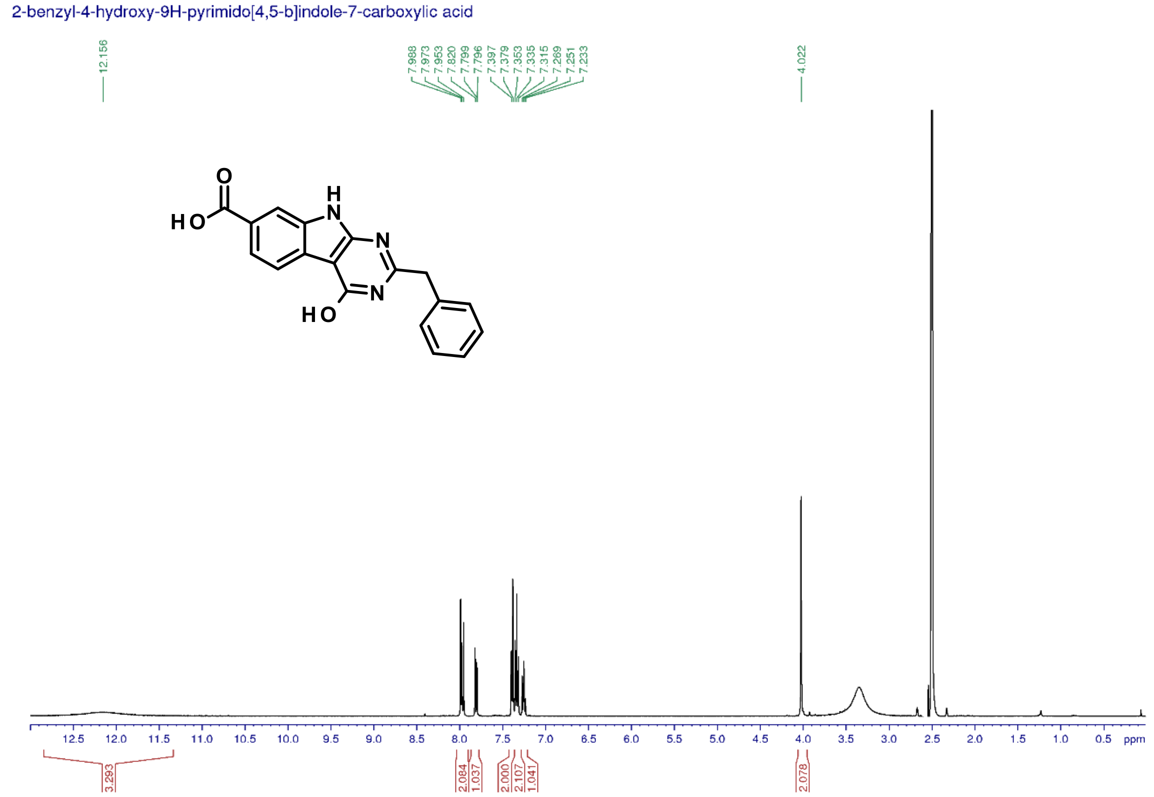


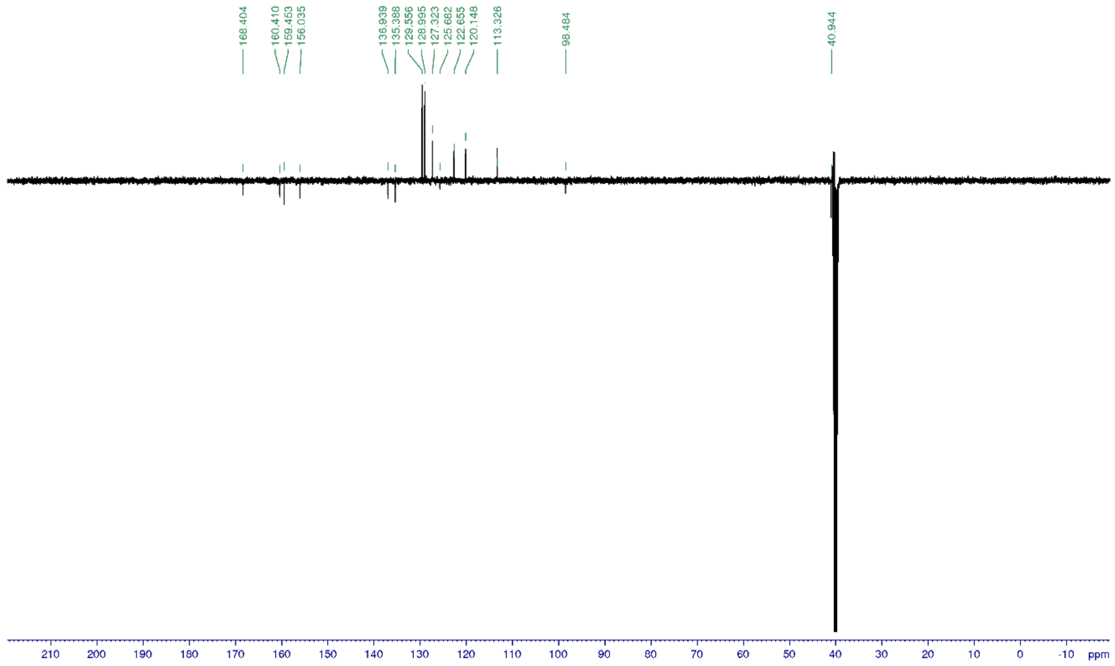

*Methyl 2-benzyl-4-hydroxy-9H-pyrimido[4,5-b]indole-7-carboxylate (**28**)*

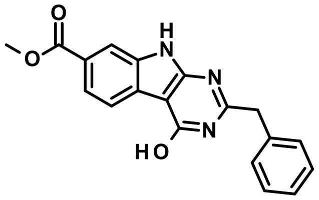

Title compound was synthesized according to the *General procedure F* using 2-benzyl-4-hydroxy-9H-pyrimido[4,5-b]indole-7-carboxylic acid (**27**, 120 mg, 0.4 mmol) to give the desired product (141 mg, 90%).

1H NMR (400 MHz, DMSO-$d_6$): δ ppm 12.54 (brs., 1 H), 12.48 (brs., 1 H), 8.03 (brs., 1 H), 8.01 (dd, J = 2.1, 1.0 Hz,1 H), 7.84 (dd, J = 8.2,1.4 Hz, 1 H), 7.42-7.22 (m, 5 H), 4.03 (s, 2 H), 3.88 (s, 3 H); 13C NMR (100 MHz, DMSO-$d_6$): δ 167.1, 160.8, 159.7, 156.3, 136.9, 135.4, 129.6, 127.3, 126.4, 122.4, 120.5, 113.3, 52.5, 40.9; UV/Vis: $\lambda_{max}$ 202 nm; HRMS (m/z): [M+H]+ calcd. for $C_{19}H_{15}N_3O_3$, 334.1191; found, 334.1189.

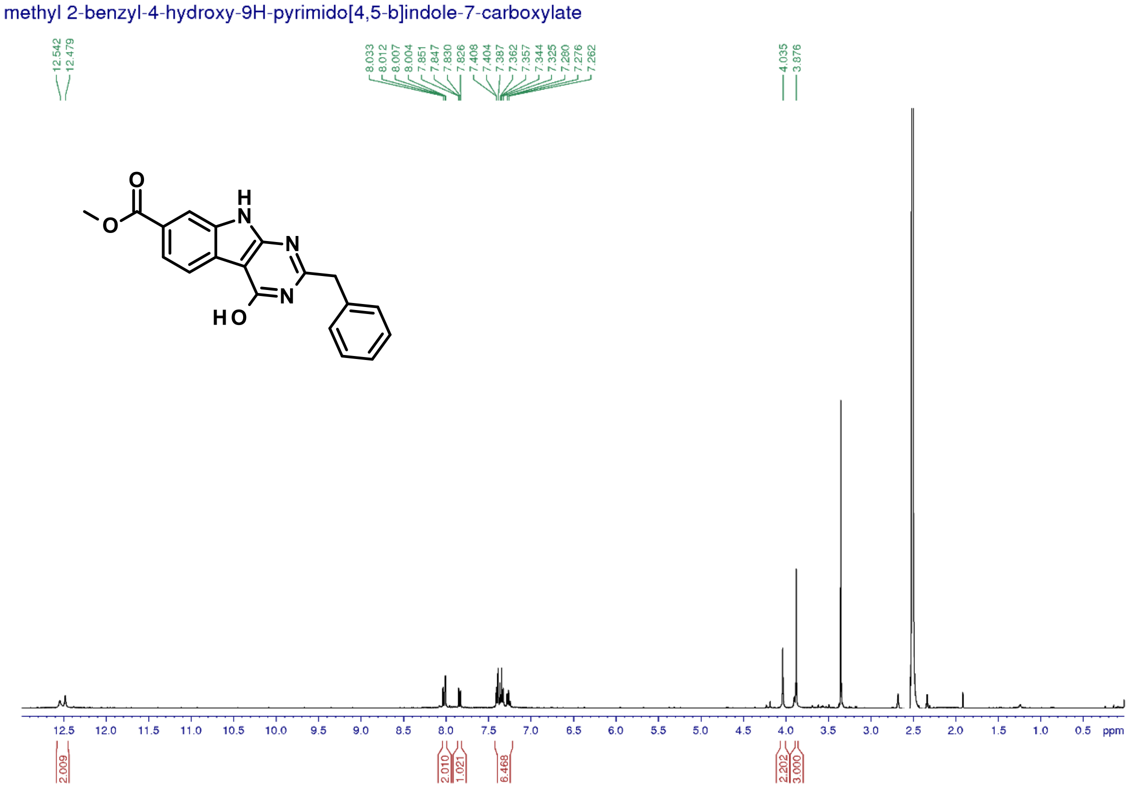

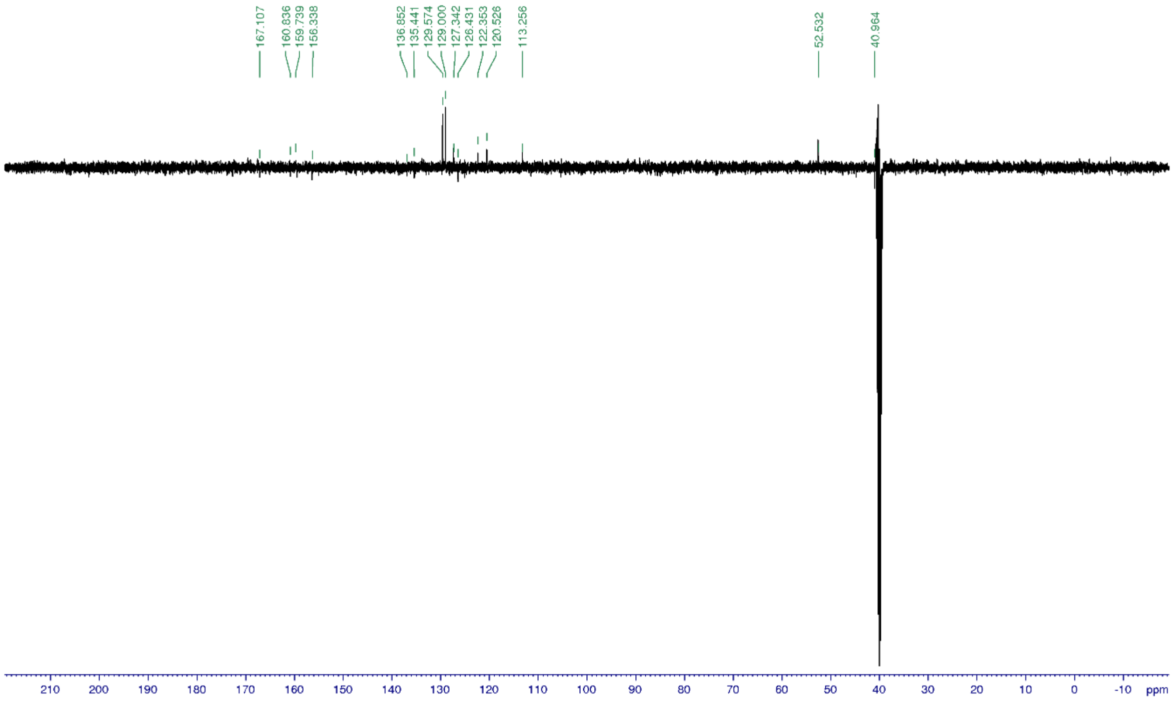


*Methyl 2-benzyl-4-chloro-9H-pyrimido[4,5-b]indole-7-carboxylate (**29**)*

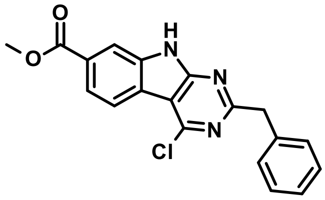


Title compound was synthesized according to the *General procedure G* using methyl 2-benzyl-4-hydroxy-9H-pyrimido[4,5-b]indole-7- carboxylate (**28**, 400 mg, 1.2 mmol) to give the desired product (402 mg, 95%).

1H NMR (400 MHz, DMSO-$d_6$): δ ppm 13.09 (brs., 1 H), 8.33 (d, J = 8.7 Hz ,1 H), 8.15 (brs.,1 H), 7.99 (dd, J = 8.3,1.5 Hz, 1 H), 7.40-7.28 (m, 5 H), 4. 31 (s, 2 H), 3.92 (s, 3 H); 13C NMR (100 MHz, DMSO-$d_6$): δ 167.1, 166.6, 158.1, 152.9, 138.5, 129.6, 128.9, 126.9, 122.7, 122.6, 122.1, 122.0, 113.6, 108.9, 52.9, 45.4; UV/Vis: $\lambda_{max}$ 241 nm; HRMS (m/z): [M+H]+ calcd. for $C_{19}H_{14}ClN_3O_2$, 352.0852; found, 352.0851.

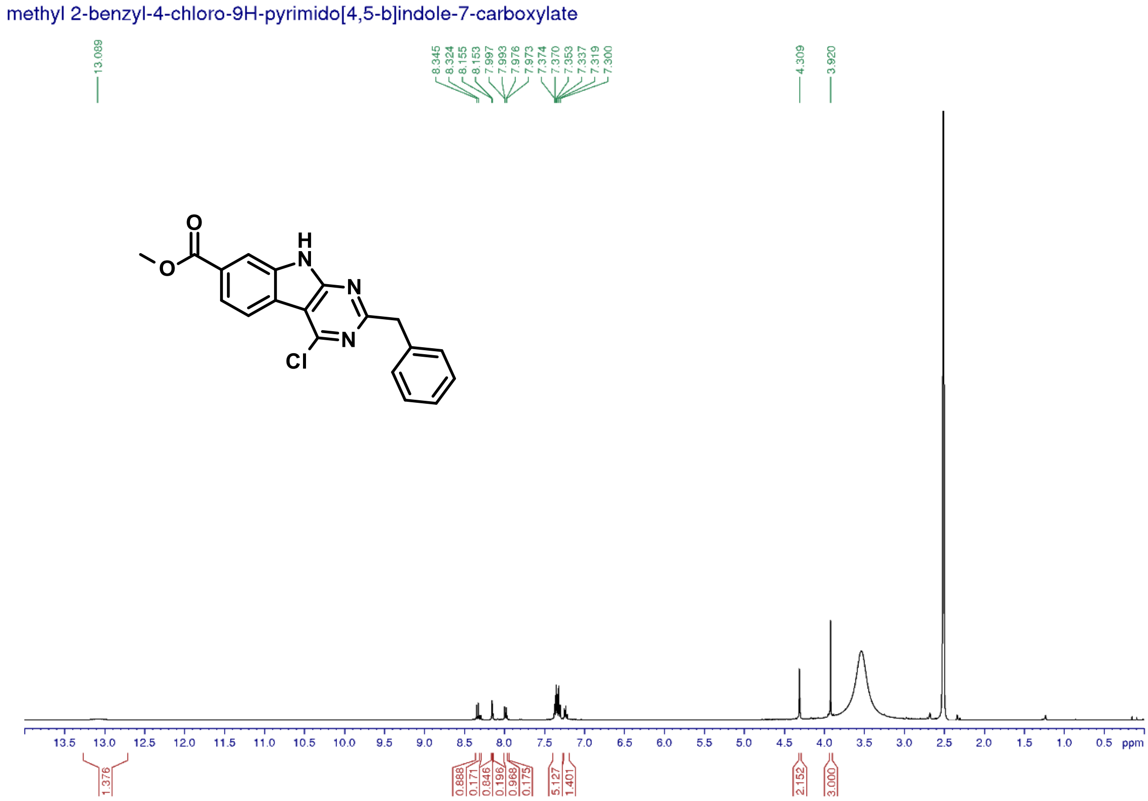


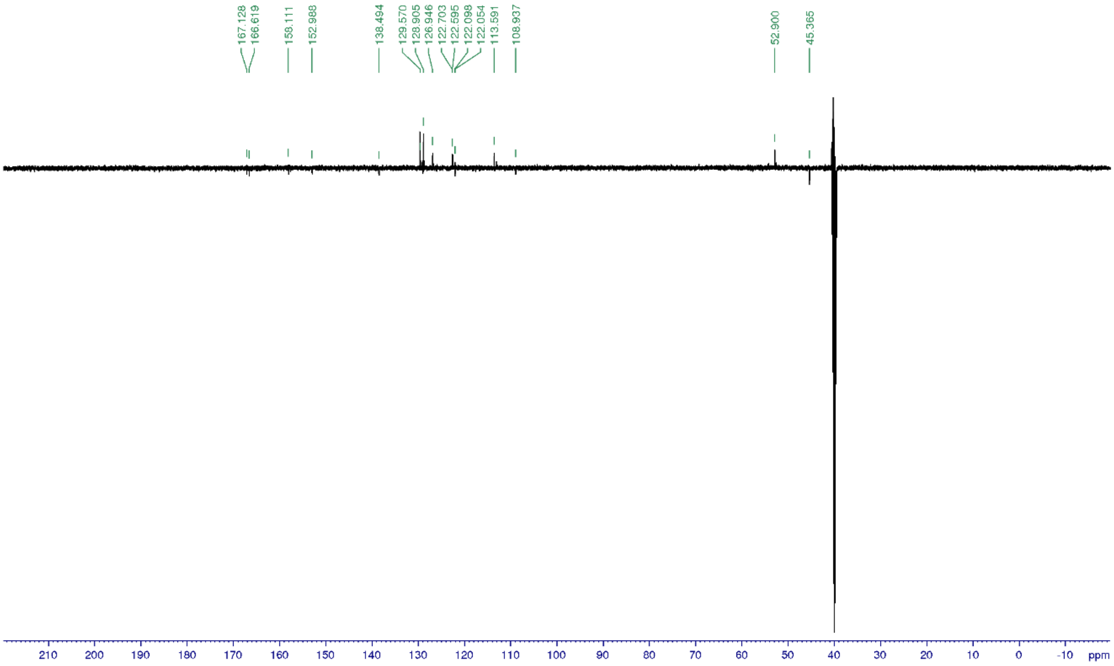

*Methyl 4-(hydroxyamino)benzoate (**30**)*

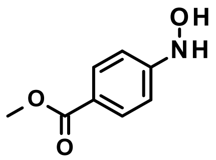

Title compound was synthesized according to the *General procedure A* using methyl 4-nitrobenzoate (9.0 g, 50 mmol) to give the desired product (4.8 g, 58%).

1H NMR (500 MHz, DMSO-$d_6$): δ ppm 8.96 (brd, 1 H), 8.65 (d, J = 1.7 Hz, 1 H), 7.77 (d, J = 8.7 Hz, 2 H), 6.83 (d, J = 8.7 Hz, 2 H), 3.77 (s, 3 H); 13C NMR (125 MHz, DMSO-$d_6$): δ 166.7, 156.4, 130.9, 119.5, 111.7, 51.9; UV/Vis: $\lambda_{max}$ 195 nm; HRMS (m/z): [M+H]+ calcd. for $C_8H_9NO_3$, 168.0660; found, 168.0654.

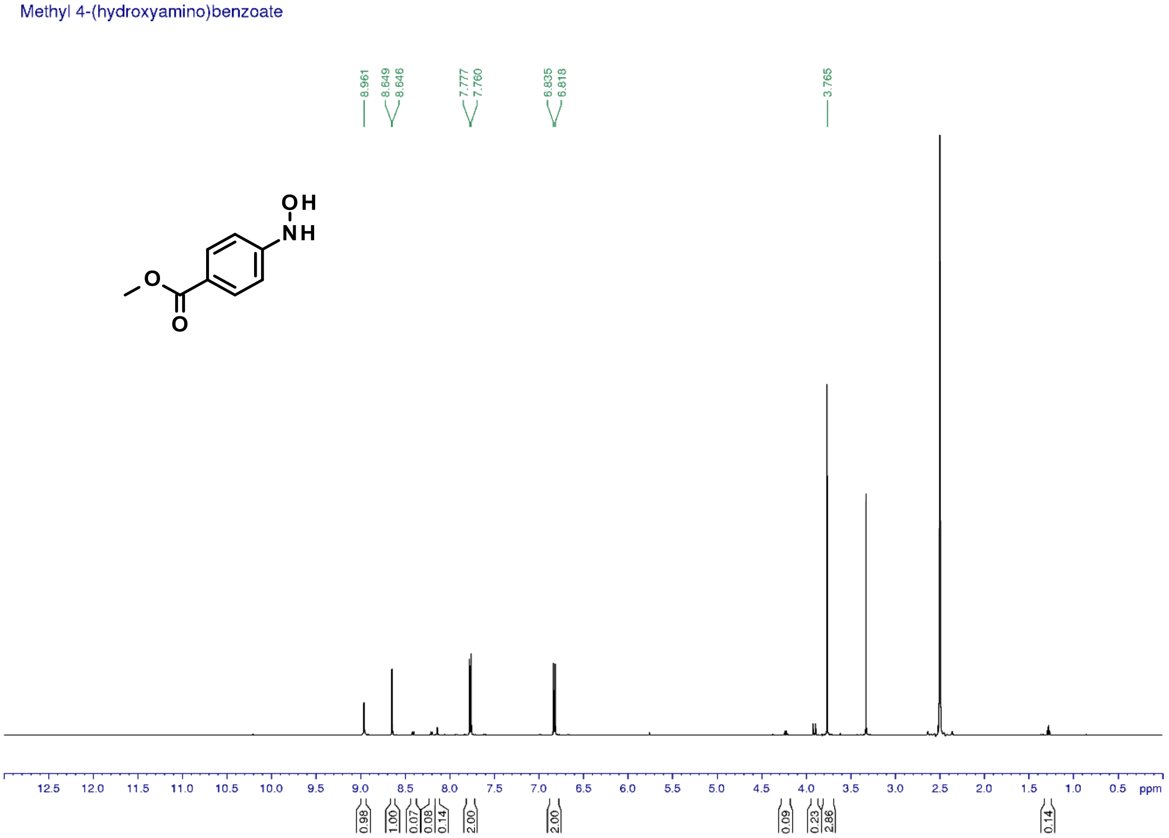

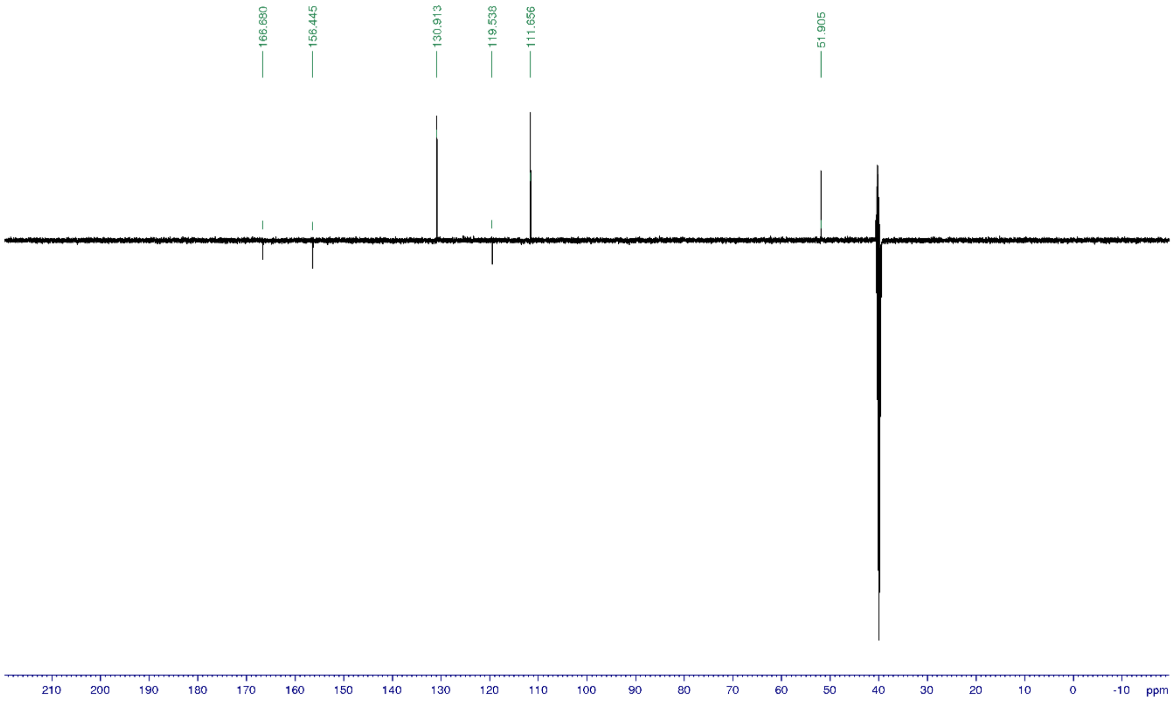


*Methyl 4-[acetyl(hydroxy)amino]benzoate (**31**)*

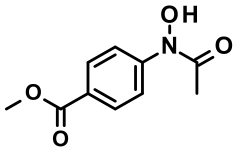

Title compound was synthesized according to the *General procedure B2* using methyl 4-(hydroxyamino)benzoate (**30**, 4.1 g, 25 mmol) and acetyl chloride (2.1 mL, 1.2 eq.) to give the desired product (5.0 g, 97%).

1H NMR (500 MHz, DMSO-$d_6$): δ ppm 10.86 (s, 1 H), 7.96 (dm, J = 9.1 Hz, 2 H), 7.84 (dm, J = 9.0 Hz, 2 H), 3.83 (s, 3 H), 2.26 (s, 3 H); 13C NMR (125 MHz, DMSO-$d_6$): δ 171.3, 166.2, 145.9, 130.3, 125.1, 118.9, 52.5, 23.4; UV/Vis: $\lambda_{max}$ 283 nm; HRMS (m/z): [M+H]+ calcd. for $C_{10}H_{11}NO_4$, 210.0766; found, 210.0761.

Methyl 4-[acetyl(hydroxy)amino]benzoate

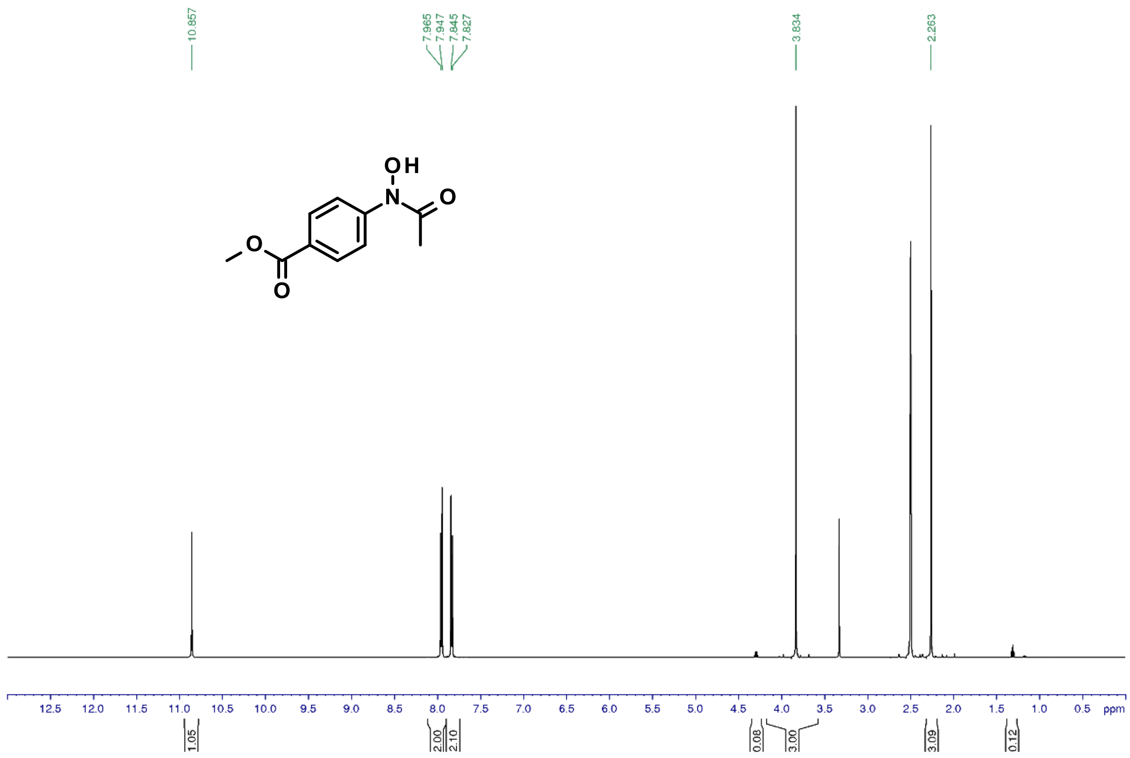


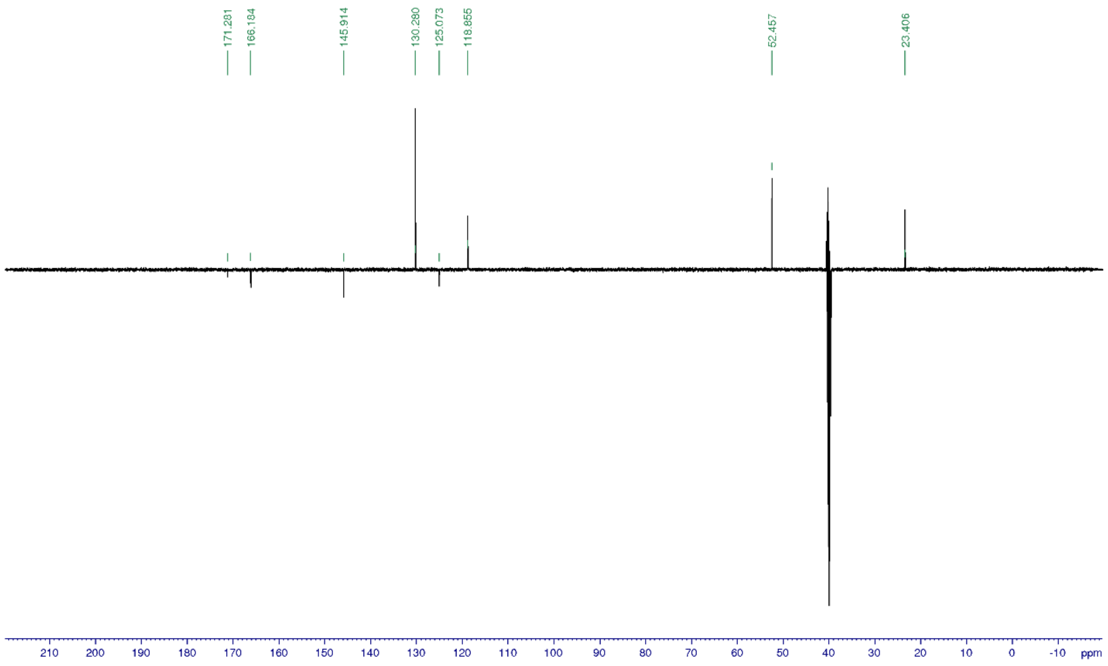


*Methyl 2-amino-3-carbamoyl-1H-indole-5-carboxylate (**32**)*

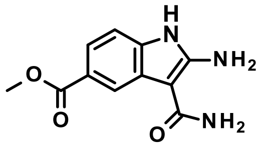


Title compound was synthesized according to the *General procedure C1* using methyl 4-[acetyl(hydroxy)amino]benzoate (**31**, 5.0 g, 24 mmol) to give the desired product (2.6 g, 45%).

1H NMR (500 MHz, DMSO-$d_6$): δ ppm 10.94 (s, 1 H), 8.11 (s, 1 H), 7.54 (dd, J = 8.3, 1.3 Hz,1 H), 7.18 (d, J = 8.2 Hz, 1 H), 6.89/6.56 (brs+brs., 4 H), 3.82 (s, 3 H); 13C NMR (125 MHz, DMSO-$d_6$): δ 168.9, 167.9, 153.9, 136.1, 126.1, 121.7, 120.8, 118.6, 109.7, 87.1, 52.0; UV/Vis: $\lambda_{max}$ 235 nm; HRMS (m/z): [M+H]+ calcd. for $C_{11}H_{11}N_3O_3$, 234.0878; found, 234.0873.

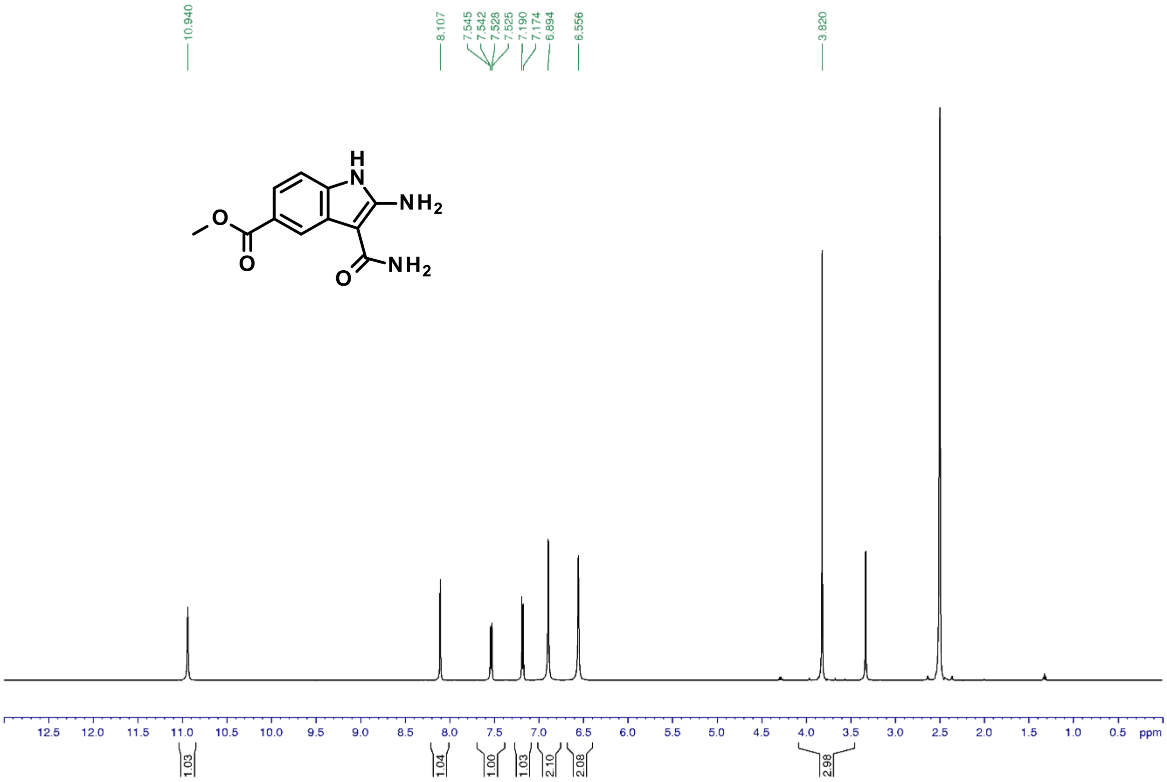

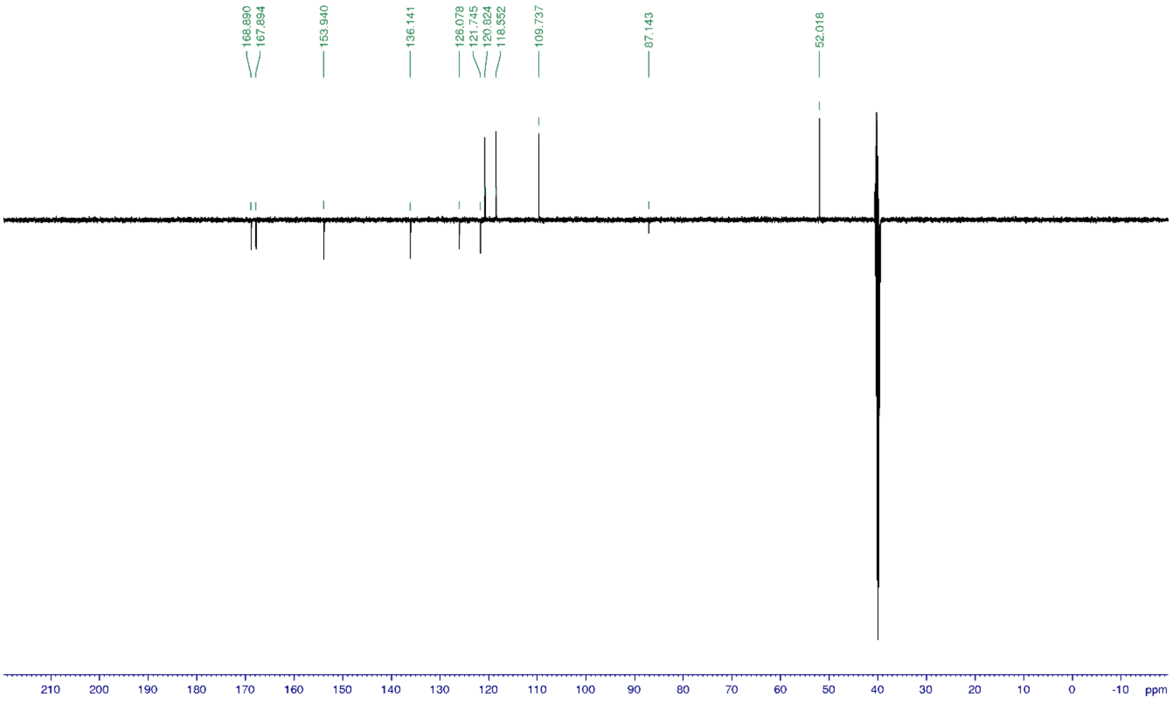


*Methyl 2-benzyl-4-hydroxy-9H-pyrimido[4,5-b]indole-6-carboxylate (**33**)*

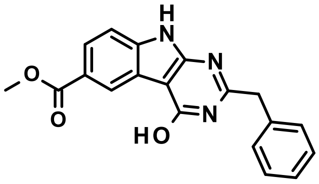

Title compound was synthesized according to the *General procedure D3* using methyl 2-amino-3-carbamoyl-1H-indole-5-carboxylate (**32**, 500 mg, 2.1 mmol) to give the desired product (409 mg, 51%).

1H NMR (500 MHz, DMSO-$d_6$): δ ppm 12.52 (brs, 2 H), 8.59 (dd, J = 1.7, 0.6 Hz, 1 H), 7.92 (dd, J = 8.5, 1.8 Hz, 1 H), 7.50 (dd, J = 8.5, 0.5 Hz, 1 H), 7.38 (d, J = 7.5 Hz, 2 H), 7.33 (t, J = 7.4 Hz, 2 H), 7.25 (t, J = 7.3 Hz, 1 H), 4.03 (s, 2 H), 3.87 (s, 3 H); 13C NMR (125 MHz, DMSO-$d_6$): δ 167.2, 160.4, 159.3/156.0, 138.9, 136.9, 129.5, 129.0, 127.3, 125.4, 122.8, 122.5, 122.2, 112.0, 98.9, 52.4, 41.0; UV/Vis: $\lambda_{max}$ 256 nm; HRMS (m/z): [M+H]+ calcd. for $C_{19}H_{15}N_3O_3$, 334.1191; found, 334.1186.

Methyl 2-benzyl-4-hydroxy-9H-pyrimido[4,5-b]indole-6-carboxylate

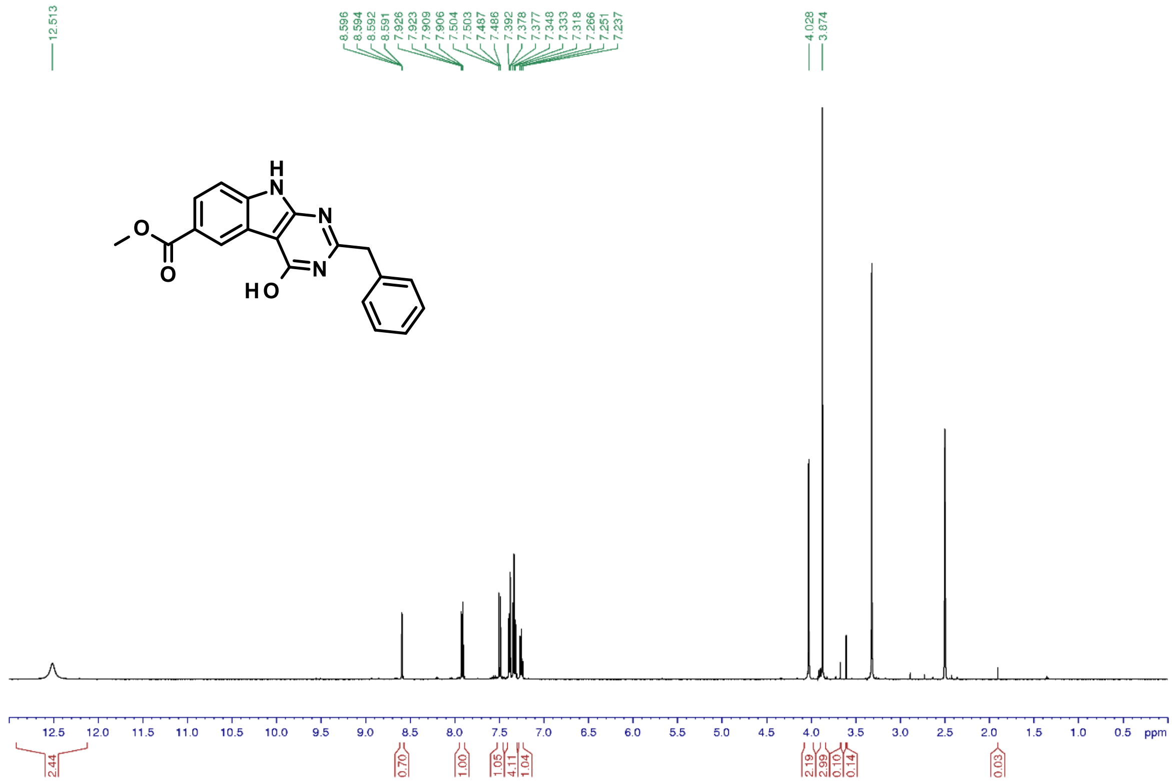


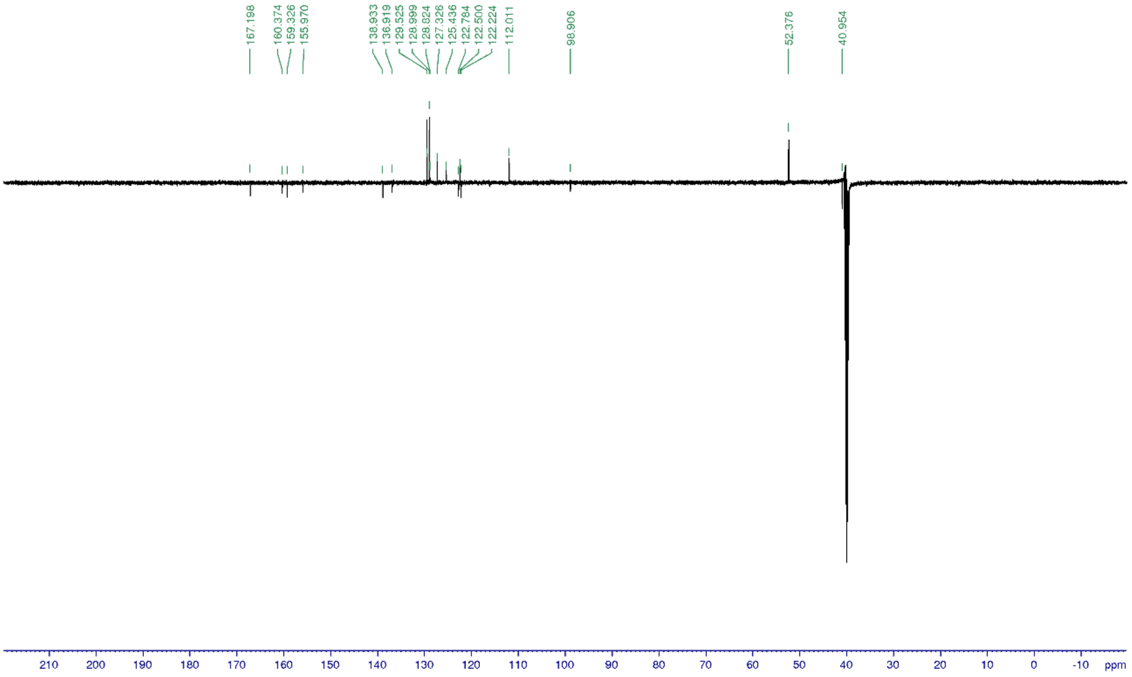


*Methyl 2-benzyl-4-chloro-9H-pyrimido[4,5-b]indole-6-carboxylate* (***34***)

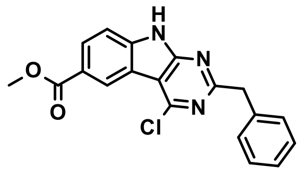

Title compound was synthesized according to the *General procedure G* using methyl 2-benzyl-4-hydroxy-9H-pyrimido[4,5-b]indole-6-carboxylate (**33**, 408 mg, 1.2 mmol) to give the desired product (389 mg, 90%).

1H NMR (400 MHz, DMSO-$d_6$): δ ppm 13.05 (brs, 1 H), 8.78 (d, J = 1.7 Hz, 1 H), 8.16 (dd, J = 8.6, 1.7 Hz, 1 H), 7.66 (d, J = 8.5 Hz, 1 H), 7.35 (d, J = 6.8 Hz, 2 H), 7.31 (t, J = 7.3 Hz, 2 H), 7.22 (t, J = 7.3 Hz, 1 H), 4.30 (s, 2 H), 3.90 (s, 3 H); 13C NMR (100 MHz, DMSO-$d_6$): δ 166.7, 166.7, 158.3/152.2, 142.1, 138.7, 129.6, 129.4, 128.9, 126.9, 124.0, 123.3/118.2, 112.9, 109.5, 52.7, 45.3; UV/Vis: $\lambda_{max}$ 266 nm; HRMS (m/z): [M+H]+ calcd. for $C_{19}H_{14}ClN_3O_2$, 352.0852; found, 352.0851.

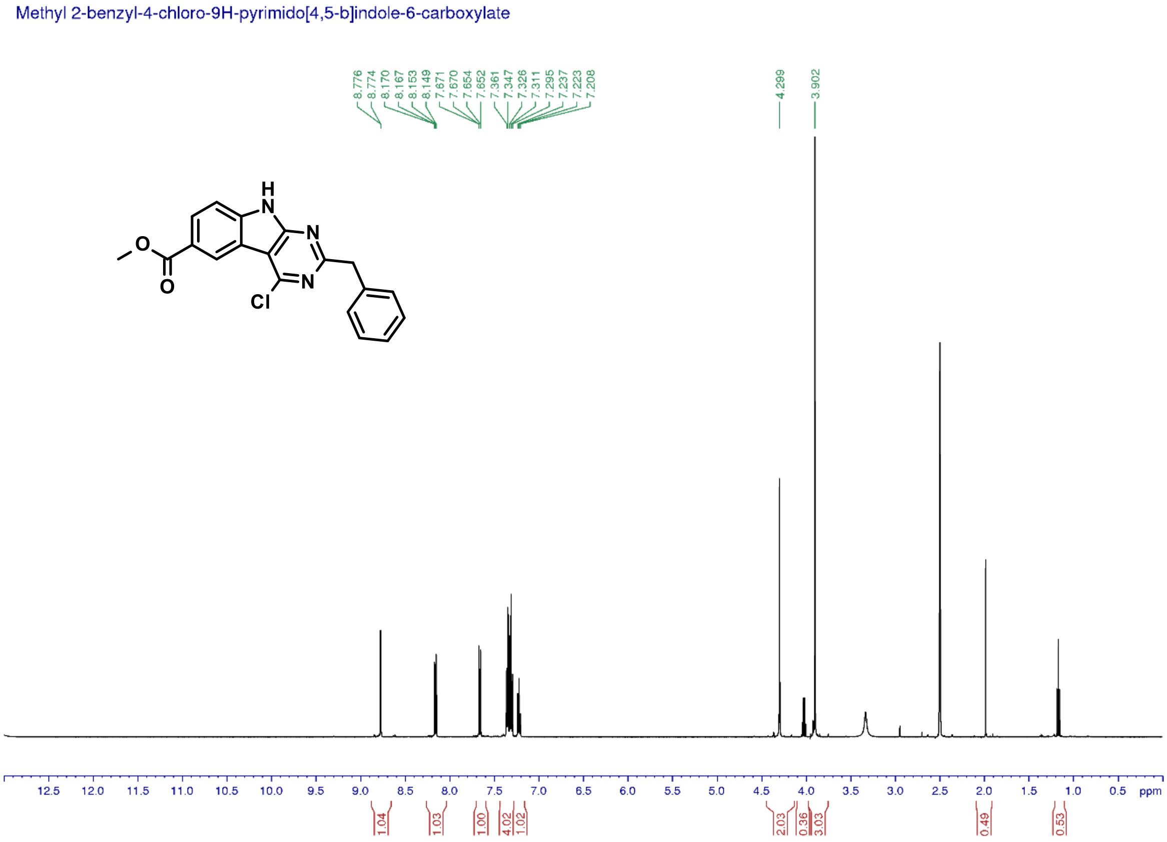

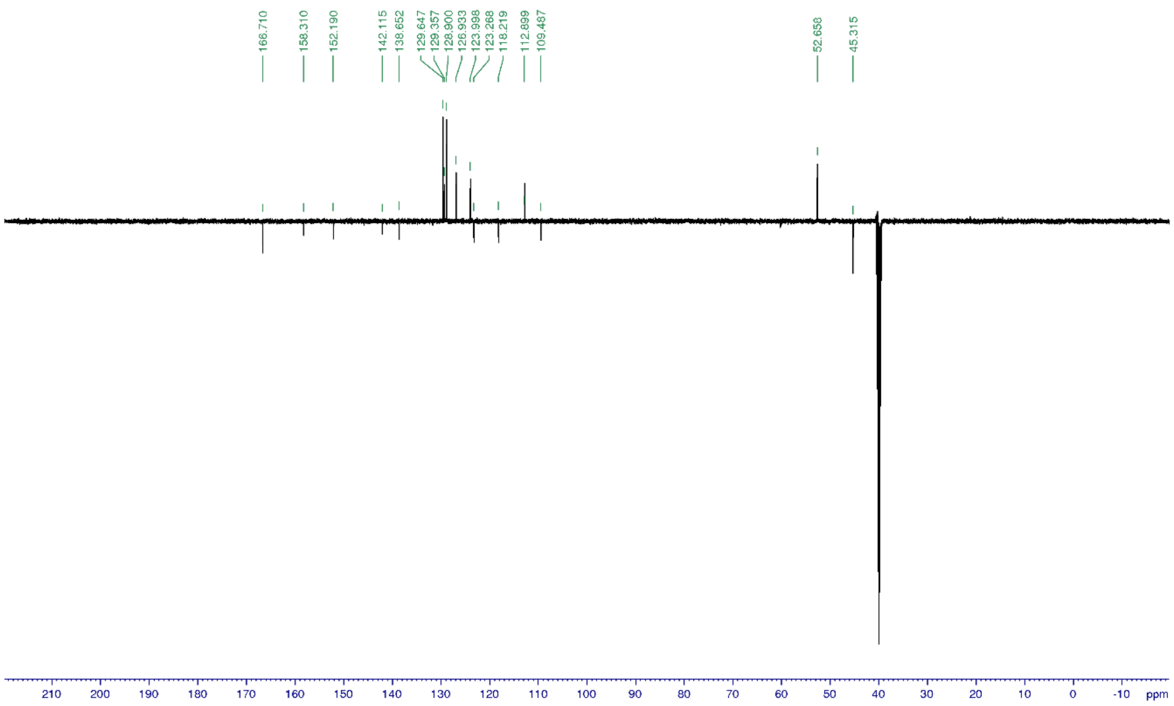


*Methyl 2-benzyl-4-[[4-(tert-butoxycarbonylamino)cyclohexyl]amino]-9H-pyrimido[4,5-b]indole-6-carboxylate (**35**)*

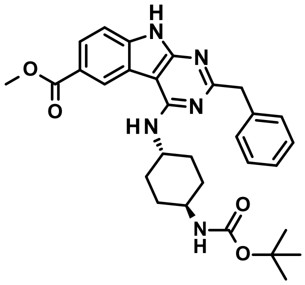

Title compound was synthesized according to the *General procedure H3* using methyl 2-benzyl-4-chloro-9H-pyrimido[4,5-b]indole-6-carboxylate (**34**, 194 mg, 0.55 mmol) and tert-butyl N-(4-aminocyclohexyl)carbamate (2.0 eq., 236 mg) to give the desired product (72 mg, 25%).

1H NMR (500 MHz, DMSO-$d_6$): δ ppm 12.15 (brs, 1 H), 8.88 (dd, J = 1.5, 0.5 Hz, 1 H), 7.94 (dd, J = 8.5, 1.7 Hz, 1 H), 7.45 (dd, J = 8.4, 0.4 Hz, 1 H), 7.38 (d, J = 7.6 Hz, 2 H), 7.29 (t, J = 7.4 Hz, 2 H), 7.19 (t, J = 7.2 Hz, 1 H), 7.09 (d, J = 7.8 Hz, 1 H), 6.80 (d, J = 7.8 Hz, 1 H), 4.21 (m, 1 H), 4.04 (s, 2 H), 3.89 (s, 3 H), 3.28 (m, 1 H), 1.93/1.62 (m+m, 4 H), 1.85/1.32 (m+m, 4 H), 1.37 (s, 9 H); 13C NMR (125 MHz, DMSO-$d_6$): δ 167.6, 166.3, 157.8, 165.5, 155.3, 139.8, 139.8, 129.7, 128.5, 126.5, 125.9, 123.6, 121.8, 120.0, 111.0, 94.2, 77.9, 52.3, 49.3, 49.2, 46.1, 32.3, 31.1, 28.8; UV/Vis: $\lambda_{max}$ 266 nm; HRMS (m/z): [M+H]+ calcd. for C30H35N5O4, 530.2767; found, 530.2764.

Methyl 2-benzyl-4-[[4-(tert-butoxycarbonylamino)cyclohexyl]amino]-9H-pyrimido[4,5-b]indole-6-carboxylate

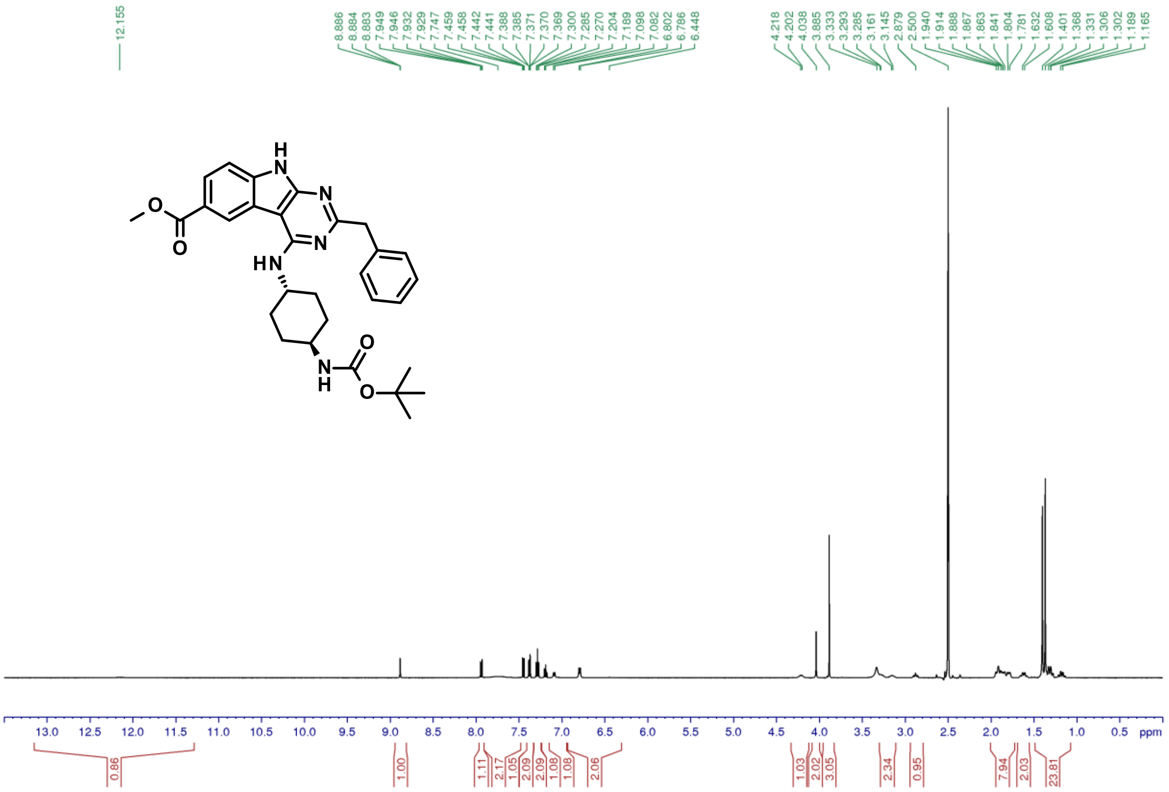


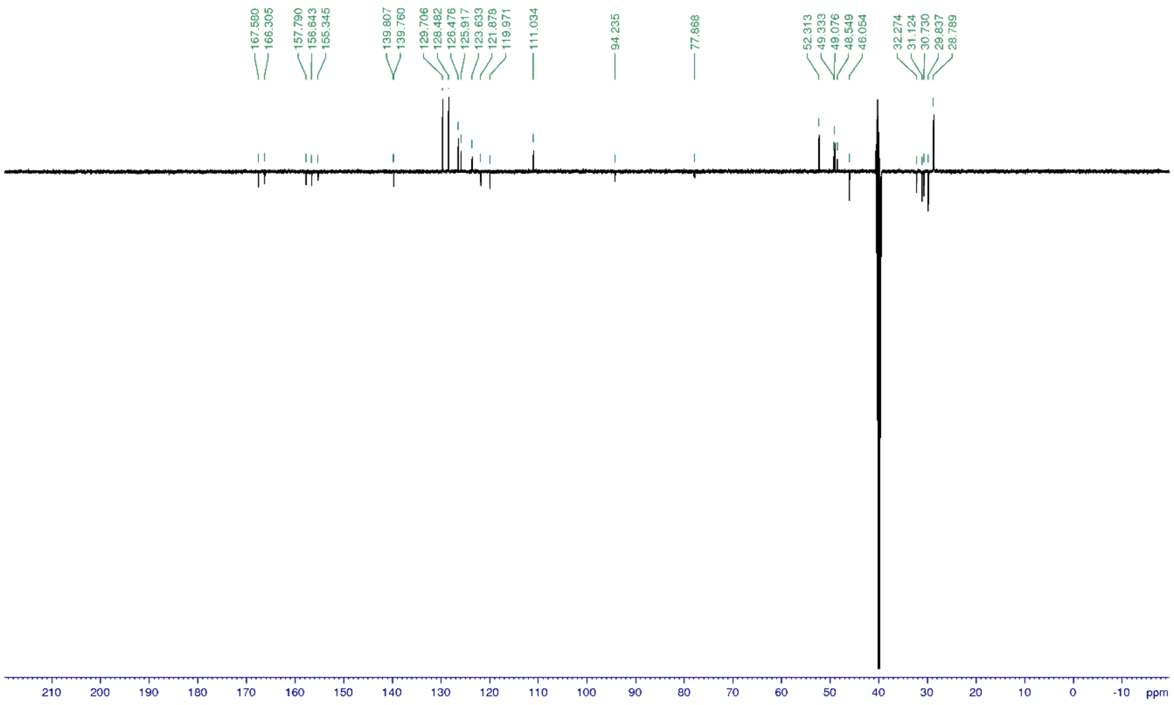

*N-[3-(2-methyltetrazol-5-yl)phenyl]hydroxylamine (**36**)*

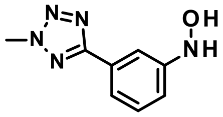

Title compound was synthesized according to the *General procedure A* using 2-methyl-5-(3-nitrophenyl)tetrazole (2.0 g, 9.7 mmol) to give the desired product (1.4 g, 74%).

1H NMR (500 MHz, DMSO-d$_6$): δ ppm 8.54 (d, J = 1.6 Hz, 1H), 8.49 (d, J = 2.1 Hz, 1H), 7.57 (t, J = 1.9 Hz, 1 H), 7.43 (dm, 1 H), 7.33 (t, J = 8.0 Hz, 1 H), 6.95 (dm, 1 H), 4.41 (s, 3 H); 13C NMR (125 MHz, DMSO-d$_6$): δ 165.0, 153.2, 129.8, 127.5, 117.5, 115.3, 110.9, 40.0; UV/Vis: $\lambda_{max}$ 239 nm; HRMS (m/z): [M+H]+ calcd. for $C_8H_9N_5O$, 192.0885; found, 192.0879.

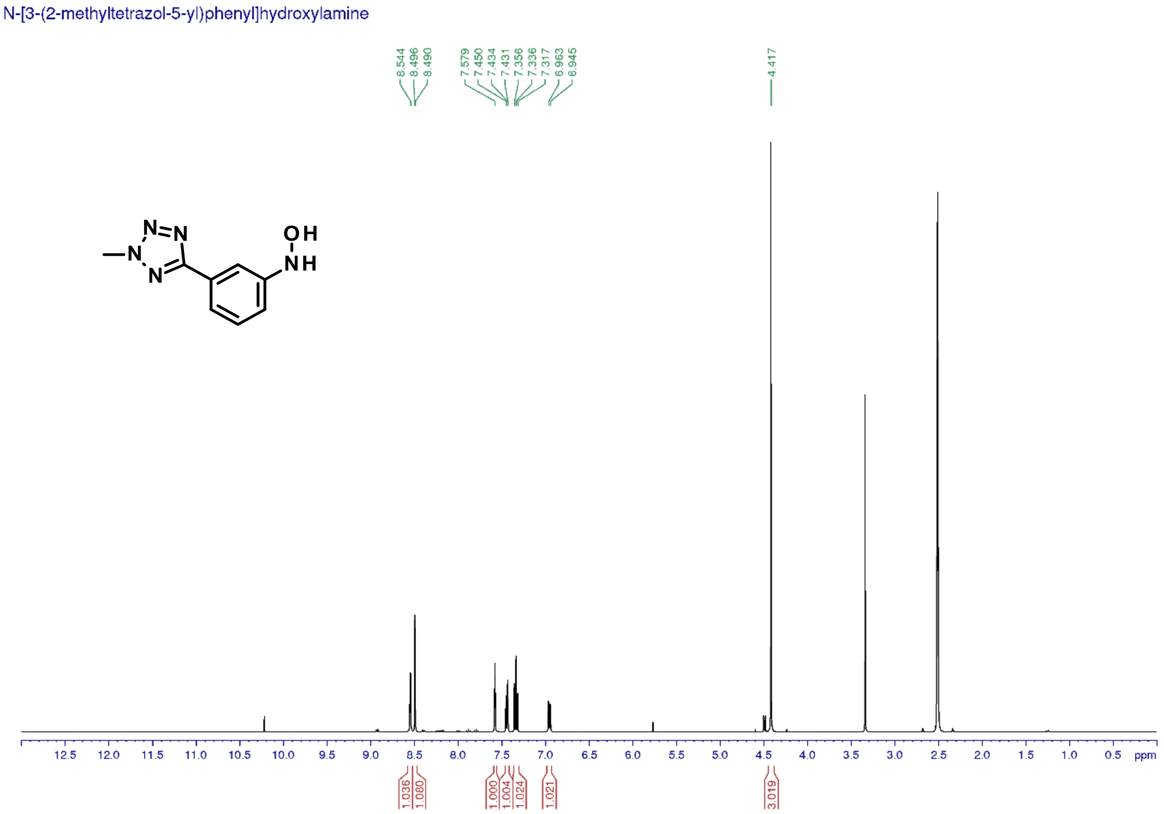

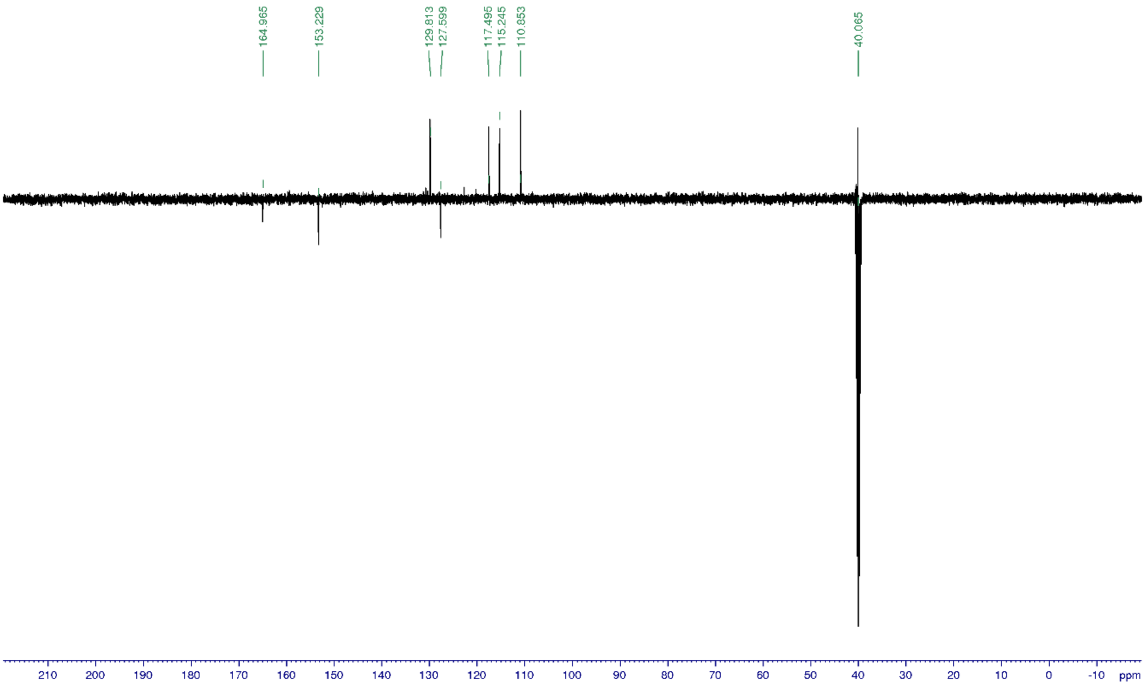


*N-hydroxy-N-[3-(2-methyltetrazol-5-yl)phenyl]acetamide (**37**)*

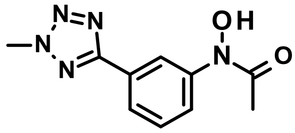

Title compound was synthesized according to the *General procedure B1* using N-[3-(2-methyltetrazol-5-yl)phenyl]hydroxylamine (**36**, 1.4 g, 7.3 mmol) and acetyl chloride (1.1 eq., 570 uL) to give the desired product (1.0 g, 62%).

1H NMR (500 MHz, DMSO-$d_6$): δ ppm 10.81 (s, 1 H), 8.40 (t, J = 1.8 Hz, 1 H), 7.84-7.79 (m, 2 H), 7.55 (t, J = 8.0 Hz, 1 H), 4.43 (s, 3 H), 2.26 (s, 3 H); 13C NMR (125 MHz, DMSO-$d_6$): δ 170.8, 164.4, 142.8, 129.9, 127.6, 117.6, 40.2, 23.1; UV/Vis: $\lambda_{max}$ 240 nm; HRMS (m/z): [M+H]+ calcd. for $C_{10}H_{11}N_5O_2$, 234.0991; found, 234.0986.

N-hydroxy-N-[3-(2-methyltetrazol-5-yl)phenyl]acetamide

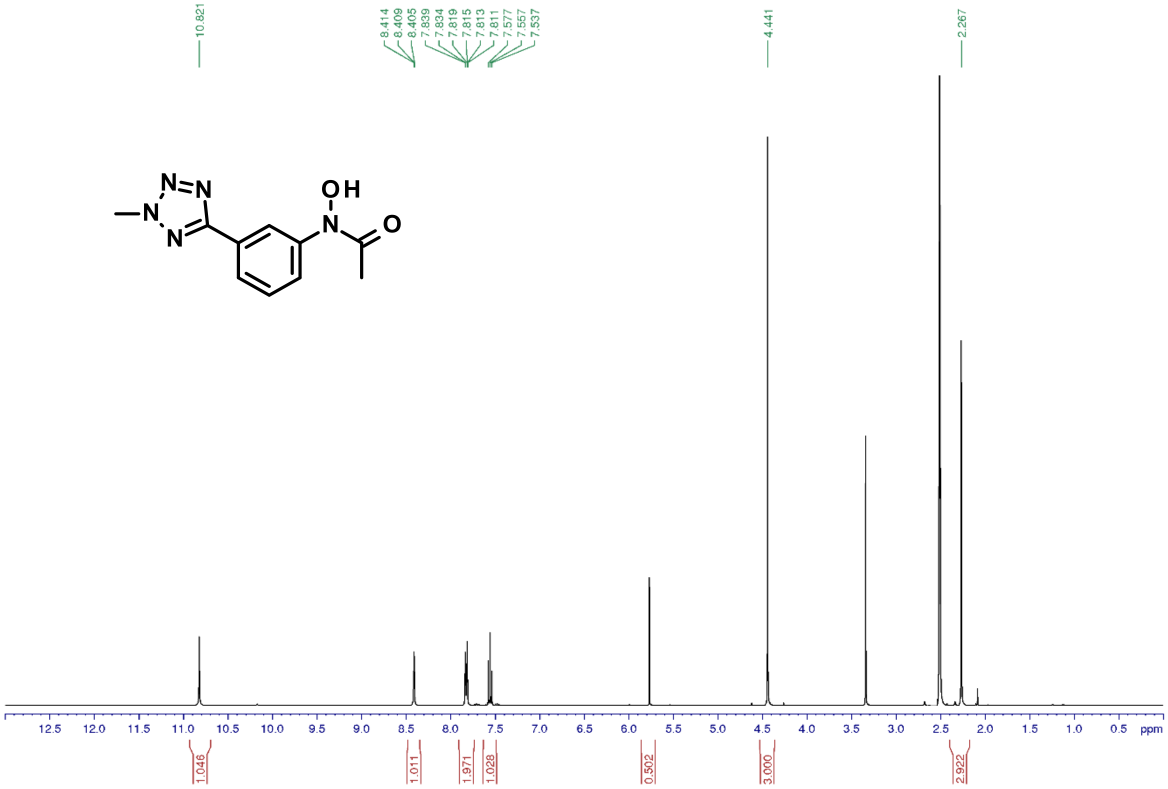


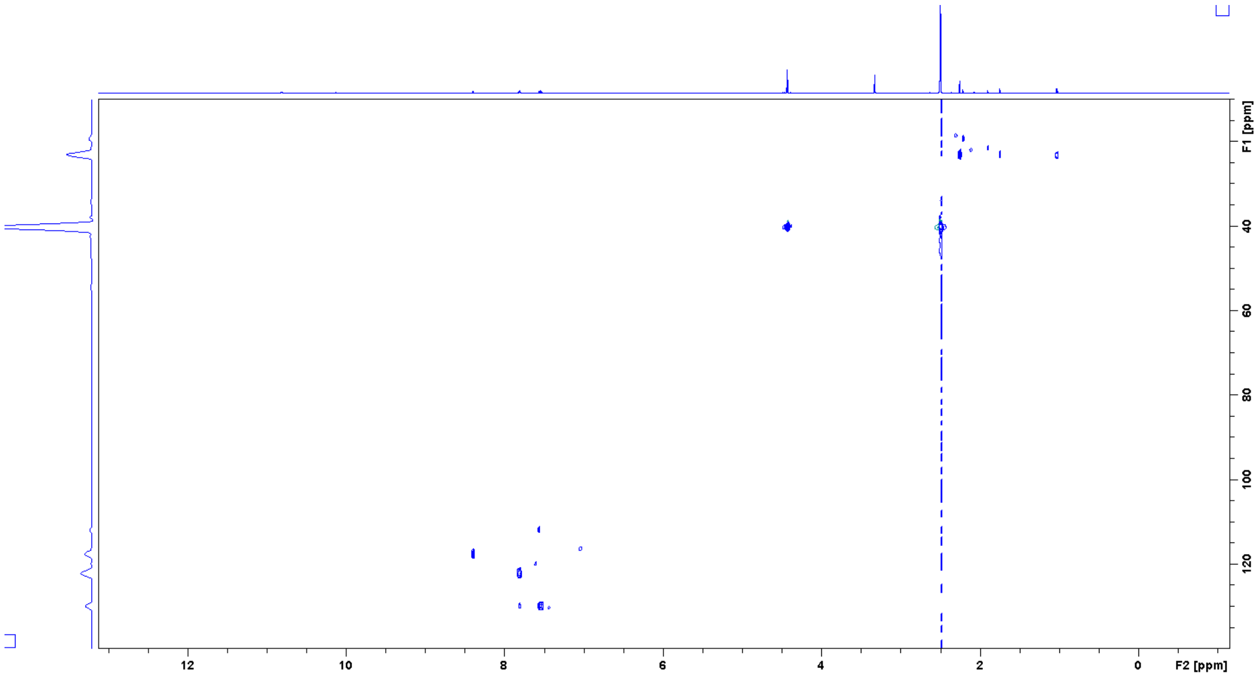

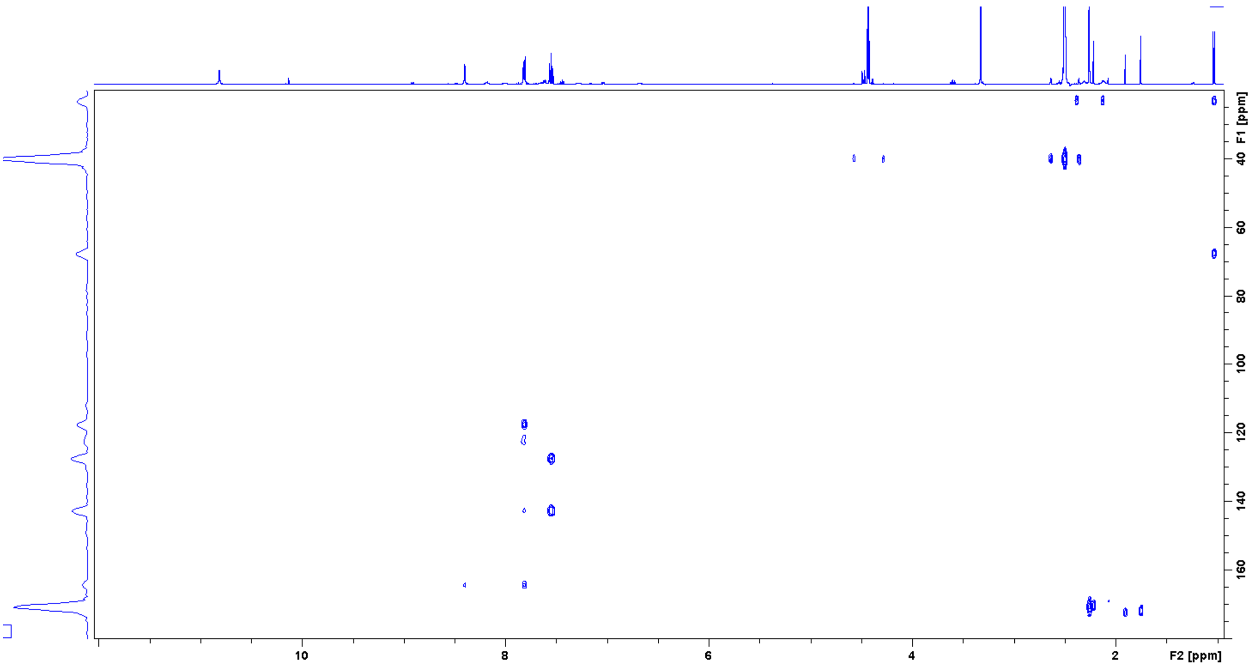


*2-Amino-6-(2-methyltetrazol-5-yl)-1H-indole-3-carboxamide (**38**)*

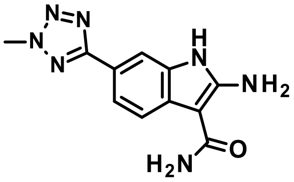

Title compound was synthesized according to the *General procedure C1* using N-hydroxy-N-[3-(2-methyltetrazol-5-yl)phenyl]acetamide (**37**, 1.0 g, 4.5 mmol) to give the desired product (322 mg, 28%).

1H NMR (500 MHz, DMSO-$d_6$): δ ppm 10.79 (s, 1 H), 7.81 (d, J = 1.3 Hz, 1 H), 7.67 (d, J = 8.2 Hz, 1 H), 7.64 (dd, J = 8.2, 1.3 Hz, 1 H), 7.6-6.0 (brs, 4 H), 4.38 (s, 3 H); 13C NMR (125 MHz, DMSO-$d_6$): δ 166.0, 154.6, 132.9, 128.0, 118.8, 117.2, 117.1, 108.0, 87.2, 39.9; UV/Vis: $\lambda_{max}$ 238 nm; HRMS (m/z): [M+H]+ calcd. for $C_{11}H_{11}N_7O$, 258.1103; found, 258.1099.

2-Amino-6-(2-methyltetrazol-5-yl)-1H-indole-3-carboxamide

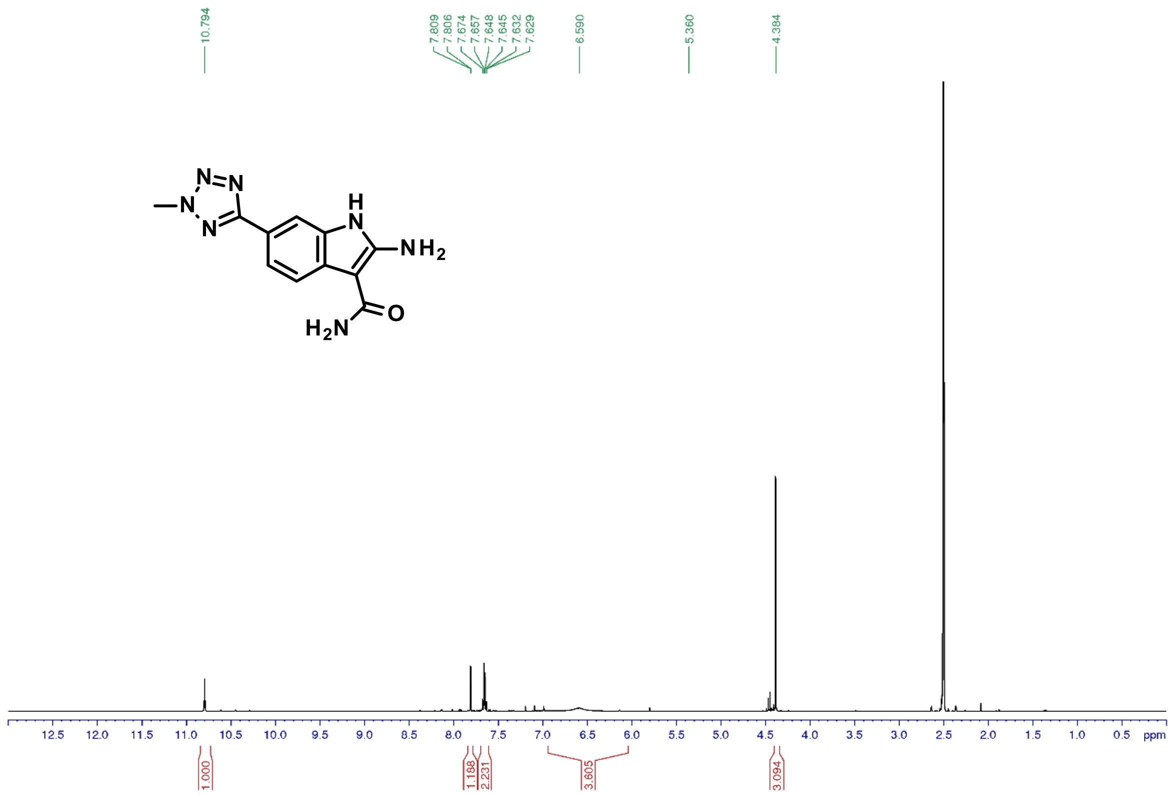


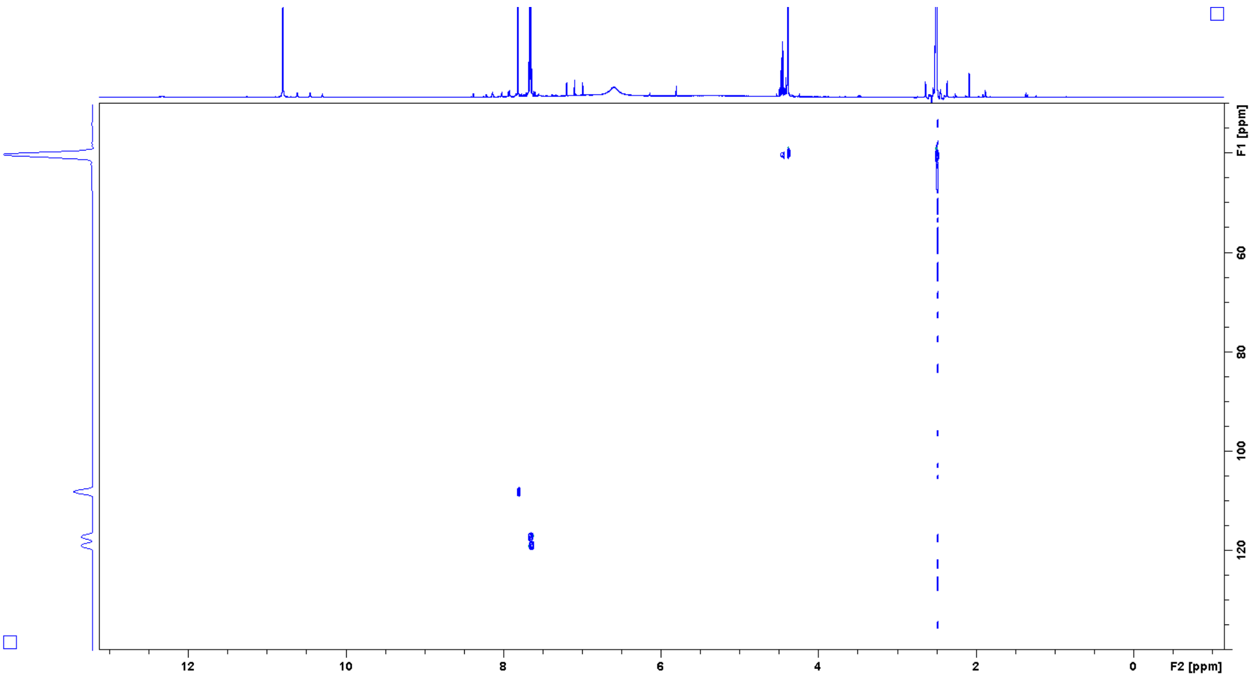

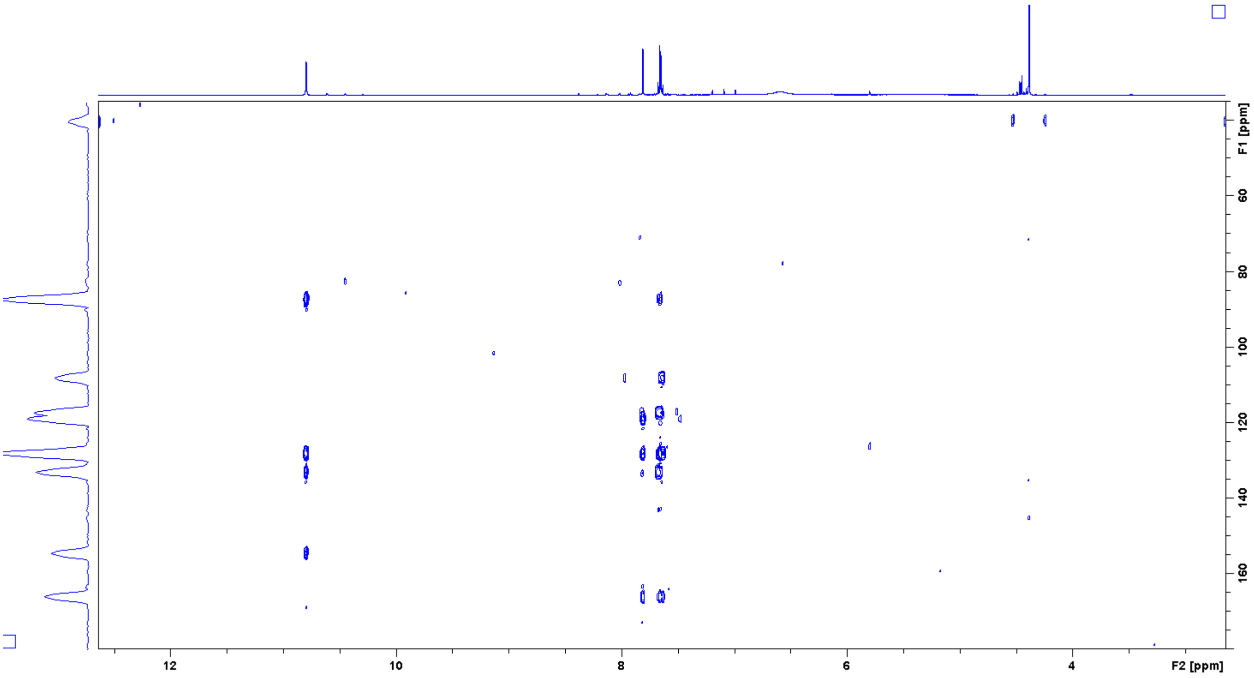


*2-Benzyl-7-(2-methyltetrazol-5-yl)-9H-pyrimido[4,5-b]indol-4-ol (**39**)*

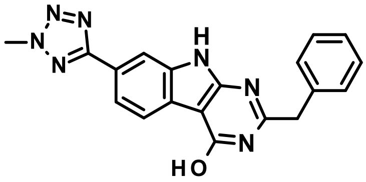


Title compound was synthesized according to the *General procedure D2* using 2-amino-6-(2-methyltetrazol-5-yl)-1H-indole-3-carboxamide (**38**, 200 mg, 0.77 mmol) to give the desired product (272 mg, 98%).

1H NMR (500 MHz, DMSO-$d_6$): δ ppm 12.48 (s, 1 H), 12.39 (s, 1 H), 8.08 (m, 1 H), 8.07 (m, 1 H), 7.92 (dd, J = 8.2, 1.4 Hz, 1 H), 7.39 (d, J = 7.3 Hz, 2 H), 7.34 (t, J = 7.5 Hz, 2 H), 7.26 (t, J = 7.3 Hz, 1 H), 4.43 (s, 3 H), 4.03 (s, 2 H); 13C NMR (125 MHz, DMSO-$d_6$): δ 165.3, 160.2, 159.4, 155.7, 136.9, 136.0, 129.6, 129.0, 127.3, 124.2, 122.7, 121.4, 119.8, 109.8, 98.6, 40.9, 40.1; UV/Vis: $\lambda_{max}$ 252 nm; HRMS (m/z): [M+H]+ calcd. for $C_{19}H_{15}N_7O$, 358.1416; found, 358.1411.

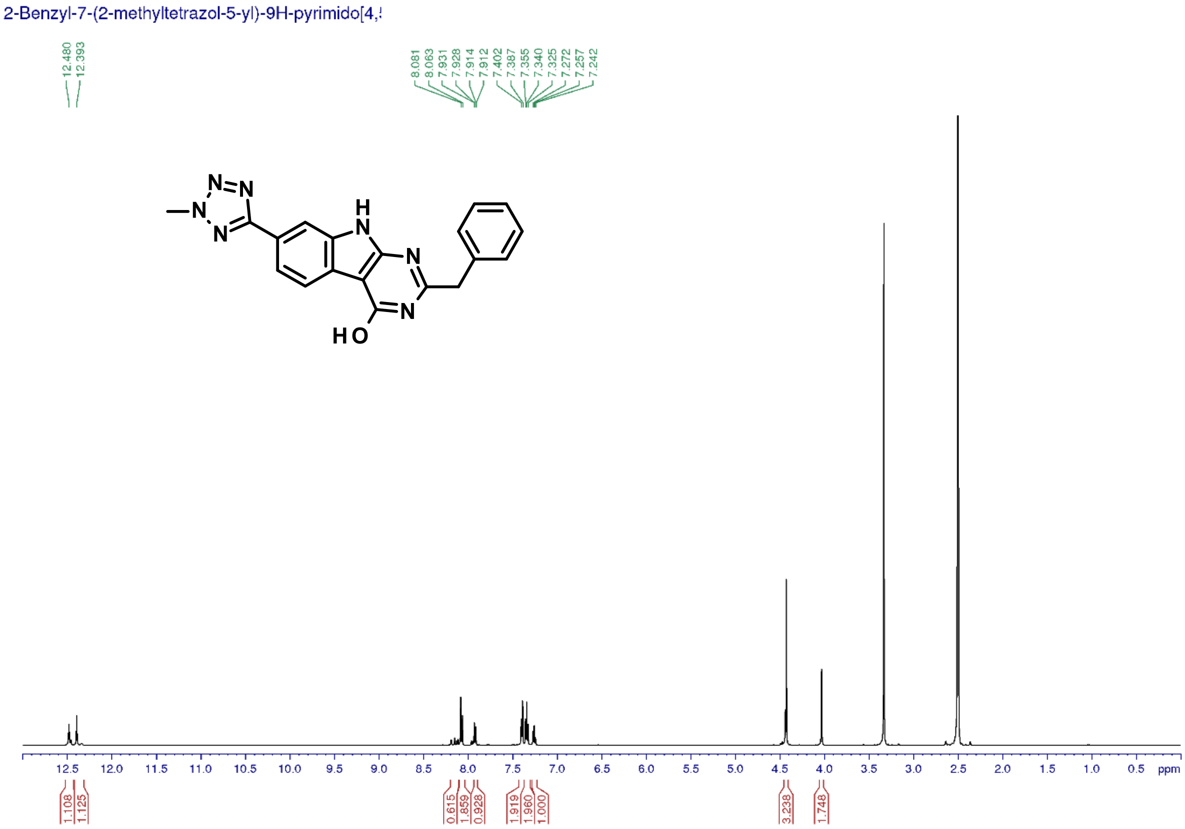


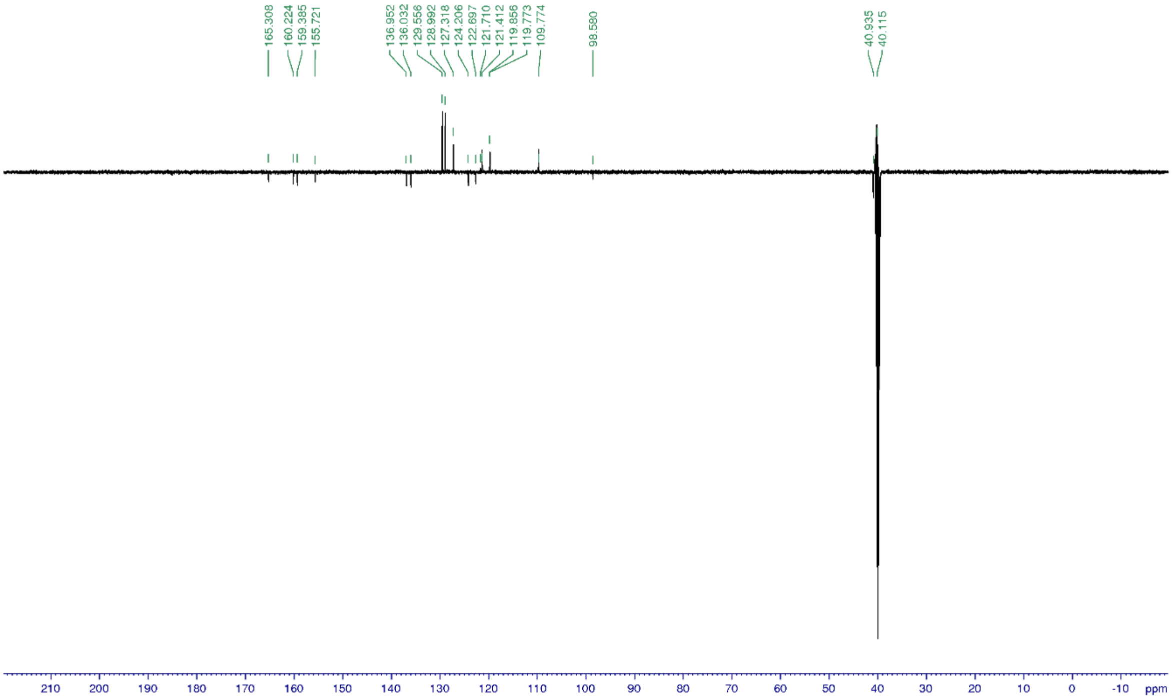


*2-Benzyl-4-chloro-7-(2-methyltetrazol-5-yl)-9H-pyrimido[4,5-b]indole*

***(40)***

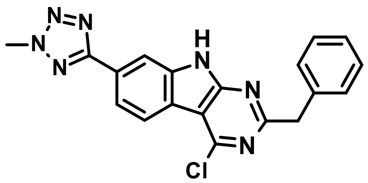

Title compound was synthesized according to the *General procedure G* using 2-benzyl-7-(2-methyltetrazol-5-yl)-9H-pyrimido[4,5-b]indol-4-ol (**39**, 13 mg, 0.04 mmol) to give the desired product (11 mg, 80%).

1H NMR (500 MHz, DMSO-d$_6$): δ ppm 12.96 (br s, 1 H), 8.38 (dd, J = 8.2, 0.6 Hz, 1 H), 8.23 (m, 1 H), 8.09 (dd, J = 8.1, 1.6 Hz, 1 H), 7.37 (m, 2 H), 7.33 (m, 2 H), 7.24 (m, 1 H), 4.47 (s, 3 H), 4.31 (s, 2 H); 13C NMR (125 MHz, DMSO-d$_6$): δ 166.6, 164.7, 157.9, 152.3, 139.8, 139.3, 129.7, 128.9, 126.9, 126.6, 123.5, 120.1, 119.9, 109.2, 45.3, 40.2; UV/Vis: $\lambda_{max}$ 245 nm; HRMS (m/z): [M+H]+ calcd. for $C_{19}H_{15}N_7O$, 358.1416; found, 358.1411.

2-Benzyl-4-chloro-7-(2-methyltetrazol-5-yl)-9H-pyrimido[4,5-b]indole

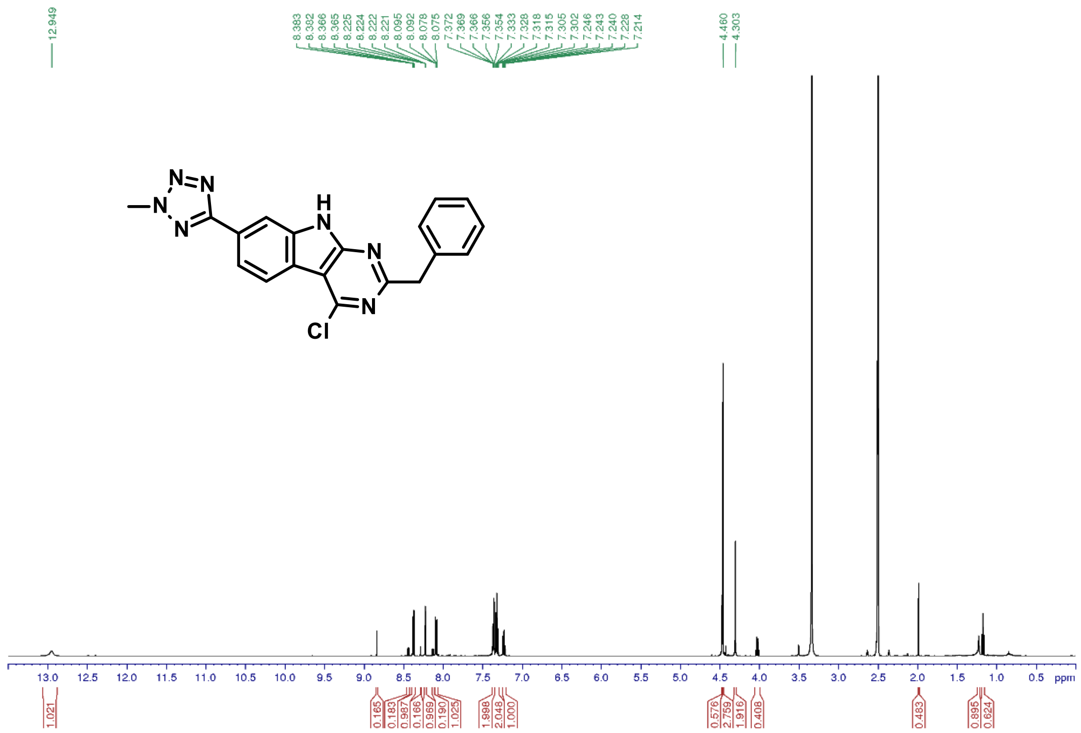

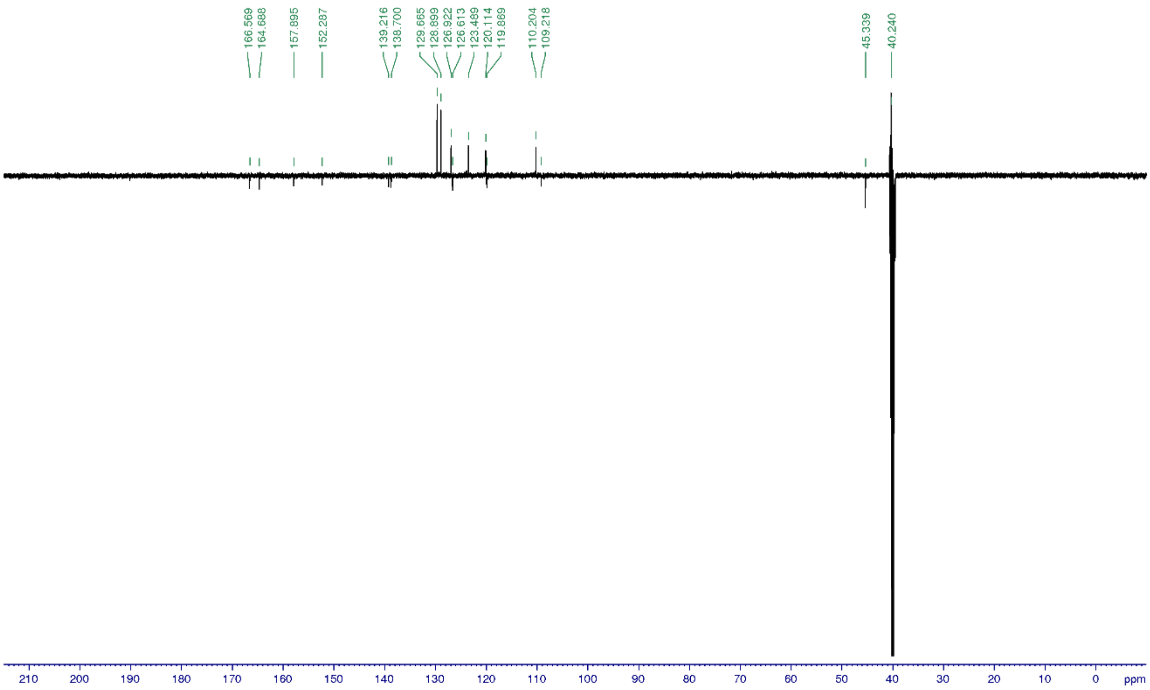


*7-(2-Methyltetrazol-5-yl)-9H-pyrimido[4,5-b]indol-4-ol (**41**)*

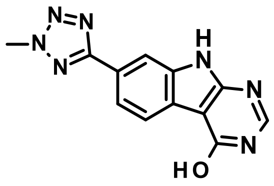

Title compound was synthesized according to the *General procedure D1* using 2-amino-6-(2-methyltetrazol-5-yl)-1H-indole-3-carboxamide (**38**, 230 mg, 0.90 mmol) to give the desired product (220 mg, 92%).

1H NMR (500 MHz, DMSO-$d_6$): δ ppm 12.33 (brs, 2 H), 8.18 (s, 1 H), 8.15 (d, J = 1.4 Hz, 1 H), 8.13 (d, J = 8.1 Hz, 1 H), 7.95 (dd, J = 8.2, 1.4 Hz, 1 H), 4.45 (s, 3 H); 13C NMR (125 MHz, DMSO-$d_6$): δ 165.4, 159.1, 155.3, 149.0, 136.0, 124.3, 122.9, 121.7, 119.8, 109.9, 100.7, 40.1; UV/Vis: $\lambda_{max}$ 248 nm; HRMS (m/z): [M+H]+ calcd. for $C_{12}H_9N_7O$, 268.0946; found, 268.0941.

7-(2-methyltetrazol-5-yl)-9H-pyrimido[4,5-b]indol-4-ol

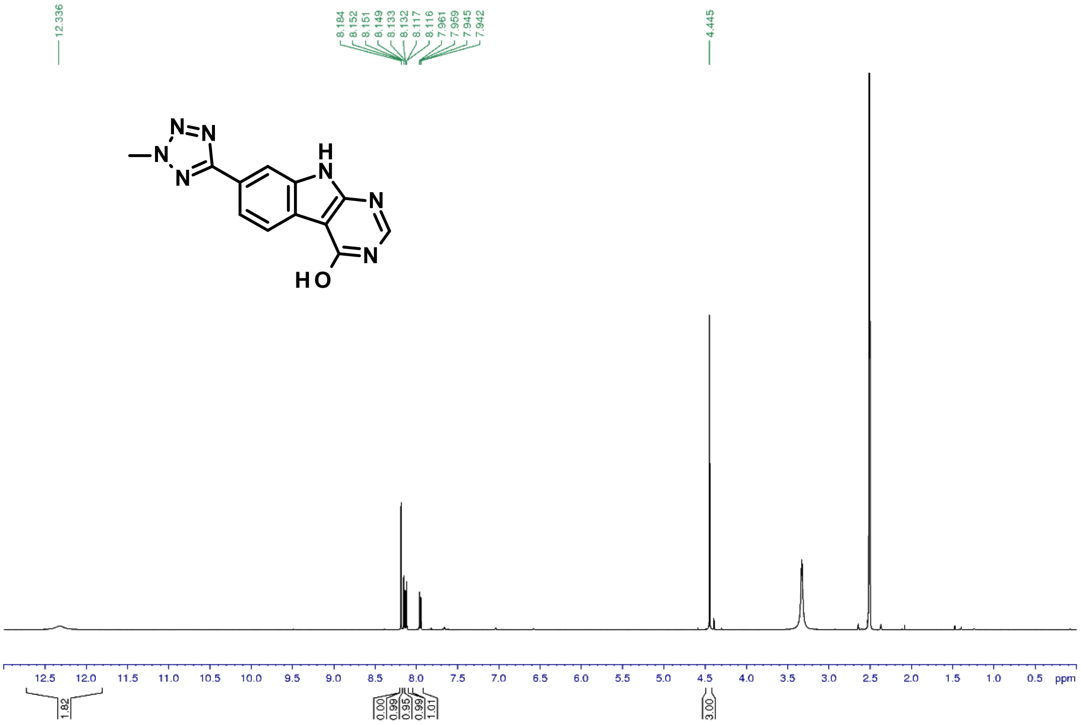


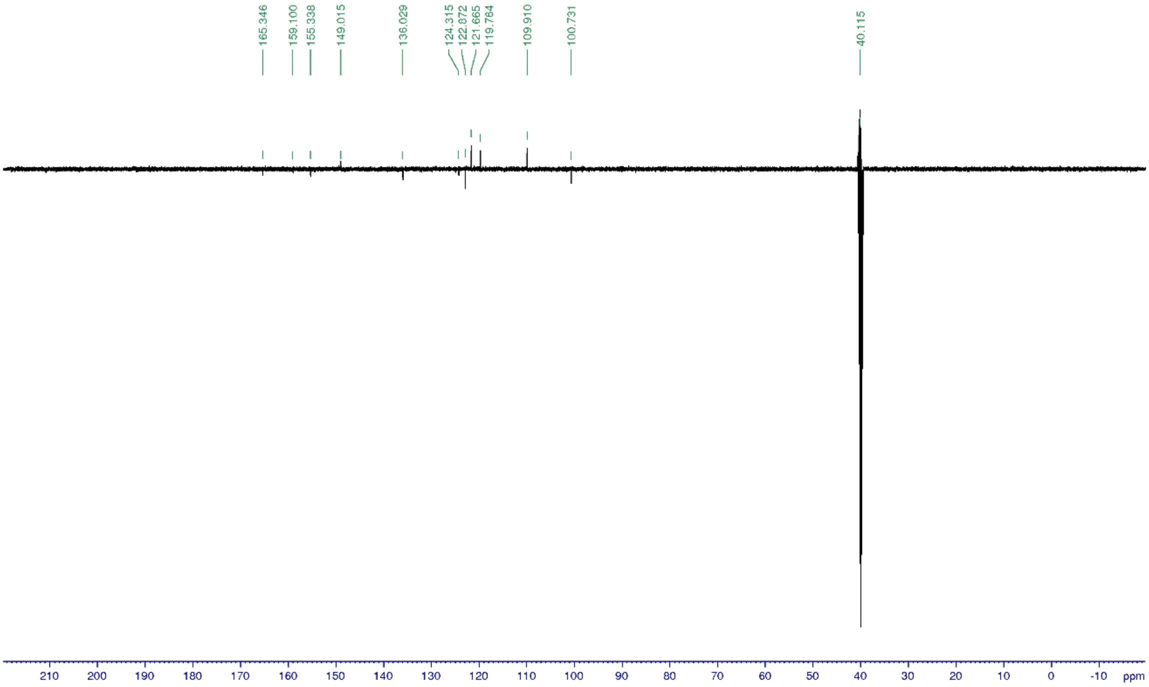


*4-Chloro-7-(2-methyltetrazol-5-yl)-9H-pyrimido[4,5-b]indole (**42**)*

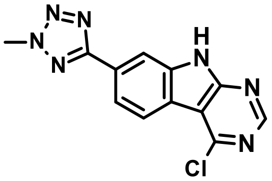

Title compound was synthesized according to the *General procedure G* using 7-(2-methyltetrazol-5-yl)-9H-pyrimido[4,5-b]indol-4-ol (**41**, 170 mg, 0.64 mmol) to give the desired product (150 mg, 83%).

1H NMR (500 MHz, DMSO-$d_6$): δ ppm 13.04 (brs, 1 H), 8.84 (s, 1 H), 8.44 (d, J = 8.4 Hz, 1 H), 8.28 (d, J = 1.4 Hz, 1 H), 8.13 (dd, J = 8.2, 1.3 Hz, 1 H), 4.48 (s, 3 H); 13C NMR (125 MHz, DMSO-$d_6$): δ 164.6, 157.0, 155.0, 152.3, 139.2, 127.0, 123.8, 120.2, 119.7, 111.4, 110.3, 40.0; UV/Vis: $\lambda_{max}$ 244 nm; HRMS (m/z): [M+H]+ calcd. for $C_{12}H_8N_7Cl$, 286.0607; found, 286.0605.

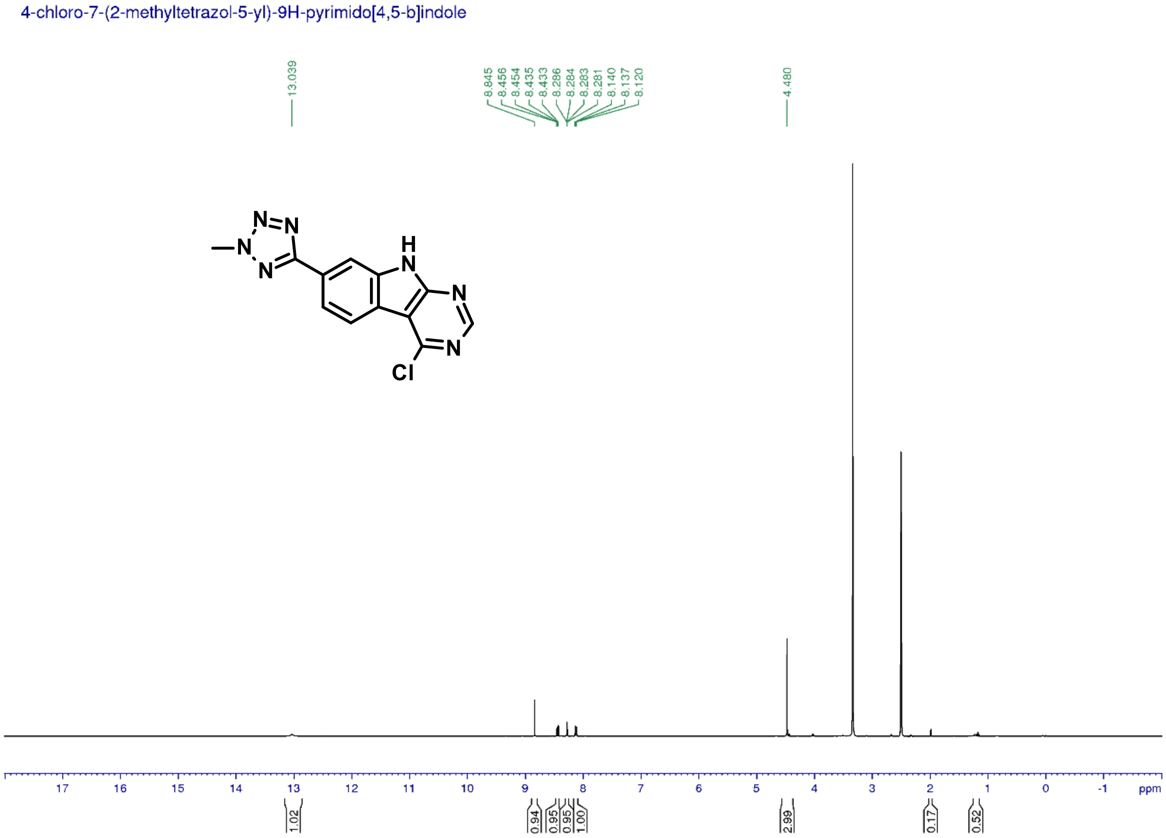

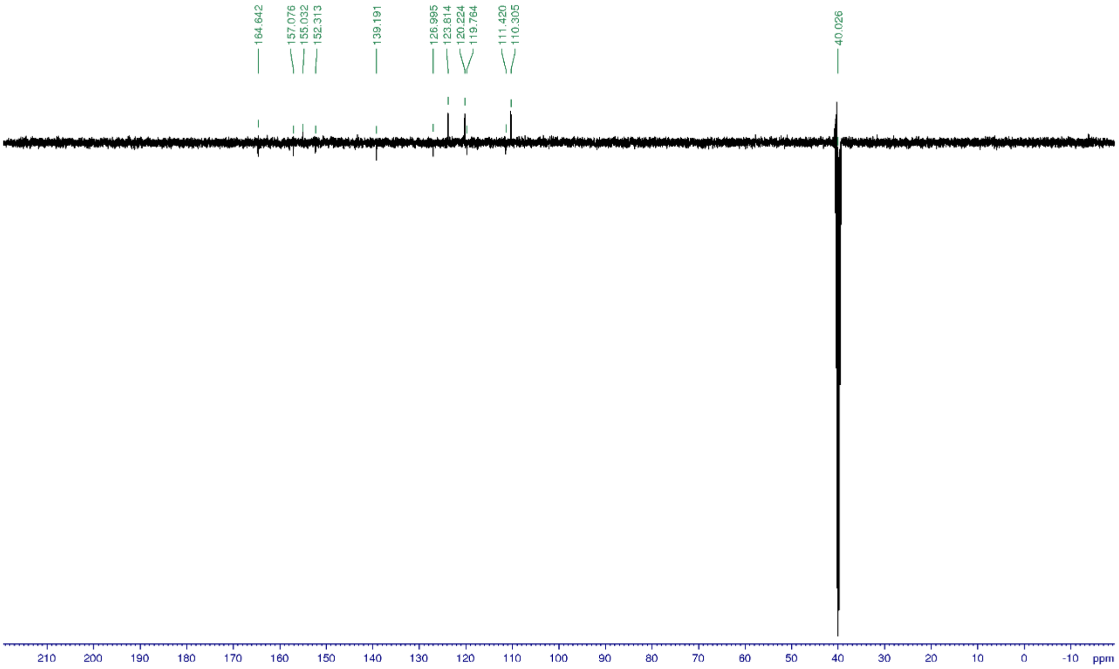


*3-(2-Methyltetrazol-5-yl)aniline (**43**)*

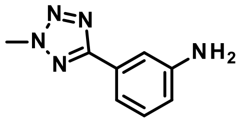

Title compound was synthesized according to the *General procedure K* using 2-methyl-5-(3-nitrophenyl)tetrazole (100 mg, 0.49 mmol) to give the desired product (79 mg, 93%).
1H NMR (500 MHz, DMSO-$d_6$): δ ppm 7.31 (m, 1H), 7.18-7.16 (m, 2H), 6.69 (m, 1 H), 5.38 (brs, 2 H), 4.40 (s, 3 H); 13C NMR (125 MHz, DMSO-$d_6$): δ 165.2, 149.8, 130.1, 127.9, 116.2, 114.0, 110.8, 40.0; UV/Vis: $\lambda_{max}$ 223 nm; HRMS (m/z): [M+H]+ calcd. for $C_8H_9N_5$, 176.0936; found, 176.0933.

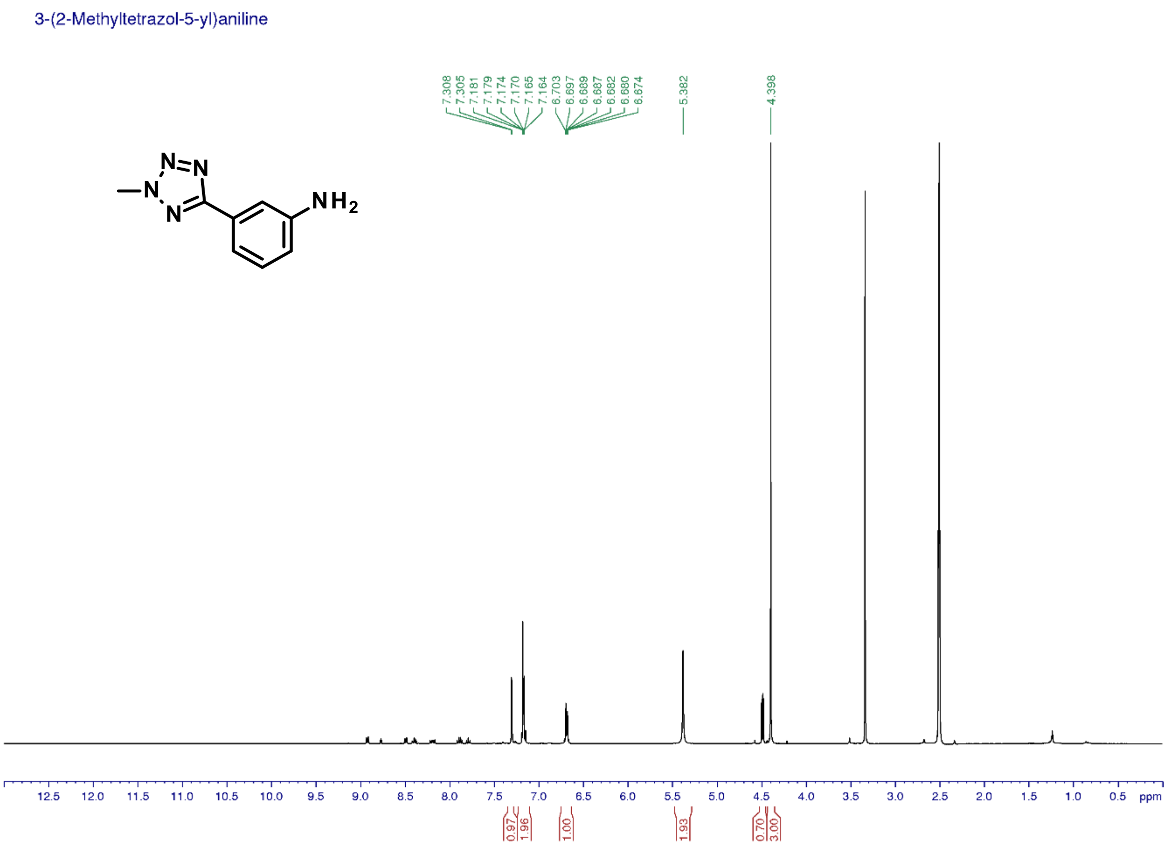


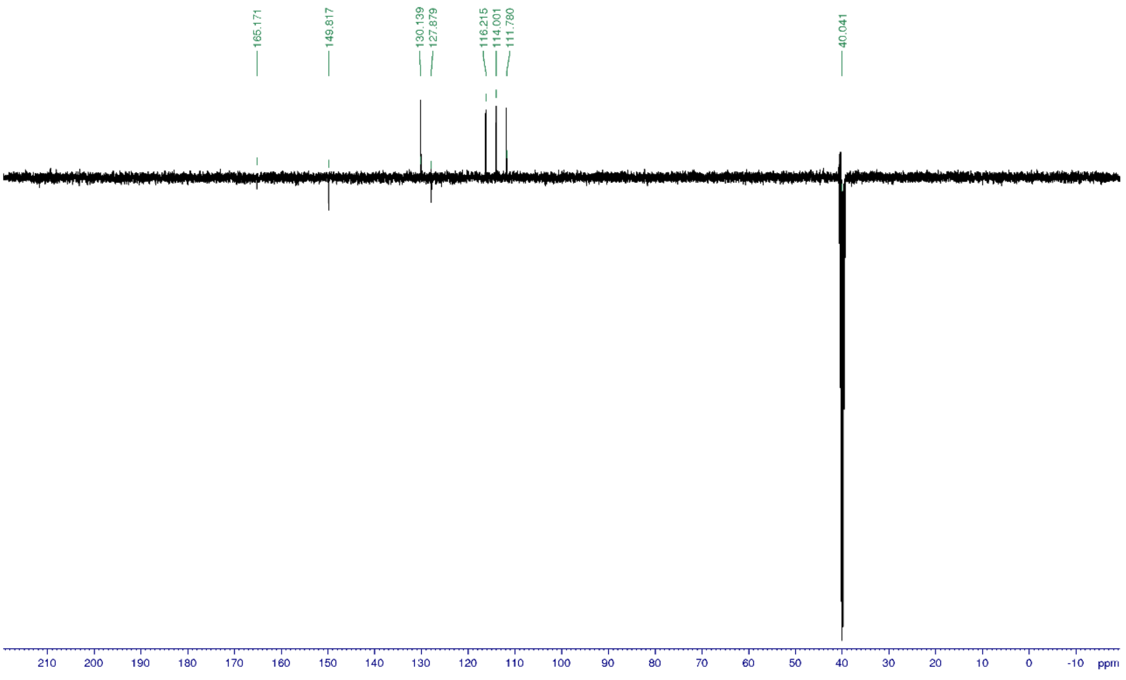

*2-Benzyl-6-chloro-N-[3-(2-methyltetrazol-5-yl)phenyl]pyrimidin-4-amine (**44**)*

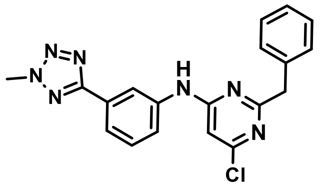


Title compound was synthesized according to the *General procedure L1* using 3-(2-methyltetrazol-5-yl)aniline (**43**, 20 mg, 0.11 mmol) to give the desired product (15 mg, 35%).

1H NMR (500 MHz, DMSO-$d_6$): δ ppm 10.13 (s, 1 H), 8.47 (s, 1 H), 7.76 (d, J = 8.1 Hz, 1 H), 7.71 (d, J = 7.7 Hz, 1 H), 7.46 (t, J = 7.9 Hz, 1 H), 7.43 (d, J = 76 Hz, 2 H), 7.31 (t, J = 6.7 Hz, 2 H), 7.23 (t, J = 7.2 Hz, 1 H), 6.71 (s, 1 H), 4.46 (s, 3 H), 4.05 (s, 2 H); 13C NMR (125 MHz, DMSO-$d_6$): δ 130.3, 129.8, 128.8, 126.9, 122.0, 120.9, 117.8, 103.4, 45.5, 40.2; UV/Vis: $\lambda_{max}$ 192 nm; HRMS (m/z): [M+H]+ calcd. for $C_{19}H_{16}ClN_7$, 378.1233; found, 378.1229.

2-Benzyl-6-chloro-N-[3-(2-methyltetrazol-5-yl)phenyl]pyrimidin-4-amine

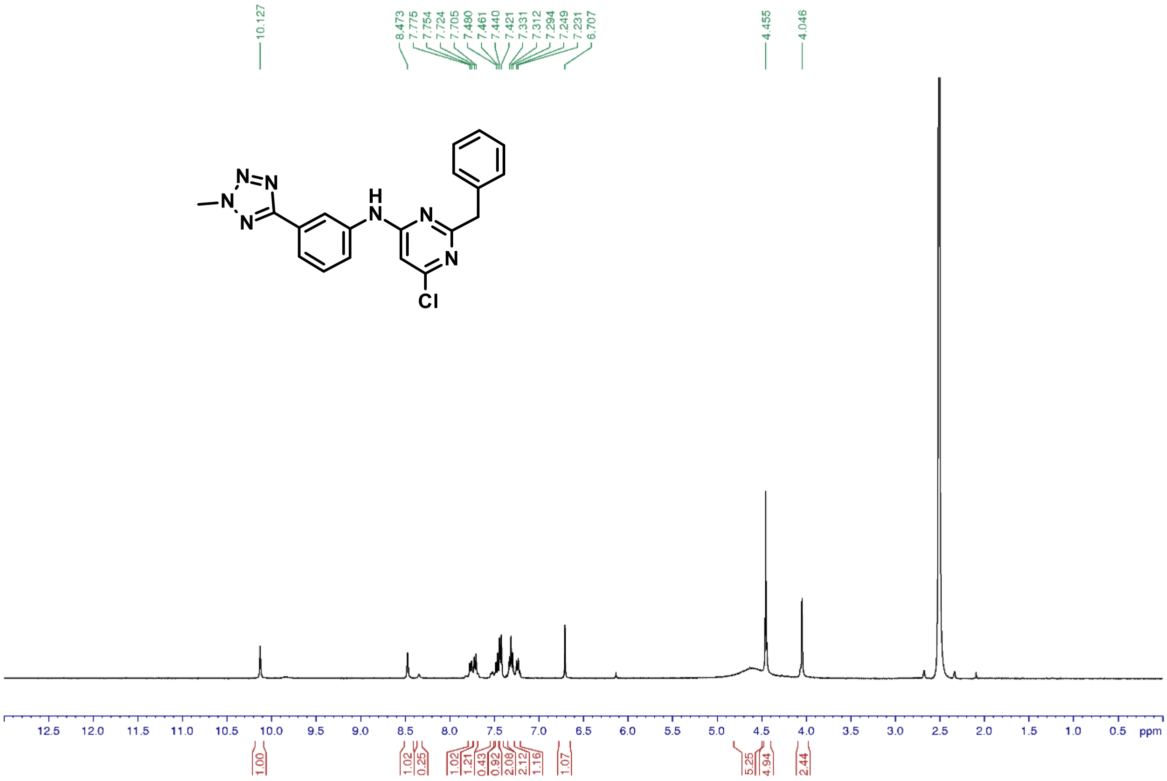

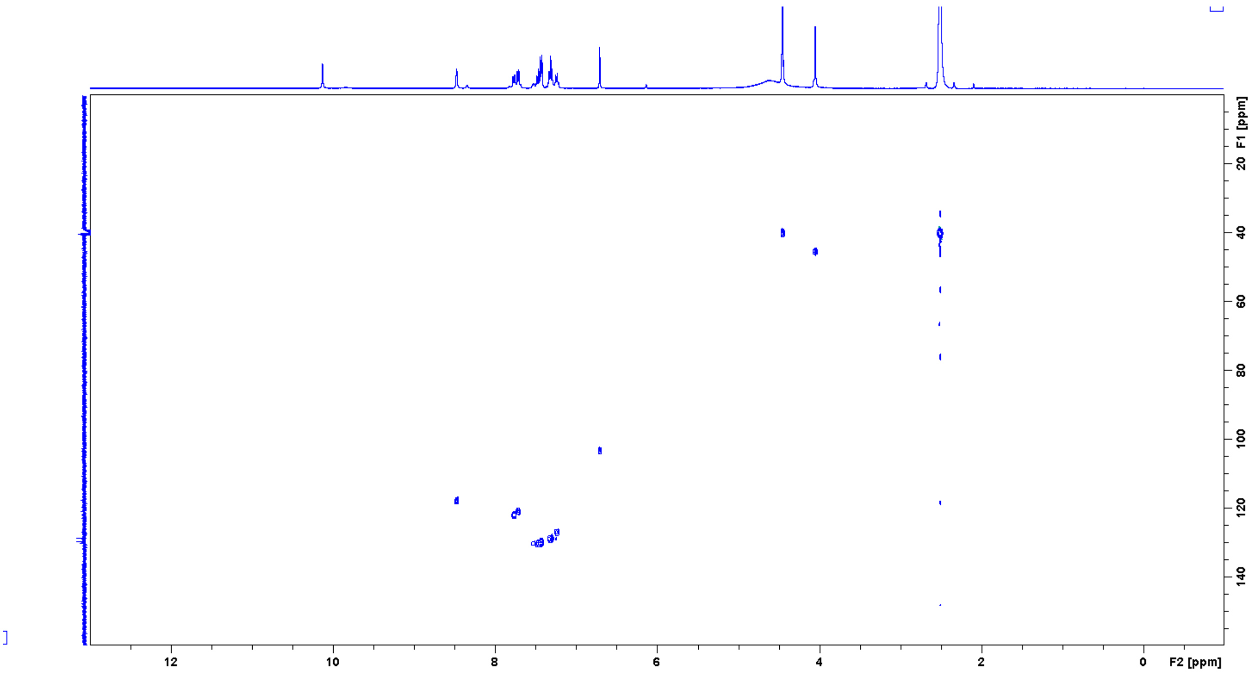


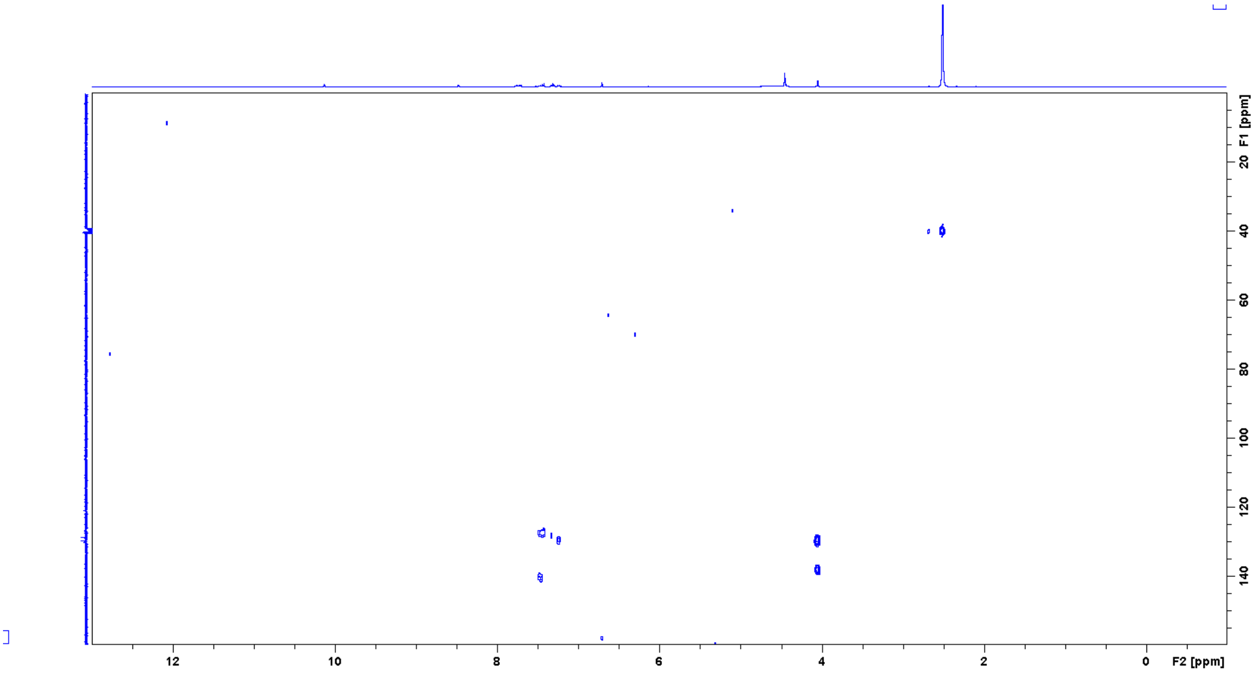


*2-Benzyl-4-chloro-9H-pyrimido[4,5-b]indole-7-carbonitrile (**45**)*

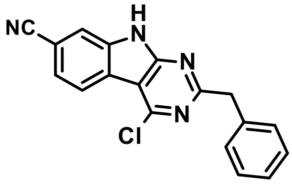


Title compound was synthesized according to the *General procedure G* using 2-benzyl-4-hydroxy-9H-pyrimido[4,5-b]indole-7-carbonitrile (**26**, 50 mg, 0.16 mmol) to give the desired product (39 mg, 73%).

1H NMR (400 MHz, DMSO-$d_6$): δ ppm 13.14 (br., 1 H), 8.38 (dd, J = 8.2, 0.7 Hz, 1 H), 8.06 (dd, J = 1.4, 0.7 Hz, 1 H), 7.79 (dd, J = 8.2, 1.5 Hz, 1 H), 7.35 (d, J = 6.8 Hz, 2 H), 7.31 (t, J =

7.3 Hz, 2 H), 7.23 (t, J = 7.2 Hz, 1 H), 4.31 (s, 2 H); 13C NMR (100 MHz, DMSO-$d_6$): δ 129.6, 129.1, 127.0, 125.2, 123.5, 116.9, 45.4; UV/Vis: $\lambda_{max}$ 249 nm; HRMS (m/z): [M+H]+ calcd. for $C_{18}H_{11}ClN_4$, 319.0750; found, 319.0746.

2-Benzyl-4-chloro-9H-pyrimido[4,5-b]indole-7-carbonitrile

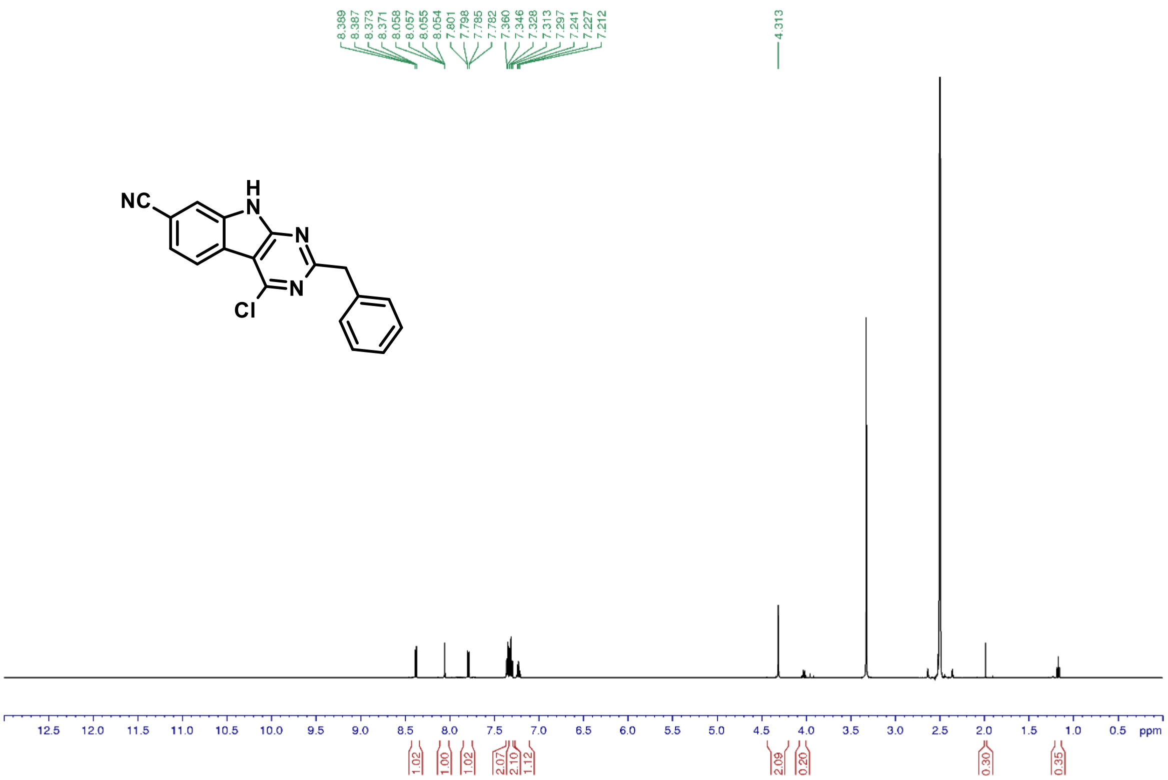


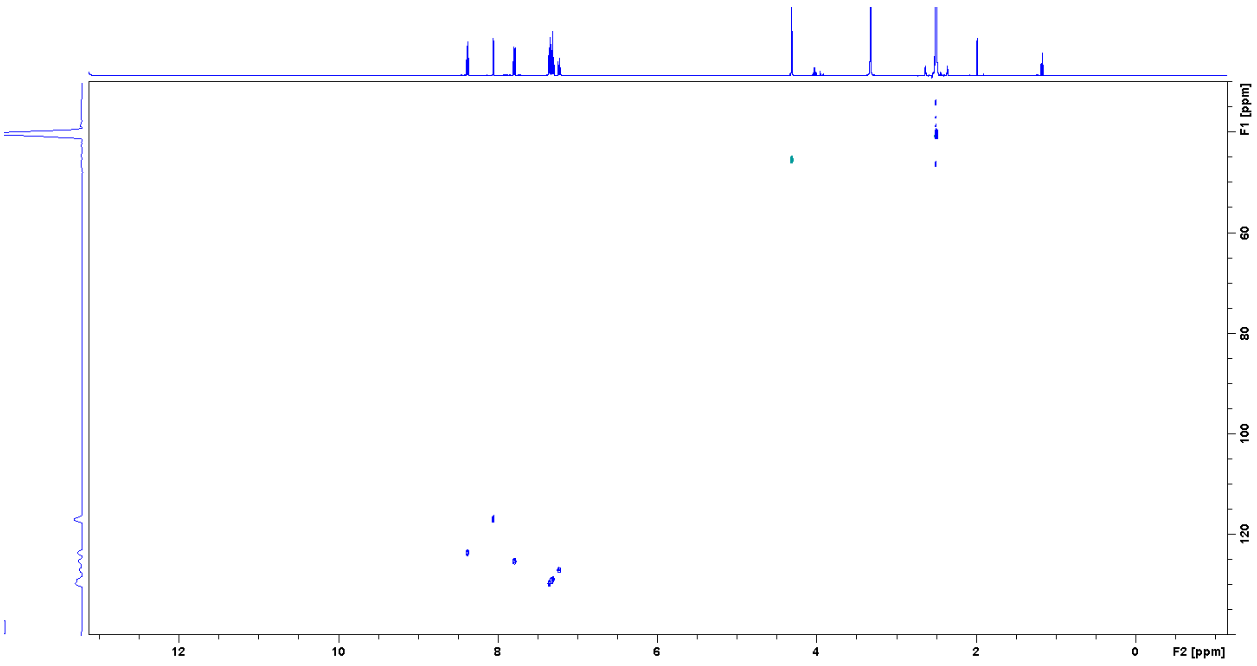

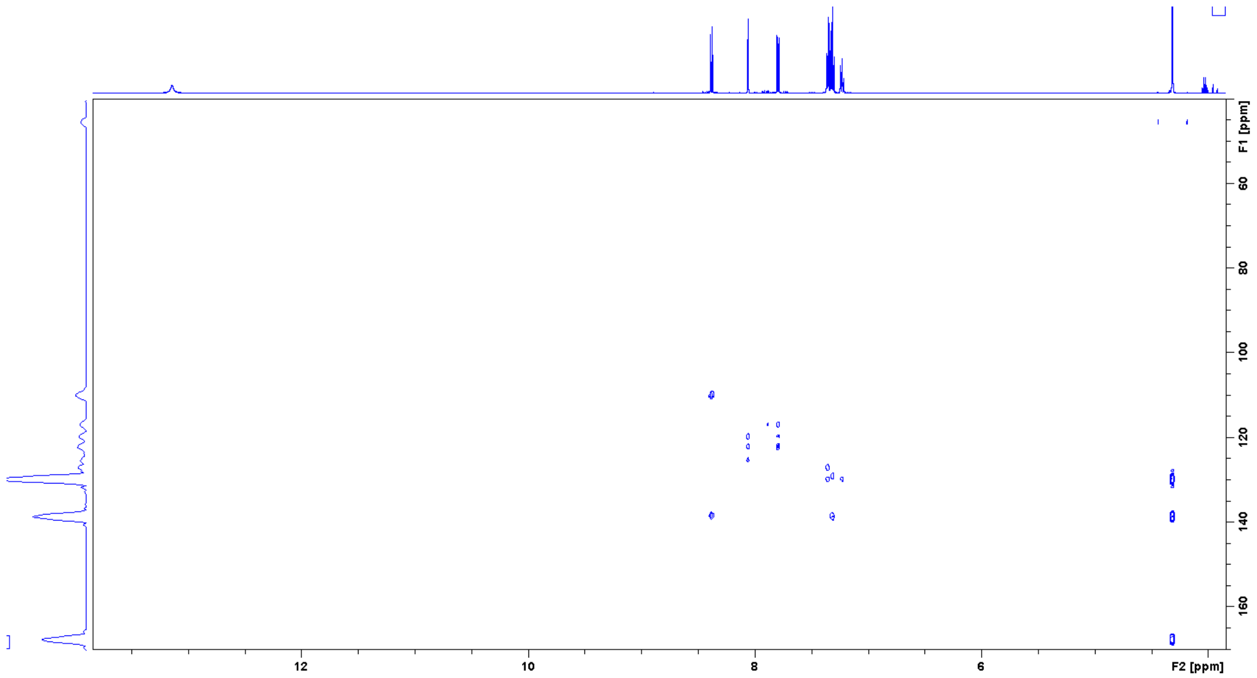

F1 [ppm]
60
80
100
120
140
160
12
10
8
6
F2 [ppm]
